# Recovery of complete genomes of canine parvovirus from clinical samples

**DOI:** 10.1101/2023.07.12.548703

**Authors:** Sara França de Araújo dos Santos, Ueric José Borges de Souza, Martha Trindade Oliveira, Jairo Jaime, Fernando Rosado Spilki, Ana Cláudia Franco, Paulo Michel Roehe, Fabrício Souza Campos

## Abstract

Canine parvovirus (CPV) is a highly pathogenic virus that affects dogs, especially puppies. CPV is believed to have evolved from feline panleukopenia virus (FPV), eventually giving rise to three antigenic types, CPV-2a, 2b, and 2c. CPV-2 is recognized for its resilience in contaminated environments, ease of transmission among dogs, and pathogenicity for puppies. Despite the relevance of the virus, complete genome sequences of CPV available at GenBank, to date, are scarce. In the current study, we have developed a methodology to allow the recovery of complete CPV-2 genomes directly from clinical samples. For this, seven fecal samples from Gurupi, Tocantins, North Brazil, were collected from puppies with clinical signals of viral enteritis, and submitted to viral DNA isolation and amplification. Two multiplex PCR strategies were designed including primers targeting fragments of 400 base pairs (bp) and 1,000 bp along the complete genome. Sequencing was performed with the Nanopore^®^ technology and results obtained with the two approaches were compared. Genome assembly revealed that the 400 bp amplicons generated larger numbers of reads, allowing a more reliable coverage of the whole genome than those attained with primers targeting the larger (1000 bp) amplicons. Nevertheless, both enrichment methodologies were efficient in amplification and sequencing. Viral genome sequences were of high quality and allowed more precise typing and subtyping of viral genomes compared to the commonly employed strategy relying solely on the analysis of the VP2 region, which is limited in scope. The CPV-2 genomes recovered in this study belong to the CPV2a and CPV-2c subtypes, closely related to isolates from the neighboring Amazonian region. In conclusion, the technique reported here may contribute to increase the number of full CPV genomes available, which is essential for understanding the genetic mechanisms underlying the evolution and spread of CPV-2.

## 1. Introduction

Canine parvovirus (CPV, *Protoparvovirus carnivoran 1;* ICTV, 2023) is the agent of a highly contagious viral disease that affects dogs, especially puppies (Carmichael and Binn, 1981; Gumbrell, 1979; Kelly and Atwell, 1979). The virus can be transmitted through direct contact with an infected dog or indirectly through contaminated objects such as food and water bowls, leashes, and even the shoes of people who have come into contact with infected dogs or their excreta (Nandi and Kumar, 2010).

CPV is believed to have evolved from the feline panleukopenia virus (FPV), which can infect cats and dogs (Kelly, 1978). FPV and CPV are very similar in their genetic makeup and structure, and both viruses bind to the same host cell receptor in target cells (Ikeda et al., 2002). However, one notable mutation that has been observed in CPV is the change of an amino acid residue from aspartic acid (D) to glutamic acid (E) at position 426 of the VP2 protein (Allison et al., 2015; Parrish et al., 1985; Zhou et al., 2017). This mutation, known as the CPV-2a mutation, is thought to have arisen due to the virus’s adaptation to the new host (dogs) and its attempts to evade the host’s immune system (Decaro and Buonavoglia, 2012). The CPV-2a mutation, associated with higher levels of virulence and mortality in infected dogs, played a crucial role in the emergence of a novel CPV strain referred to as CPV-2b. A new subtype, named CPV-2b, was reported in the mid-1980s (Hong et al., 2007; Ikeda et al., 2002; Martella et al., 2004). This variant has also been linked to increased virulence and evasion from vaccine-induced immune response. In addition, a third variant known as CPV-2c, also containing the D426E substitution, was reported in Italy in 2000 (Buonavoglia et al., 2001). CPV-2c has similarly been associated with increased virulence and resistance to vaccination. Together, the emergence of CPV-2b and CPV-2c highlights the ongoing evolution and diversification of CPV strains, emphasizing the need for continued research on the genetic characteristics and epidemiology of these subtypes (Hoelzer and Parrish, 2010; Truyen, 2006). Based on these amino acid changes in the VP2, a traditional system of classification of CPV-2 was created (Decaro et al., 2009; Martella et al., 2004; Parrish et al., 1991). However, these changes are not sufficient to classify all sequences available in GenBank to date into subtypes, and a new classification system based on genogroups was recently proposed (de Oliveira Santana et al., 2022).

In 1988, Reed et al. defined the CPV’s nucleotide sequence and genome organization of CPV. However, sequencing the 5’ and 3’ ends, which harbor hairpin structures, has proven difficult. This remains to date, in such a way that the majority of the CPV genomes available at GenBank are not complete (currently, 256 sequences with more than 4000 bp and only 71 sequences with more than 5000 bp are available in the database). So, despite the small size of the genome and the virus’ importance for puppies, the genomic information remains scarce. In order to fulfill this gap, in attempting to facilitate recovery of complete CPV genomes, we have adapted the multiplex PCR methodology to provide a more comprehensive sequencing approach using two sets of primers: the first was designed to amplify short fragments (400 bp), and the second designed to amplify longer (1000 bp) fragments along the complete CPV genome. The two were compared to evaluate the best strategy to provide full-length CPV genomes.

## 2. Materials and methods

### 2.1 Samples

Rectal swab samples were collected from animals tested with the Rapid CPV Antigen Test Kit (Alere®, Bioesay Inc., Korea) between November 2022 and January 2023, at a veterinary clinic in the city of Gurupi, Tocantins, North Brazil (Table 1). Samples were placed in sterile microtubes (2 mL) and frozen at −20°C until processing. Nucleic acid extraction was carried out at the Molecular Biology Laboratory of the Federal University of Tocantins, as described below.

**Table 1.** – Data from animals.

| Animal | Age (months) | Sex | Breed | Collection data | Neighborhood | CPV antigen test | Vaccine status | Outcome |
| --- | --- | --- | --- | --- | --- | --- | --- | --- |
| 1 | 4 | Female | Mixed | 12/05/22 | Jardim São Lucas | Positive | Incomplete | Recovery |
| 2 | 7 | Male | Rottweiler | 12/01/22 | Loteamento Águas Claras | Positive | Incomplete | Recovery |
| 3 | 3 | Female | Dachshund | 12/30/22 | Vila Independência | Positive | Complete | Death |
| 4 | 8 | Male | Pinscher | 12/29/22 | Parque das Acácias | Positive | Incomplete | Recovery |
| 5 | 5 | Female | Blue Heeler | 12/01/22 | Zona Rural de Gurupi | Positive | Incomplete | Death |
| 6 | 4 | Female | Mixed | 11/02/22 | Centro | Negative | Incomplete | Death |
| 7 | 6 | Male | Pinscher | 01/18/23 | Jardim Guanabara | Negative | Incomplete | Death |

### 2.2 Nucleic acid extraction

Dry swabs of fecal samples were thawed and suspended in 400 μL of buffer from the kit described below. Nucleic acid extraction and purification were performed using the Quick-DNA/RNA™ Viral MagBead (Zymo Research, Irvine, California, USA) following the manufacturer’s recommendations, in the automated extractor (Loccus Extracta 32, São Paulo, Brazil). Briefly, the 400 μL of viral DNA/RNA buffer of sample suspension was mixed with 10 μL of beads, and 4 μL of proteinase K. Subsequently, three washing steps were executed. The first step utilized 250 µL of MagBead DNA/RNA Wash 1, the second step employed 250 µL of MagBead DNA/RNA Wash 2, and finally, a last wash was performed using 250 µL of buffer with 80% ethanol. In the final step, the samples were eluted in 50 µL of DNase/RNase-free water.

### 2.3 Multiplex PCR to obtain CPV complete genome

Two sets of primers were designed aiming amplification of 400-bp-long and 1000-bp-long genome fragments using the primal scheme tool available at https://primalscheme.com/ (Quick et al., 2017). Primers were constructed using the CPV reference sequence available at GenBank (accession number NC_001539.1), with the default designing scheme settings. Additionally, all complete or nearly complete genomes available at GenBank (a total of 250 CPV genomes, Supplementary Table S1) were aligned using the Mafft alignment and primer design tools from Geneious Prime version 2023.0 with default settings to add primers at the 5’ and 3’ ends of genomes. For multiplex PCR, two sets of primers (named Pool 1 and Pool 2) for each of the targeted fragment sizes (400 and 1000 bp). The designed primers were acquired commercially (Exxtend, São Paulo, Brazil) (Tables 2 and 3).

**Table 2.**
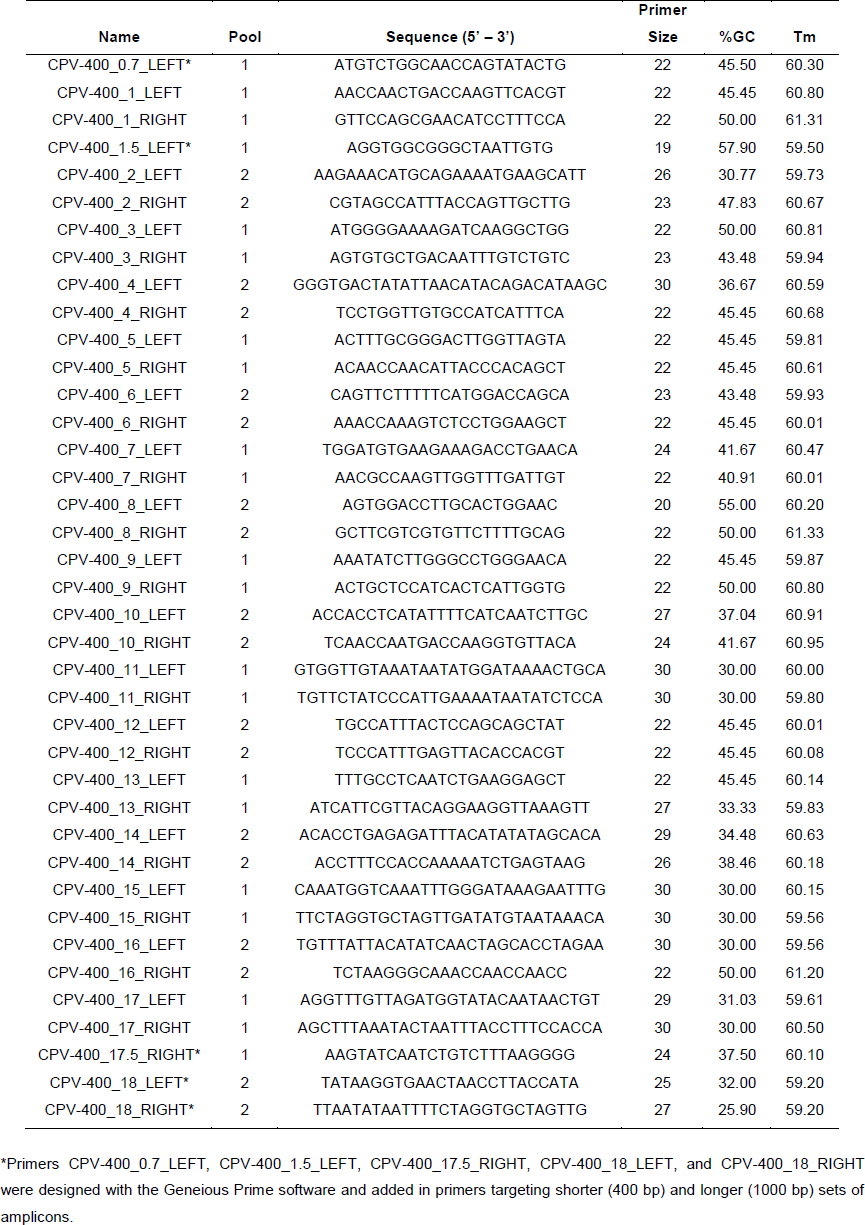
Primers targeting 400 bp amplicons employed in this study.

**Table 3.**
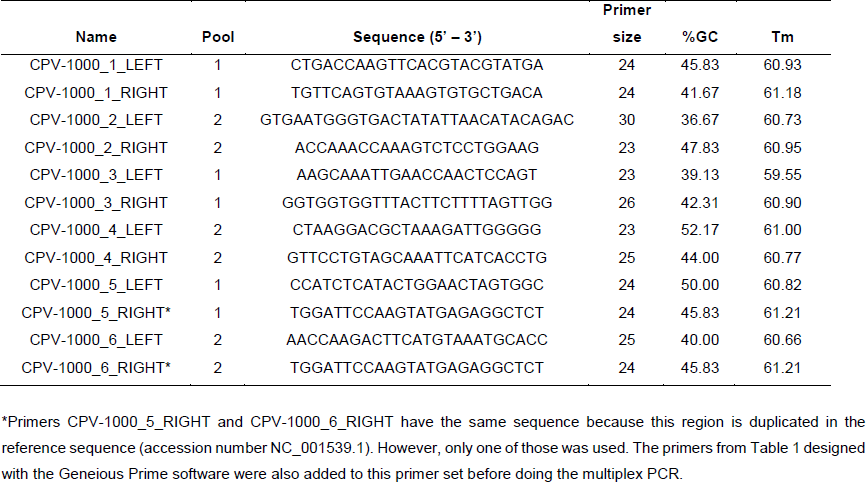
Primers targeting 1000 bp amplicons employed in this study.

DNA was amplified by Q5® High-Fidelity 2X Master Mix (New England Biolabs) in a 25 μl reaction. For primers targeting 400 bp amplicons, the two reactions included: 12.5 μl 2x master mix, 5 μl DNA, 5.9 μl nuclease-free H_2_O, and 1.6 μl of a primer pool at a final concentration of 10 nanomolar (nM). For primers targeting the 1000 bp amplicons, components were: 12.5 μl 2x master mix, 5 μl DNA, 7.1 μl nuclease-free H_2_O, and 0.4 μl of a primer pool at a final 10 nM concentration. For both sets of primers, the PCR temperature and timing conditions were: 98°C for 3 min, 35 cycles of 98°C for 15 s, and 63°C for 5 min, and held at 4°C.

### 2.4 Sequencing and genome assembly

The amplicons obtained with each of the strategies (using pools 1 and 2) were pooled in each group and purified with one volume of AMPure XP beads (Beckman Coulter, Brea, CA, USA). The beads were washed with 80% ethanol, and the clean PCR products’ concentrations were measured using a Qubit dsDNA HS Assay Kit on a Qubit 3.0 fluorometer (ThermoFisher Scientific Corporation, Waltham, MA, USA). The MinION library preparation was performed using a Ligation Sequencing kit SQK-LSK-109 and Native Barcoding kits EXP-NBD104 and EXP-NBD114 (Oxford Nanopore, Oxford, UK). The resulting library was loaded on a R9.4 Oxford MinION flowcell (FLO-MIN106) and sequenced using a MinION Mk1B device. The ONT MinKNOW software was used to collect raw data. High-accuracy base-calling of raw FAST5 files and barcode demultiplexing were performed using Guppy (v6.0.1).

Reads were trimmed using Geneious Prime version 2023.1.1 with an error probability limit of 0.05 (trim regions with more than a 5% chance of no error per base). Assembly and annotation of the viral genomes were done using template-assisted assembly on the CPV reference genome (GenBank accession number NC_001539.1) using the map to reference tools of Geneious Prime version 2023.1.1 with the following settings: medium sensitivity, fast time, and fine-tuning of iterates up to 5 times. After assembly, the seven CPV genome sequences reported in this study were generated to fullness. The resulting genomes obtained with either the shorter (400 bp) or the longer (1000 bp) amplicons were mounted separately, totalizing fourteen new, full-length CPV genomes. These were compared in the number of reads and the degrees of genome coverage achieved.

### 2.5 Phylogenetic analyses

To investigate the phylogenetic relationships among CPV genomes, the newly generated sequences were analyzed together with 305 CPV-2 genomes (selected based on the length of the sequenced fragment, which should be equal to or greater than 4000 bases; Supplementary Table S2) and 692 VP2 gene sequences (Supplementary Table S3), both obtained from GenBank (http://www.ncbi.nlm.nih.gov/ accessed on May 5, 2023). Sequences were aligned using MAFFT v7.490 (Katoh and Standley, 2013) with default settings, and manually inspected using AliView v1.28 (Larsson, 2014). The Maximum Likelihood (ML) analyses were performed under the TPM3u model of nucleotide substitution with the empirical base frequencies (+F) plus FreeRate model (+R2), as selected by the ModelFinder software (TPM3u+F+R2) (Kalyaanamoorthy et al., 2017) and 1000 replicates of ultrafast bootstrapping (−B 1000) and an SH-aLRT branch test (−alrt 1000). Tree visualization was performed using FigTree v1.4.4 (Rambaut, 2023).

To infer a time-scaled phylogeny, the ML tree of partial full-genomes was inspected in TempEst v1.5.3 (Rambaut et al., 2016) to investigate the temporal signal through a regression analysis of root-to-tip genetic distance against sampling dates. Outlier sequences were removed from the analysis, leaving in the final dataset 185 sequences (Supplementary Table S4). The phylogeny was reconstructed using Bayesian inference with Markov Chain Monte Carlo (MCMC) sampling, as implemented in the BEAST v1.10.4 package (Suchard et al., 2018). A stringent model selection analysis using both path-sampling (PS) and stepping stone (SS) procedures were employed to estimate the most appropriate molecular clock model for the Bayesian phylogenetic analysis by running 100 path steps of 1 million interactions each (Baele et al., 2012). For that, we used the strict molecular clock model and the more flexible uncorrelated log-normal relaxed molecular clock model with two non-parametric population growth models: (i) Bayesian skygrid coalescent model and (ii) the standard Bayesian skyline plot (BSP; 10 groups) (Drummond et al., 2005; Drummond et al., 2006; Gill et al., 2013). Both SS and PS estimators indicated the Bayesian skyline plot with an uncorrelated log-normal relaxed molecular clock as the best-fitted model for the dataset under analysis (Supplementary Table S5). Triplicate MCMC runs of 100 million states each were then computed, with sampling every 10000 steps to ensure stationary and adequate effective sample size (ESS) for all statistical parameters. The three independent runs were merged with Log combiner V1.10.4 and the convergence of the MCMC chain was assessed using Tracer v1.7.2 (Rambaut et al., 2018). The TreeAnnotator V1.10.4 was used to summarize the maximum clade trees from the MCMC samples with 10% removed as burn-in and the MCMC phylogenetic tree was visualized using the ggtree R package (Yu et al., 2017).

## 3. Results

The number of reads and contigs generated with the two sets of primers targeting smaller (400 bp) and larger (1000 bp) amplicons, after sequencing and genome assembly, are shown in Table 4. Also shown are the mean coverages and the resulting final genome sizes after assembly.

**Table 4.** - Number of reads and contigs generated with the two sets of primers targeting smaller (400 bp) and larger (1000 bp) amplicons along the CPV genome.

| Sample | Barcode* | Product<br>(bp) | Total<br>reads | Used<br>reads | Mean<br>coverage | Maximum<br>Coverage | Contig<br>size | Genome<br>size |
| --- | --- | --- | --- | --- | --- | --- | --- | --- |
| 1 | 1 | 400 | 24,001 | 22,914 | 511.8 | 1,965.0 | 6,300 | 5,351 |
| 2 | 9 | 400 | 28,894 | 28,030 | 529.3 | 2,624.0 | 6,166 | 5,376 |
| 3 | 17 | 400 | 31,238 | 30,014 | 616.4 | 2,627.0 | 7,072 | 5,369 |
| 4 | 25 | 400 | 36,317 | 34,910 | 742.7 | 3,276.0 | 6,843 | 5,370 |
| 5 | 33 | 400 | 24,729 | 24,061 | 532.6 | 2,579.0 | 5,919 | 5,376 |
| 6 | 41 | 400 | 26,418 | 25,474 | 496.0 | 2,326.0 | 7,896 | 5,367 |
| 7 | 49 | 400 | 27,966 | 26,997 | 544.7 | 2,652.0 | 7,103 | 5,376 |
| 1 | 65 | 1000 | 3,105 | 2,985 | 95.1 | 313.0 | 5,735 | 5,362 |
| 2 | 73 | 1000 | 3,551 | 3,427 | 83.9 | 357.0 | 7,977 | 5,348 |
| 3 | 81 | 1000 | 2,578 | 2,399 | 84.3 | 254.0 | 5,646 | 5,370 |
| 4 | 89 | 1000 | 6,171 | 5,999 | 206.8 | 680.0 | 5,984 | 5,358 |
| 5 | 2 | 1000 | 4,816 | 4,592 | 129.1 | 581.0 | 7,474 | 5,370 |
| 6 | 10 | 1000 | 4,503 | 4,369 | 154.9 | 485.0 | 5,708 | 5,370 |
| 7 | 18 | 1000 | 6,757 | 6,476 | 216.4 | 836.0 | 6,330 | 5,368 |

\*Barcode used for nanopore sequencing.

All genomes were assembled to completeness, using the size of the NCBI reference sequence NC_001539.1 as a reference, which is equal to or greater than 5,323 bp. However, it is worth noting that the region from nucleotide 5,068 onwards exhibits repetition (5,069-5,319 nt) due to the presence of a hairpin structure at the 3’ end of the genome. Additionally, the coverage of the assembled genomes using 400-bp-long fragments exhibited higher read counts and coverage, as compared to the genomes assembled using the 1000-bp-long fragments. Nonetheless, both enrichment methodologies proved valuable in achieving sufficient depth of coverage for full genome assembly. The seven CPV genomes generated were deposited in GenBank with the accession numbers: OR230510-OR230516.

### 3.1 Phylogenetic Analysis

With the sequences generated in this study, in addition to the CPV sequences currently available at GenBank as described above, we conducted a comprehensive analysis to compare the phylogenetic clustering of CPV sequences. While this analysis is typically based on the sequence of the VP2 coding region, we utilized the complete genome sequences to perform a similar analysis. Tables 5 and 6 display the nucleotide positions in VP2, with non-synonymous mutations and amino acid differences, respectively, between the antigenic types CPV-2a, 2b, and 2c, along with the consensus genomes sequenced in this study. Thus, the comparison of the VP2-coding region of the CPV-2 genomes recovered here led to the clustering of samples 2, 3, 5, 6, and 7 along with the CPV2a subtype, whereas samples 1 and 4 clustered along with the CPV-2c subtype (Tables 5 and 6). Genogrouping through analysis of whole genome sequences revealed that the genomes recovered from the Tocantins state were categorized into Clade I (Figure 1) and further divided into two subclades: W3 including samples 2, 5 and 7 (OR230511, OR230514 and OR230516) and W4 comprising samples 1, 3, 4 and 6 (OR230510, OR230512, OR230513, and OR230515) (Figure 2).

**Figure 1.**
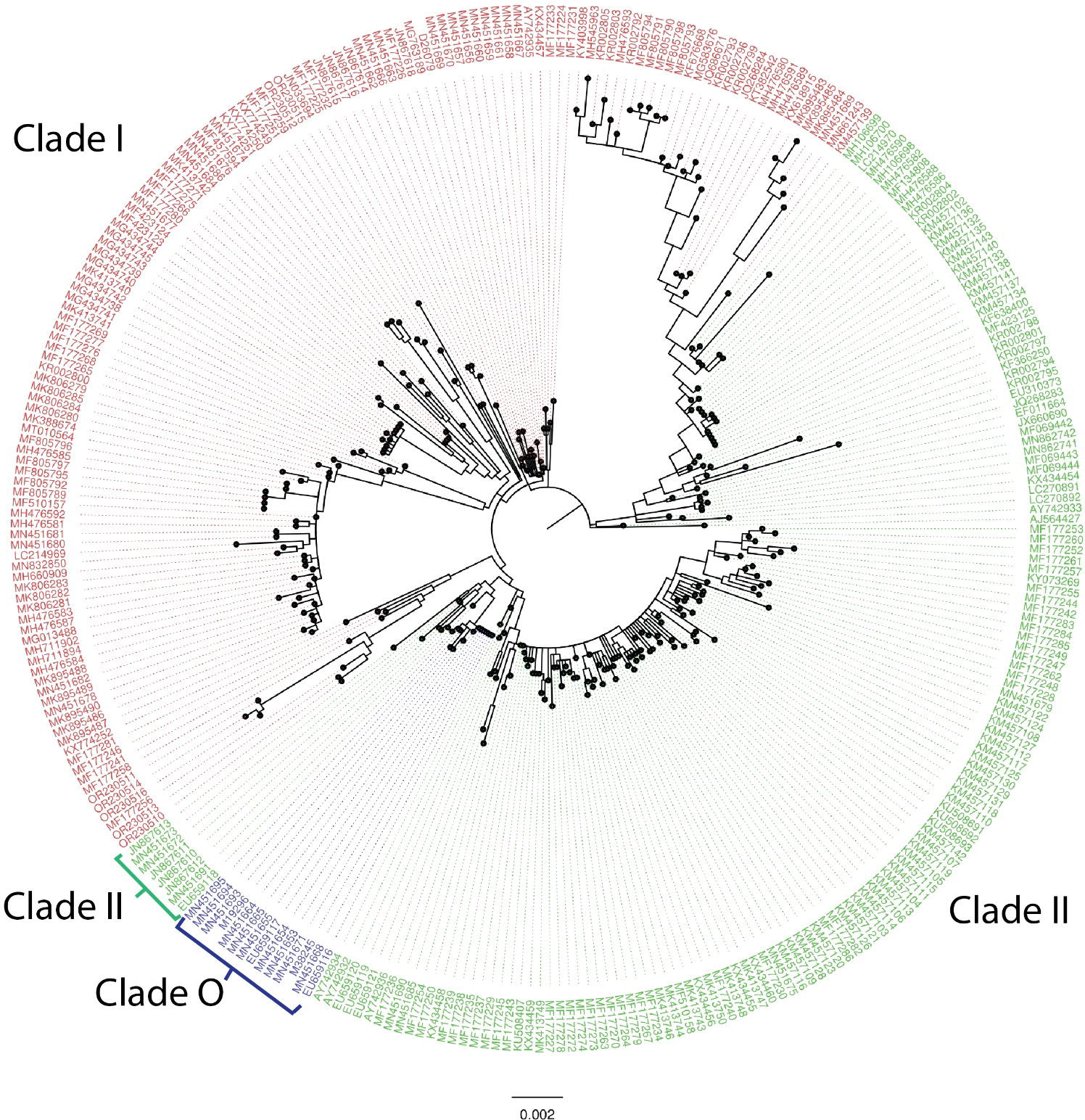
- Maximum likelihood phylogenetic tree of CPV, based on whole genome sequencing. Clade I is shown in red, II in green, and O in blue.

**Figure 2.**
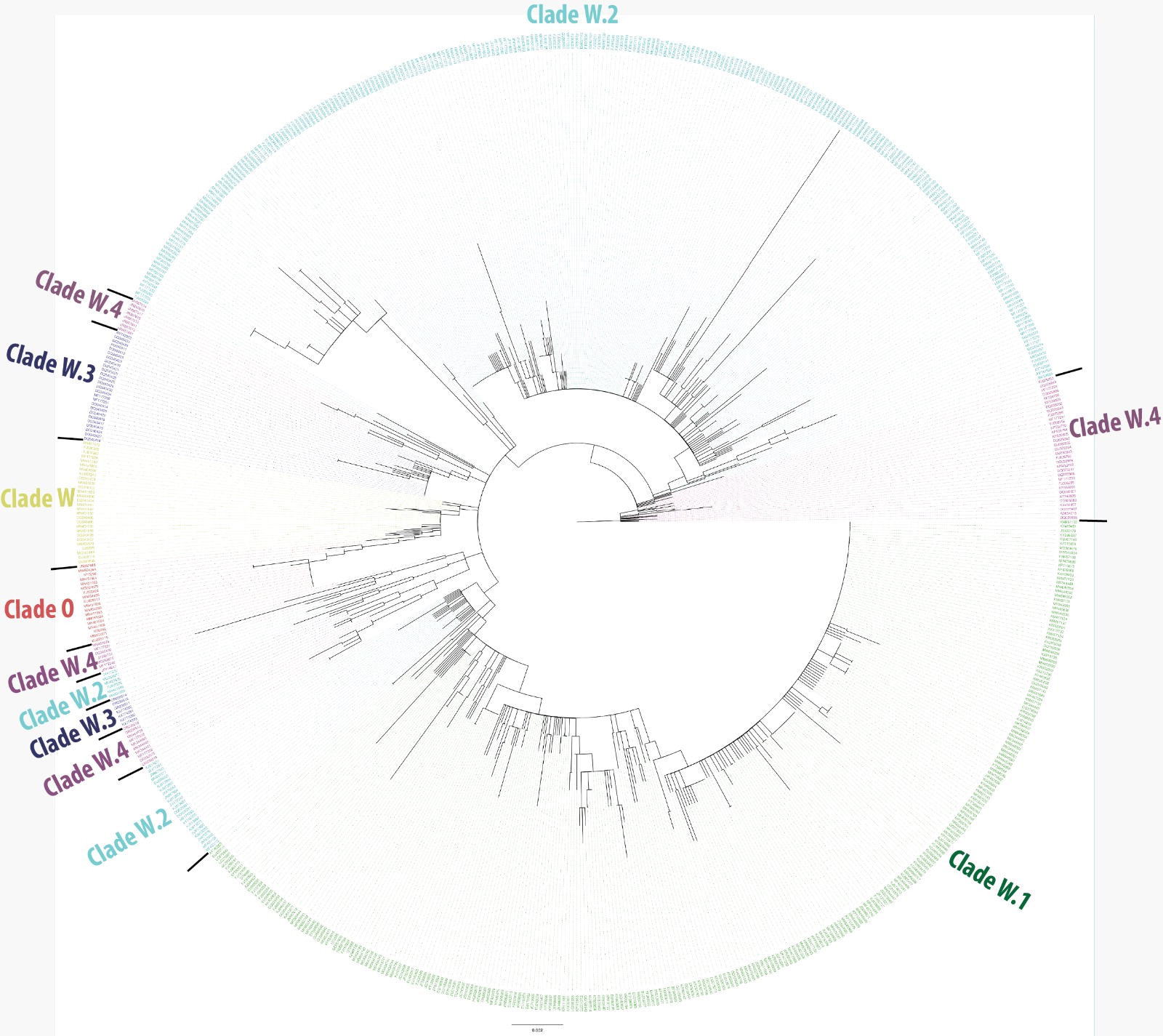
- Maximum likelihood phylogenetic tree of CPV based on the VP2 nucleotide sequence. Subclade O is shown in red, W in orange, W.1 in green, W.2 in light blue, W.3 in dark blue, and W.4 in magenta.

**Table 5.** - Comparative analysis of the CPV-2a, 2b, and 2c antigenic types, identifying non-synonymous nucleotide mutations in the VP2 gene of the CPV-2 genomes obtained in this study and reference sequences available at GenBank.

| Viruses/type | GenBank number | VP2 nucleotide positions with non-synonymous mutations |  |  |  |  |  |  |  |  |  |  |  |  |  |  |  |  |  |  |  |  |  |  |  |  |  |  |
| --- | --- | --- | --- | --- | --- | --- | --- | --- | --- | --- | --- | --- | --- | --- | --- | --- | --- | --- | --- | --- | --- | --- | --- | --- | --- | --- | --- | --- |
|  |  | 40 | 239 | 259 | 271 | 279 | 302 | 308 | 655 | 694 | 800 | 889 | 890 | 896 | 899 | 913 | 967 | 970 | 971 | 972 | 1099 | 1232 | 1276 | 1278 | 1318 | 1684 | 1691 | 1703 |
| MEV <sup>(1)</sup> | KT899746 | G | A | A | G | A | T | T | G | A | T | T | C | G | C | G | G | T | A | T | G | C | A | T | A | C | A | C |
| FPLV <sup>(2)</sup> | MZ712026 | G | A | A | G | A | C | T | A | G | T | T | C | A | C | G | G | T | A | T | G | A | A | T | A | G | A | C |
| BFPV <sup>(3)</sup> | MN451652 | G | A | A | A | A | C | T | A | G | T | T | C | G | T | G | G | T | A | T | G | A | A | T | A | G | A | C |
| CPV-2 <sup>(4)</sup> | NC_001539 | G | G | A | G | C | T | C | A | A | T | T | C | G | C | G | A | T | A | T | T | A | A | T | A | G | G | G |
| CPV-2a <sup>(5)</sup> | EU310373 | G | G | T | G | C | C | C | A | A | T | G | C | G | G | T | A | A | T | T | G | A | A | T | A | G | G | G |
| CPV-2b <sup>(6)</sup> | JQ268284 | G | G | T | G | C | C | C | A | A | A | G | C | G | G | T | A | A | T | T | G | A | G | T | G | G | G | G |
| CPV-2c <sup>(7)</sup> | KM457103 | G | G | T | G | C | C | C | A | A | T | G | C | G | G | T | A | T | T | T | G | A | G | A | A | G | G | G |
| Sample 1*(CPV-2c) | OR230510 | G | G | T | G | C | C | C | A | A | T | A | A | G | G | T | A | T | T | A | G | A | G | G | A | G | G | G |
| Sample 2* (CPV-2a) | OR230511 | G | G | T | G | C | C | C | A | A | T | G | C | G | G | T | A | T | T | A | G | A | G | C | A | G | G | G |
| Sample 3* (CPV-2a) | OR230512 | G | G | T | G | C | C | C | A | A | T | G | C | G | G | T | A | T | T | A | G | A | G | T | A | G | G | G |
| Sample 4* (CPV-2c) | OR230513 | G | G | T | G | C | C | C | A | A | T | A | A | G | G | T | A | T | T | A | G | A | G | G | A | G | G | G |
| Sample 5* (CPV-2a) | OR230514 | A | G | T | G | C | C | C | A | A | T | G | C | G | G | T | A | T | T | A | G | A | G | C | A | G | G | G |
| Sample 6* (CPV-2a) | OR230515 | G | G | T | G | C | C | C | A | A | T | G | C | G | G | T | A | T | T | A | G | A | G | T | A | G | G | G |
| Sample 7* (CPV-2a) | OR230516 | G | G | T | G | C | C | C | A | A | T | G | C | G | G | T | A | T | T | A | G | A | G | C | A | G | G | G |

(1) Mink enteritis virus strain MEV-L. (2) Feline panleukopenia virus isolate AMPV2020. (3) Feline panleukopenia virus isolate BFPV. (4) NCBI Reference Sequence. (5) Canine parvovirus isolate cpv/nj01/06 from China. (6) Canine parvovirus 2b strain CPV-LZ2 from China. (7) Canine parvovirus 2c isolate UY12 from Uruguay. \*Samples reported in this study.

**Table 6.** - Comparative analysis of the CPV-2a, 2b, and 2c antigenic types identifying amino acid variations in the VP2 protein of the CPV-2 genomes obtained in this study and reference strains available at GenBank.

| Viruses | GenBank number* | VP2 aminoacid positions |  |  |  |  |  |  |  |  |  |  |  |  |  |  |  |  |  |  |  |  |  |  |
| --- | --- | --- | --- | --- | --- | --- | --- | --- | --- | --- | --- | --- | --- | --- | --- | --- | --- | --- | --- | --- | --- | --- | --- | --- |
|  |  | 14 | 80 | 87 | 91 | 93 | 101 | 103 | 219 | 232 | 267 | 297 | 299 | 300 | 305 | 323 | 324 | 367 | 411 | 426 | 440 | 562 | 564 | 568 |
| MEV <sup>(1)</sup> | KT899746 | A | K | M | A | K | I | V | V | I | F | S | G | P | D | D | Y | D | A | N | T | L | N | A |
| FPLV <sup>(2)</sup> | MZ712026 | A | K | M | A | K | T | V | I | V | F | S | E | A | D | D | Y | D | E | N | T | V | N | A |
| BFPV <sup>(3)</sup> | MN451652 | A | k | M | T | K | T | V | I | V | F | S | G | V | D | D | Y | D | E | N | T | V | N | A |
| CPV-2 <sup>(4)</sup> | NC_001539 | A | R | M | A | N | I | A | I | I | F | S | G | A | D | N | Y | Y | E | N | T | V | S | G |
| CPV-2a <sup>(5)</sup> | EU310373 | A | R | L | A | N | T | A | I | I | F | A | G | G | Y | N | I | D | E | N | T | V | S | G |
| CPV-2b <sup>(6)</sup> | JQ268284 | A | R | L | A | N | T | A | I | I | Y | A | G | G | Y | N | I | D | E | D | A | V | S | G |
| CPV-2c <sup>(7)</sup> | KM457103 | A | R | L | A | N | T | A | I | I | F | A | G | G | Y | N | Y | D | E | E | T | V | S | G |
| Sample 1* (CPV-2c) | OR230510 | A | R | L | A | N | T | A | I | I | F | N | G | G | Y | N | L | D | E | E | T | V | S | G |
| Sample 2* (CPV-2a) | OR230511 | A | R | L | A | N | T | A | I | I | F | A | G | G | Y | N | L | D | E | D | T | V | S | G |
| Sample 3* (CPV-2a) | OR230512 | A | R | L | A | N | T | A | I | I | F | A | G | G | Y | N | L | D | E | D | T | V | S | G |
| Sample 4* (CPV-2c) | OR230513 | A | R | L | A | N | T | A | I | I | F | N | G | G | Y | N | L | D | E | E | T | V | S | G |
| Sample 5* (CPV-2a) | OR230514 | T | R | L | A | N | T | A | I | I | F | A | G | G | Y | N | L | D | E | D | T | V | S | G |
| Sample 6* (CPV-2a) | OR230515 | A | R | L | A | N | T | A | I | I | F | A | G | G | Y | N | L | D | E | D | T | V | S | G |
| Sample 7* (CPV-2a) | OR230516 | A | R | L | A | N | T | A | I | I | F | A | G | G | Y | N | L | D | E | D | T | V | S | G |

(1) Mink enteritis virus strain MEV-L. (2) Feline panleukopenia virus isolate AMPV2020. (3) Feline panleukopenia virus isolate BFPV. (4) NCBI Reference Sequence. (5) Canine parvovirus isolate cpv/nj01/06 from China. (6) Canine parvovirus 2b strain CPV-LZ2 from China. (7) Canine parvovirus 2c isolate UY12 from Uruguay. \*Samples reported in this study.

### 3.2 Bayesian analysis

The temporal analysis of the CPV sequences showed a sufficient temporal signal between the genetic divergence from root to tip against the sampling time after outlier removal (with R2 = 0.71 and a correlation coefficient of 0.84; Supplementary Figure S1), supporting the use of temporal calibration directly from the sequences. The evolutionary rate estimated in this study was 2.04 × 10^−4^ substitutions per site, per year (95% highest posterior density interval-HDP: 1.73 × 10^−4^ to 2.39 × 10^−4^ substitutions per site, per year). The time of the most recent common ancestor (TMRCA) for the phylogenetic evolutionary tree was generated 42 years ago (1980), based on the most recently analyzed sequence (2023) (95%: HDP: 1977 to 1981) (Figure 3). The TMRCA for the CPV-2c Tocantins clade was estimated to be around July 2017 (95%: HDP: October 2014 to April 2020) and was grouped closely to a previously Brazilian Canine parvovirus/CPV genome identified in 2013 (MF177259). Curiously, the CPV-2a genomes were grouped into two different subclades (Figure 3). A subclade with two Tocantins samples (OR230510 and OR230513) that grouped closely to two previously sequenced Brazilian genomes obtained in 2013 (MF177256 and MF177258) and five samples of institutionalized dogs from Brazilian Amazon from 2019 (OP093952, OP093953, OP093954, OP093955, OP093956). We also observed another single, well-supported subclade (posterior probability = 1.0) with three Tocantins samples, which can be an indication of local transmission. The TMRCA for the subclade with three Tocantins samples was estimated to be around February 2020 (95%: HDP: March 2018 to August 2021), while the TMRCA for the subclade with two sequences was estimated to be around July 2021 (95%: HDP: December 2019 to November 2021).

**Figure 3.**
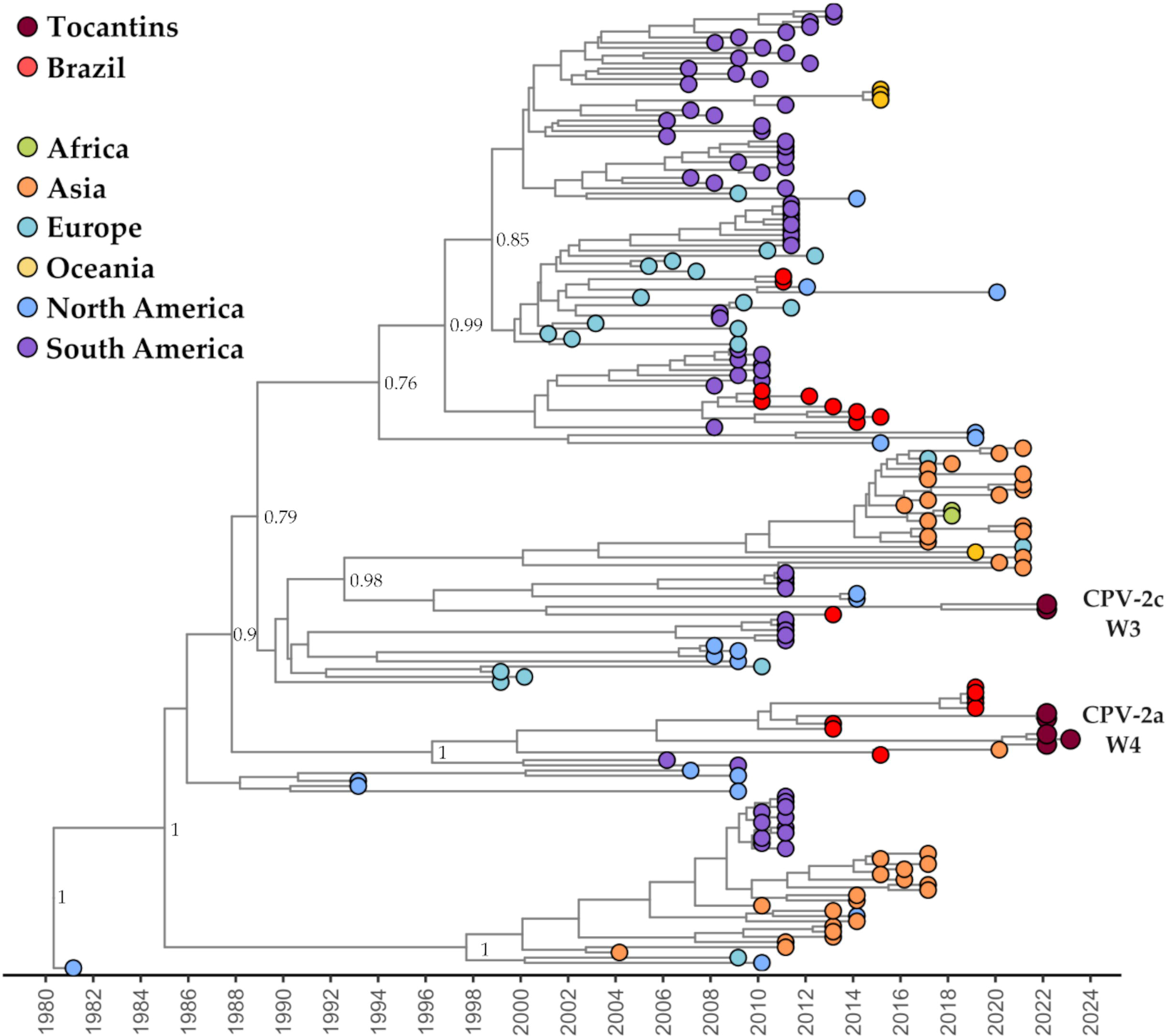
Time-scaled phylogenetic tree of 185 complete and near-complete CPV genome sequences available at GenBank (accession date 20 July 2023). Colors represent different sampling locations according to the legend on the left of the tree. Genomes reported in the presented study are colored in brown.

## 4. Discussion

In this study, we adapted two methods for directly amplifying complete genomes of CPV from clinical samples. The methodology was similar in both, except that multiplex-PCR primers targeting amplicons of different sizes, namely 400-bp and 1000-bp, were employed. Our findings clearly demonstrate that amplification targeting shorter fragments (400 bp) outperformed the method based on amplification of longer fragments (1000 bp) in terms of number of reads and genome coverage. Nevertheless, both strategies were similarly effective in successfully amplifying complete CPV genomes. Furthermore, by combining the two amplification strategies with nanopore sequencing, we were able to obtain complete sequencing of the genomes’ 5’ and 3’ hairpin ends, which are often missed in similar studies (Rosales-Munar et al., 2020; Wang et al., 2021). The sequencing of complete genomes, including the challenging hairpin ends, is a significant advantage when compared to previously employed strategies (Ogbu et al., 2020; Pérez et al., 2014).

Various methodologies have been developed to amplify specific genes or the entire CPV genome (Calderon et al., 2007; Ogbu et al., 2020; Prateli et al., 2000; Pérez et al., 2014; Zhou et al., 2016). However, these methods have often resulted in partial genomes, most likely due to the challenge represented by the sequencing of the hairpin regions, or the lack of specific primers designed for these regions. In a recent study (Temizkan and Sevinc Temizkan, 2023), five complete CPV genomes were obtained using NGS Illumina technologies without primer enrichment, as proposed in the methodology reported here. However, the utilization of primer enrichment in our methodology provides a targeted amplification approach, specifically designed to overcome challenges associated with the non-translated regions with hairpin structures. This enrichment step increases the likelihood of obtaining complete genomes, as it enhances the amplification efficiency and coverage of CPV genomes. Therefore, the methodology introduced here offers a distinct advantage by facilitating the generation of complete CPV genomes, which is undoubtedly preferred for more comprehensive genetic analyses of this viral pathogen. In addition, the application of full or nearly full genome sequencing to CPV studies may provide a significant contribution to a more precise diagnosis and typing/subtyping of CPV, which on its turn may aid in clarifying aspects of viral pathogenesis and its associations with particular genotypes. Nevertheless, further investigations involving a larger sample size, including control groups and additional dogs without clinical signs, would be necessary to provide a more comprehensive assessment of the diagnostic performance and sensitivity of our method.

Most PCR-based methodologies used to obtain CPV genomes have been based on the amplification of specific genes, such as the VP2 gene, generating fragments of around 1745 bp (Battilani et al., 2001; 2006; Mokizuki et al., 1996) or fragments around 2225 bp (NS1) and 2788 bp (VP2) aiming to capture the whole-genome sequence, resulting in partial genome coverage (Pérez et al., 2014). In contrast, the two strategies presented in this study, involving multiple fragments of 400 bp and 1000 bp, proved effective in amplifying complete CPV genomes. It is conceivable that by conducting separate sequencing and utilizing the Nanopore Rapid Barcode Kit, the 1000 bp approach might have yielded a comparable number of reads and coverage to that of the 400 bp fragments. Notably, nanopore sequencing enabled the sequencing of the hairpin region of the genome, potentially converting single-stranded (ss) DNA to double-stranded (ds) DNA for this short, initial part of the genome. This feature of nanopore sequencing allows for sequencing from the ds end of the genome and then extending "backwards" to the nucleic acid end, as suggested by Balazs Harrach (personal communication, 2023).

Sequencing of full genomes allowed for the unequivocal determination of the genogroups of the viruses whose genomes were recovered in this study. The classic amino acid changes in the CPV-2a genome that gave rise to CPV-2b (N426D, Parrish et al., 1991), and later on, the VP2 substitution that generated CPV-2c (D426E, Martella et al., 2004) were detected in the sequenced genomes (Tables 5 and 6). Additionally, the majority of samples exhibited an additional non-synonymous mutation, S297A, previously reported by Ohshima et al. (2008). Two samples, 1 and 4, classified as CPV-2c, displayed a novel alteration, S297N. Moreover, a consistent shift was identified at position 324 (Y324L) across all samples, specifically affecting a loop region within the VP2 protein structure. This flexible loop region plays a crucial role in protein function and interactions. The significance of these observed changes in terms of viral pathogenicity and epidemiology is yet to be determined, and further studies will provide valuable insights into unraveling these aspects and their potential implications. In light of these findings, it is worth noting that a recent study conducted in Brazil identified CPV-2 sequences spread across all four clades, indicating substantial genetic diversity of this virus within the country (De Oliveira Santana et al., 2022). However, in order to achieve appropriate epidemiological support, the number of CPV genomes available for classification still needs to be increased. Furthermore, the limited number of genomes has hindered a more precise taxonomic separation, as a monophyletic branch with only a few sequences may represent part of a clade, which can be better resolved with the sequencing of additional genomes. Therefore, a comprehensive analysis involving a larger dataset of CPV genomes would provide a more accurate understanding of the virus’s phylogenetic structure, including a more precise assessment of monophyletic branch formation and its relevance in taxonomic classification.

Various factors have influenced CPV evolution, including natural selection, mutation rates, and host immunity (Li et al., 2022). The classification of CPV into genogroups (De Oliveira Santana et al., 2022) has been challenging due to several factors: genetic diversity, rapid evolution, recombination events, and complex phylogenetic relationships (Truyen, 2006). As shown here, the advancements in sequencing technologies can aid in understanding CPV’s genetic diversity. Instead of focusing solely on the VP2 gene, sequencing of the entire genome can provide (i) a more comprehensive view of their genetic makeup, (ii) allow analyses of multiple genes and regions, and (iii) enable more accurate classification and understanding of the evolutionary relationships among strains. Furthermore, ongoing surveillance of CPV strains is crucial to monitor the emergence of new variants, in attempting to improve our understanding of the evolution of such virus (De Oliveira Santana et al., 2022).

The time estimate of the circulation of the most recent common ancestor (TMRCA) of CPV-2 was calculated to be 43 years ago (1980). This estimate, based on the most recently analyzed sequence (2022), is consistent with a previous study by Giraldo-Ramirez et al. (2020), which estimated the TMRCA for CPV-2 to have occurred in 1979 (44 years ago) based on available genomes up to 2019. These findings support the hypothesis that CPV-2 likely emerged and diverged from an ancestral FPV in the early 1970s (Kelly, 1978). Additionally, the estimated evolutionary rate of 2.06 × 10^−4^ substitutions per site, per year in this study is similar to the rates observed in other RNA viruses (Holmes, 2003). For instance, the evolutionary rate of the Dengue virus is estimated to be around 6-8 × 10^−4^ substitutions per site, per year (Weaver and Vasilakis, 2009). In contrast, the SARS-CoV-2 virus, which causes COVID-19, has shown a higher evolutionary rate, estimated to be around 10^−3^ to 10^−4^ substitutions per site, per year (Porter et al., 2023; Wang et al., 2022). It’s important to note that these rates can vary depending on the specific viral lineage, geographic region, and time period analyzed.

Despite the limitations inherent to this study, which involved adapting previous methodology to allow sequencing of a relatively small number of samples, our findings unequivocally establish the effectiveness of both strategies (400 and 1000 bp with overlapping) in successfully amplifying the complete CPV genome sequence, including the intricate hairpin ends. The choice between these strategies may depend on specific research goals and available resources. In conclusion, the methodology presented in this study provides a valuable strategy for sequencing whole CPV genomes. This is expected to contribute to the future development of improved prevention and treatment strategies for this debilitating canine disease.

## Supplementary Materials

**Supplementary Figure 1** in PDF to read the sequence accession numbers. **Supplementary Figure 2** in PDF to read the sequence accession numbers. **Figure S1**. Root-to-tip regression of genetic distances and sampling dates for 185 CPV genomes in the final dataset. **Table S1**. CPV sequences available in GenBank on December of 2022. **Table S2**. CPV whole genome sequences available in GenBank used in phylogenetic analyses. **Table S3**. CPV VP2 nucleotide sequence available in GenBank used in phylogenetic analyses. **Table S4**. Details of the 185 complete and near-complete CPV genome sequences sampled from NCBI and used in the Bayesian phylogenetic reconstruction. **Table S5**. Model comparison of strict molecular clock, uncorrelated relaxed clock and demographic growth models through path sampling (PS) and stepping stone (SS) methods.

## Author Contributions

Conceived and designed the experiments: S.F.A.S., U.J.B.d.S., and F.S.C. Performed the experiments: S.F.A.S., and U.J.B.d.S. Analyzed the data: U.J.B.d.S., M.T.O., J.J., F.R.S. and F.S.C. Contributed to the writing of the manuscript: U.J.B.d.S., M.T.O., J.J., F.R.S., A.C.F., P.M.R., and F.S.C. All authors have read and agreed to the published version of the manuscript.

## Informed Consent Statement

Prior to participating in the study, dog owners provided signed informed consent letters. Every effort was made to minimize stress and discomfort for the patients during the sample collection process. The procedures for sample collection strictly adhered to the guidelines outlined by the Brazilian government’s law no. 11.794/2008, also known as the "Law on the Use of Animals in Scientific Research". This included ensuring the use of appropriate techniques and equipment, maintaining a sterile environment, and prioritizing the well-being and safety of the dogs throughout the entire process.

## Data Availability Statement

The authors affirm that all data substantiating the discoveries in this study are provided within the paper. Additionally, we have deposited the raw data (reads) utilized for CPV genome assembly on the SRA platform, with the following accessions: PRJNA989242 for the bioproject, and SAMN36082971, SAMN36082972, SAMN36082973, SAMN36082974, SAMN36082975, SAMN36082976, SAMN36082977 for the biosamples. The protocol details are available on protocols.io: https://protocols.io/view/sequencing-of-canine-parvovirus-cpv-from-rectal-sw-cwj6xcre

## Acknowledgments

U.J.B.d.S. and M.T.O. were granted a post-doctoral scholarship (DTI-A) from CNPq. F.R.S., A.C.F., P.M.R., and F.S.C. are CNPq research fellows.

