## Supplementary figures and images for "Recovery of complete genomes of canine parvovirus from clinical samples"

### Supplementary Figure 1

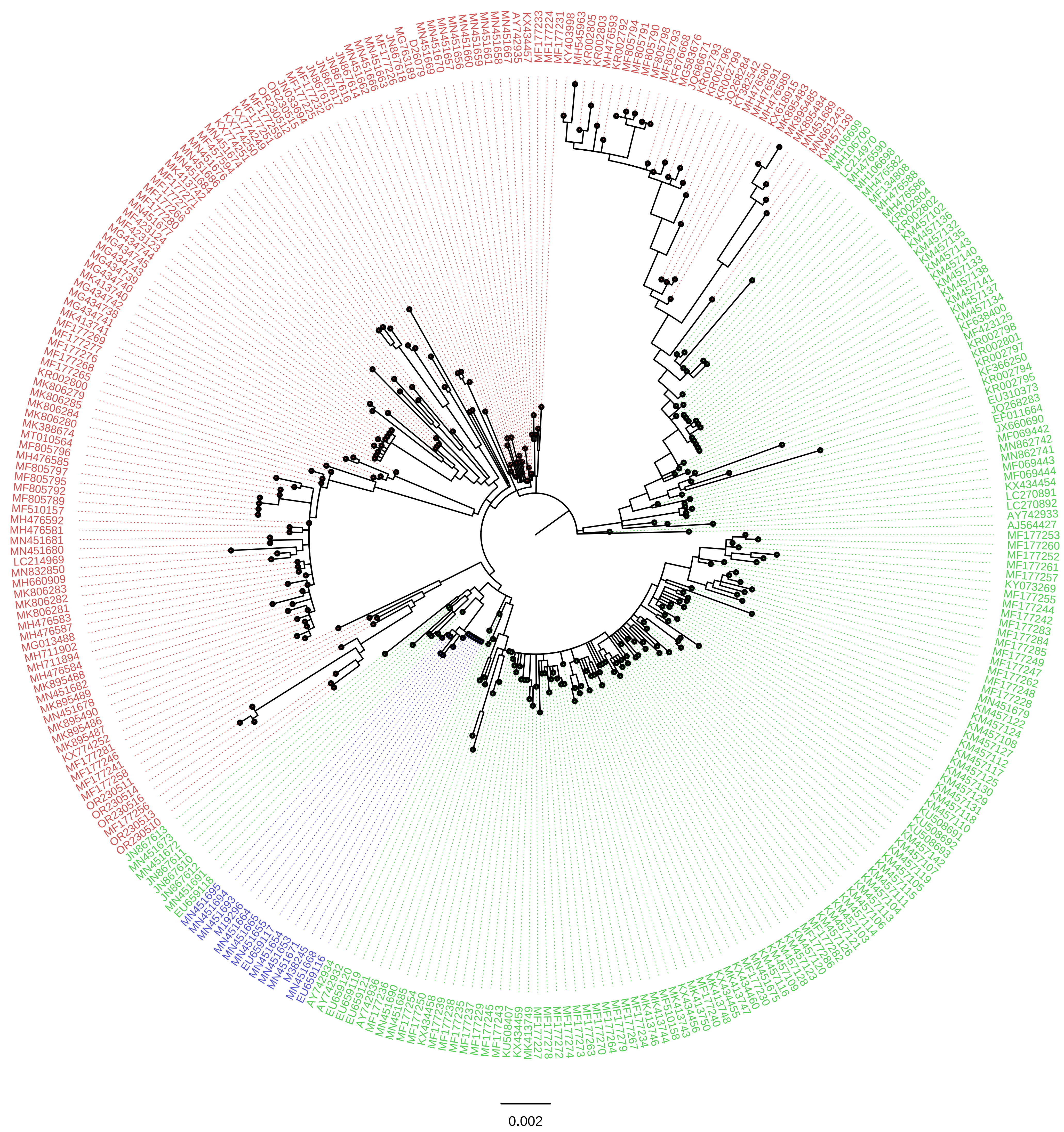

### Supplementary Figure 2

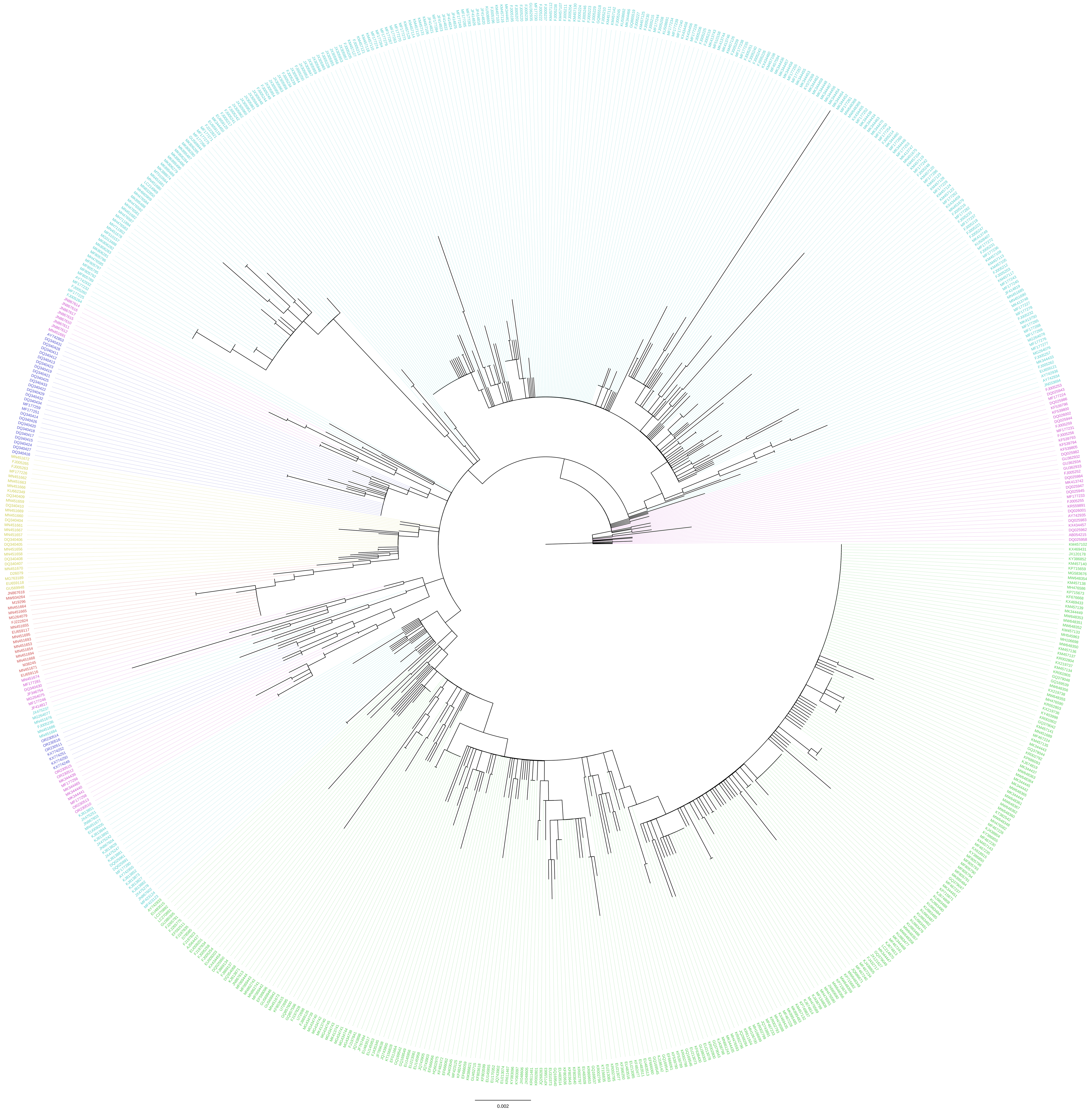
