## Supplementary Figure S1 for "Recovery of complete genomes of canine parvovirus from clinical samples"

**Figure S1**. Root-to-tip regression of genetic distances and sampling dates for 185 CPV genomes in the final dataset. Correlation coefficient (r) and r squared are depicted above the graph.

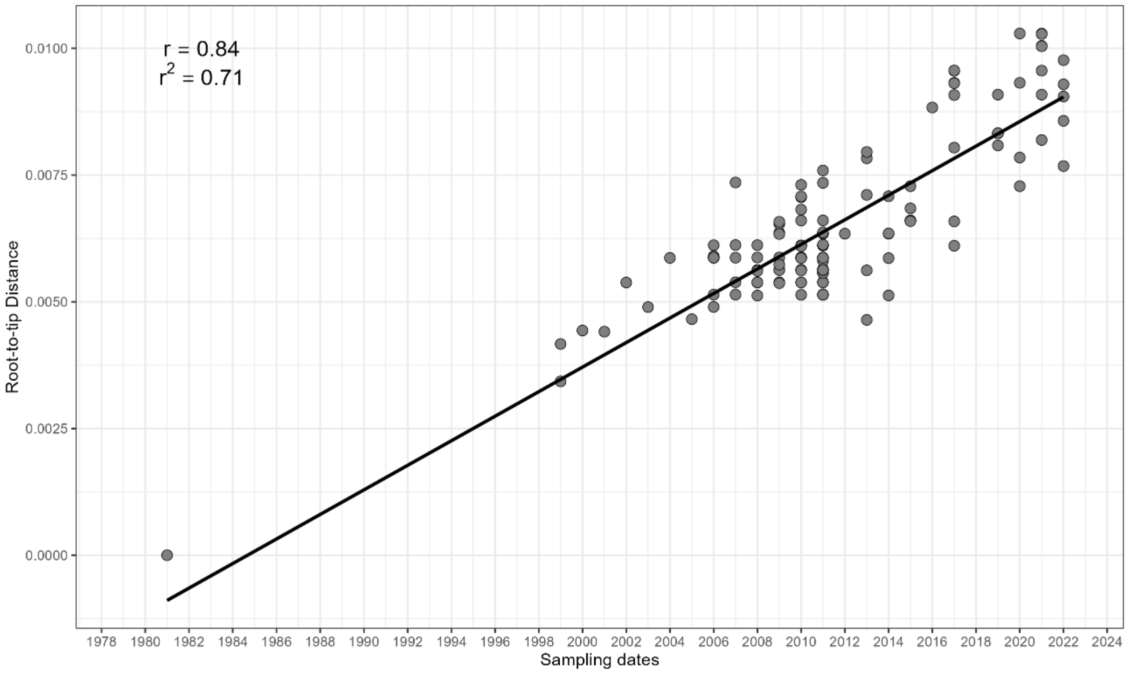
