## Supplementary Table S1 for "Recovery of complete genomes of canine parvovirus from clinical samples"

**Supplementary Table S1 -** CPV sequences available in GenBank on December of 2022.

| Quant. | Accession number | Description | Length (bp) | % GC |
|---|---|---|---|---|
| 1 | D26079 | Canine parvovirus DNA, complete genome | 5075 | 36.9% |
| 2 | EF011664 | Canine parvovirus nonstructural protein 1 (NS1), nonstructural protein 2 (NS2), VP1, and VP2 genes, complete cds | 4623 | 36.1% |
| 3 | EU310373 | Canine parvovirus isolate cpv/nj01/06, complete genome | 4559 | 36.0% |
| 4 | JN033694 | Canine parvovirus strain Laika-1993, complete genome | 5126 | 36.9% |
| 5 | JQ268283 | Canine parvovirus 2a strain CPV-LZ1, complete genome | 5053 | 37.0% |
| 6 | JQ268284 | Canine parvovirus 2b strain CPV-LZ2, complete genome | 5053 | 37.1% |
| 7 | JX660690 | Canine parvovirus isolate SC02/2011 non-structural protein 1 (NS1), non-structural protein 2 (NS2), capsid protein 1 (VP1), and capsid protein 2 (VP2) genes, complete cds | 4699 | 36.0% |
| 8 | KF638400 | Canine parvovirus isolate s5 nonstructural protein NS1 (NS1), nonstructural protein NS2 (NS2), VP1 protein (VP1), and VP2 protein (VP2) genes, complete cds | 4702 | 36.0% |
| 9 | KM457102 | Canine parvovirus 2a isolate UY243, complete genome | 4269 | 36.5% |
| 10 | KM457103 | Canine parvovirus 2c isolate UY12, complete genome | 4269 | 36.4% |
| 11 | KM457104 | Canine parvovirus 2c isolate UY47, complete genome | 4269 | 36.5% |
| 12 | KM457105 | Canine parvovirus 2c isolate UY52, complete genome | 4269 | 36.4% |
| 13 | KM457106 | Canine parvovirus 2c isolate UY55, complete genome | 4269 | 36.4% |
| 14 | KM457107 | Canine parvovirus 2c isolate UY72, complete genome | 4269 | 36.4% |
| 15 | KM457108 | Canine parvovirus 2c isolate UY82, complete genome | 4269 | 36.4% |
| 16 | KM457109 | Canine parvovirus 2c isolate UY95, complete genome | 4269 | 36.5% |
| 17 | KM457110 | Canine parvovirus 2c isolate UY101, complete genome | 4269 | 36.6% |
| 18 | KM457111 | Canine parvovirus 2c isolate UY120, complete genome | 4269 | 36.5% |
| 19 | KM457112 | Canine parvovirus 2c isolate UY135, complete genome | 4269 | 36.4% |
| 20 | KM457113 | Canine parvovirus 2c isolate UY152, complete genome | 4269 | 36.5% |
| 21 | KM457114 | Canine parvovirus 2c isolate UY169, complete genome | 4269 | 36.5% |
| 22 | KM457115 | Canine parvovirus 2c isolate UY173, complete genome | 4269 | 36.6% |
| 23 | KM457116 | Canine parvovirus 2c isolate UY185, complete genome | 4269 | 36.5% |
| 24 | KM457117 | Canine parvovirus 2c isolate UY187, complete genome | 4269 | 36.5% |
| 25 | KM457118 | Canine parvovirus 2c isolate UY190, complete genome | 4269 | 36.5% |
| 26 | KM457119 | Canine parvovirus 2c isolate UY235, complete genome | 4269 | 36.4% |
| 27 | KM457120 | Canine parvovirus 2c isolate UY242, complete genome | 4269 | 36.4% |
| 28 | KM457121 | Canine parvovirus 2c isolate UY247, complete genome | 4269 | 36.4% |
| 29 | KM457122 | Canine parvovirus 2c isolate UY258, complete genome | 4269 | 36.4% |
| 30 | KM457123 | Canine parvovirus 2c isolate UY261, complete genome | 4269 | 36.4% |
| 31 | KM457124 | Canine parvovirus 2c isolate UY307, complete genome | 4269 | 36.4% |
| 32 | KM457125 | Canine parvovirus 2c isolate UY317, complete genome | 4269 | 36.4% |
| 33 | KM457126 | Canine parvovirus 2c isolate UY318, complete genome | 4269 | 36.4% |
| 34 | KM457127 | Canine parvovirus 2c isolate UY326, complete genome | 4269 | 36.4% |
| 35 | KM457128 | Canine parvovirus 2c isolate UY346, complete genome | 4269 | 36.4% |
| 36 | KM457129 | Canine parvovirus 2c isolate UY349, complete genome | 4269 | 36.4% |
| 37 | KM457130 | Canine parvovirus 2c isolate UY354, complete genome | 4269 | 36.4% |
| 38 | KM457131 | Canine parvovirus 2c isolate UY368, complete genome | 4269 | 36.5% |
| 39 | KM457132 | Canine parvovirus 2a isolate UY245, complete genome | 4269 | 36.5% |
| 40 | KM457133 | Canine parvovirus 2a isolate UY250, complete genome | 4269 | 36.5% |

| | | | | |
|---|---|---|---|---|
| 41 | KM457134 | Canine parvovirus 2a isolate UY280, complete genome | 4269 | 36.5% |
| 42 | KM457135 | Canine parvovirus 2a isolate UY306, complete genome | 4269 | 36.5% |
| 43 | KM457136 | Canine parvovirus 2a isolate UY315, complete genome | 4269 | 36.5% |
| 44 | KM457137 | Canine parvovirus 2a isolate UY344, complete genome | 4269 | 36.5% |
| 45 | KM457138 | Canine parvovirus 2a isolate UY363, complete genome | 4269 | 36.5% |
| 46 | KM457139 | Canine parvovirus 2a isolate recUY364, complete genome | 4269 | 36.5% |
| 47 | KM457140 | Canine parvovirus 2a isolate UY365, complete genome | 4269 | 36.5% |
| 48 | KM457141 | Canine parvovirus 2a isolate UY370a, complete genome | 4269 | 36.5% |
| 49 | KM457142 | Canine parvovirus 2c isolate UY370c, complete genome | 4269 | 36.5% |
| 50 | KM457143 | Canine parvovirus 2a isolate UY364, complete genome | 4269 | 36.5% |
| 51 | KR002792 | Canine parvovirus 2a strain CPV/CN/SH1/2013, complete genome | 4269 | 36.6% |
| 52 | KR002793 | Canine parvovirus 2b strain CPV/CN/HB1/2013, complete genome | 4269 | 36.6% |
| 53 | KR002794 | Canine parvovirus 2a strain CPV/CN/HB3/2013, complete genome | 4269 | 36.6% |
| 54 | KR002795 | Canine parvovirus 2a strain CPV/CN/JL1/2013, complete genome | 4269 | 36.5% |
| 55 | KR002796 | Canine parvovirus 2b strain CPV/CN/JL3/2013, complete genome | 4269 | 36.6% |
| 56 | KR002797 | Canine parvovirus 2a strain CPV/CN/JL4/2013, complete genome | 4269 | 36.6% |
| 57 | KR002798 | Canine parvovirus 2a strain CPV/CN/JL5/2013, complete genome | 4269 | 36.4% |
| 58 | KR002799 | Canine parvovirus 2b strain CPV/CN/JL6/2013, complete genome | 4269 | 36.6% |
| 59 | KR002800 | Canine parvovirus 2a strain CPV/CN/LN1/2014, complete genome | 4269 | 36.6% |
| 60 | KR002801 | Canine parvovirus 2a strain CPV/CN/SD6/2014, complete genome | 4269 | 36.5% |
| 61 | KR002802 | Canine parvovirus 2a strain CPV/CN/SD9/2014, complete genome | 4269 | 36.5% |
| 62 | KR002803 | Canine parvovirus 2a strain CPV/CN/SD10/2014, complete genome | 4269 | 36.6% |
| 63 | KR002804 | Canine parvovirus 2a strain CPV/CN/SD18/2014, complete genome | 4269 | 36.5% |
| 64 | KR002805 | Canine parvovirus 2a strain CPV/CN/SD19/2014, complete genome | 4269 | 36.6% |
| 65 | KT382542 | Canine parvovirus isolate CPV-SH14, complete genome | 5062 | 37.2% |
| 66 | KT899746 | Mink enteritis virus strain MEV-L, complete genome | 5220 | 36.4% |
| 67 | KY073269 | Canine parvovirus 2c isolate UFMT, complete genome | 5318 | 35.7% |
| 68 | KY403998 | Canine parvovirus strain CPV-YH, complete genome | 4921 | 36.7% |
| 69 | M19296.1 | Canine parvovirus strain CPV-N, complete cds | 5323 | 35.6% |
| 70 | MF069442 | Canine parvovirus strain CPV/Raccoon/RC14/BC_2010, complete genome | 4468 | 35.8% |
| 71 | MF069443 | Canine parvovirus strain CPV/Raccoon/RC19/BC_2016, complete genome | 4468 | 35.8% |
| 72 | MF069444 | Canine parvovirus strain CPV/Raccoon/RC20/BC_2016, complete genome | 4459 | 35.9% |
| 73 | MF423123 | Canine parvovirus isolate CPV/Coyote/C16/NL_2014, complete genome | 4468 | 36.0% |
| 74 | MF423124 | Canine parvovirus isolate CPV/Coyote/C55/NL_2014, complete genome | 4468 | 36.0% |
| 75 | MF423125 | Canine parvovirus isolate CPV/Coyote/C67/NL_2014, complete genome | 4468 | 35.9% |
| 76 | MF457594 | Canine parvovirus strain OH20219, complete genome | 4306 | 36.4% |
| 77 | MF805789 | Canine parvovirus strain Canine/China/01/2016, complete genome | 5051 | 37.0% |
| 78 | MF805790 | Canine parvovirus strain Canine/China/02/2016, complete genome | 5051 | 37.1% |
| 79 | MF805791 | Canine parvovirus strain Canine/China/03/2016, complete genome | 5051 | 37.1% |
| 80 | MF805792 | Canine parvovirus strain Canine/China/04/2016, complete genome | 5051 | 37.1% |
| 81 | MF805793 | Canine parvovirus strain Canine/China/05/2016, complete genome | 5051 | 37.0% |
| 82 | MF805794 | Canine parvovirus strain Canine/China/06/2016, complete genome | 5051 | 37.1% |
| 83 | MF805795 | Canine parvovirus strain Canine/China/07/2016, complete genome | 5051 | 37.1% |
| 84 | MF805796 | Canine parvovirus strain Canine/China/08/2016, complete genome | 5051 | 37.0% |
| 85 | MF805797 | Canine parvovirus strain Canine/China/09/2016, complete genome | 5051 | 37.0% |
| 86 | MF805798 | Canine parvovirus strain Canine/China/10/2016, complete genome | 5051 | 37.0% |
| 87 | MG013488 | Canine parvovirus isolate CPV-SH1516, complete genome | 5059 | 37.0% |

| | | | | |
|---|---|---|---|---|
| 88 | MG583676 | Canine parvovirus 2a strain CPV/CN/ya1/2017, complete genome | 4666 | 36.0% |
| 89 | MG763189 | Canine parvovirus isolate CPV-L, complete genome | 4774 | 35.6% |
| 90 | MH106698 | Canine parvovirus isolate CPV-BJL1, complete genome | 5054 | 37.0% |
| 91 | MH106699 | Canine parvovirus isolate CPV-BJL2, complete genome | 5056 | 37.1% |
| 92 | MH106700 | Canine parvovirus isolate CPV-BJL3, complete genome | 5054 | 37.1% |
| 93 | MH476580 | Canine parvovirus strain Canine/China/11/2017, complete genome | 5053 | 37.1% |
| 94 | MH476581 | Canine parvovirus strain Canine/China/12/2017, complete genome | 5054 | 37.0% |
| 95 | MH476582 | Canine parvovirus strain Canine/China/13/2017, complete genome | 5053 | 37.1% |
| 96 | MH476583 | Canine parvovirus strain Canine/China/14/2017, complete genome | 5053 | 37.0% |
| 97 | MH476584 | Canine parvovirus strain Canine/China/15/2017, complete genome | 5053 | 37.0% |
| 98 | MH476585 | Canine parvovirus strain Canine/China/16/2017, complete genome | 5051 | 37.0% |
| 99 | MH476586 | Canine parvovirus strain Canine/China/17/2017, complete genome | 5053 | 37.1% |
| 100 | MH476587 | Canine parvovirus strain Canine/China/18/2017, complete genome | 5051 | 37.1% |
| 101 | MH476588 | Canine parvovirus strain Canine/China/19/2017, complete genome | 5052 | 37.1% |
| 102 | MH476589 | Canine parvovirus strain Canine/China/20/2017, complete genome | 5053 | 37.1% |
| 103 | MH476590 | Canine parvovirus strain Canine/China/21/2017, complete genome | 5051 | 37.1% |
| 104 | MH476591 | Canine parvovirus strain Canine/China/22/2017, complete genome | 5053 | 37.1% |
| 105 | MH476592 | Canine parvovirus strain Canine/China/23/2017, complete genome | 5052 | 37.1% |
| 106 | MH476593 | Canine parvovirus strain Canine/China/24/2017, complete genome | 5050 | 37.1% |
| 107 | MH545963 | Canine parvovirus 2a strain TN/CPV2a/2018, complete genome | 4921 | 36.8% |
| 108 | MH660909 | Canine parvovirus 2c strain 5 MGL, complete genome | 5075 | 36.9% |
| 109 | MH711894 | Canine parvovirus 2c strain CU24, complete genome | 4269 | 36.5% |
| 110 | MH711902 | Canine parvovirus 2c strain CU21, complete genome | 4269 | 36.5% |
| 111 | MK144544 | Canine parvovirus 2c isolate K01708-1 NS1, VP1, and VP2 genes, complete cds | 4269 | 36.6% |
| 112 | MK144545 | Canine parvovirus 2a isolate K01708-2 NS1, VP1, and VP2 genes, complete cds | 4269 | 36.7% |
| 113 | MK144546 | Canine parvovirus 2a isolate K01708-3 NS1, VP1, and VP2 genes, complete cds | 4269 | 36.6% |
| 114 | MK388674 | Canine parvovirus isolate HB2017, complete genome | 4723 | 36.2% |
| 115 | MK895483 | Carnivore protoparvovirus 1 strain IZSSI_PA1464/19_idUV1 nonstructural protein NS1 (NS1), nonstructural protein NS2 (NS2), structural protein VP1 (VP1), and structural protein VP2 (VP2) genes, complete cds | 4269 | 36.6% |
| 116 | MK895484 | Carnivore protoparvovirus 1 strain IZSSI_PA1464/19_idYV8 nonstructural protein NS1 (NS1), nonstructural protein NS2 (NS2), structural protein VP1 (VP1), and structural protein VP2 (VP2) genes, complete cds | 4269 | 36.7% |
| 117 | MK895485 | Carnivore protoparvovirus 1 strain IZSSI_PA1464/19_idUV6 nonstructural protein NS1 (NS1), nonstructural protein NS2 (NS2), structural protein VP1 (VP1), and structural protein VP2 (VP2) genes, complete cds | 4269 | 36.6% |
| 118 | MK895486 | Carnivore protoparvovirus 1 strain IZSSI_PA1464/19_idYV2 nonstructural protein NS1 (NS1), nonstructural protein NS2 (NS2), structural protein VP1 (VP1), and structural protein VP2 (VP2) genes, complete cds | 4269 | 36.5% |
| 119 | MK895487 | Carnivore protoparvovirus 1 strain IZSSI_PA1464/19_idEV8 nonstructural protein NS1 (NS1), nonstructural protein NS2 (NS2), structural protein VP1 (VP1), and structural protein VP2 (VP2) genes, complete cds | 4269 | 36.6% |

| | | | | |
|---|---|---|---|---|
| 120 | MK895488 | Carnivore protoparvovirus 1 strain IZSSI_PA1464/19_idJOE2 nonstructural protein NS1 (NS1), nonstructural protein NS2 (NS2), structural protein VP1 (VP1), and structural protein VP2 (VP2) genes, complete cds | 4269 | 36.6% |
| 121 | MK895489 | Carnivore protoparvovirus 1 strain IZSSI_PA1464/19_idNC nonstructural protein NS1 (NS1), nonstructural protein NS2 (NS2), structural protein VP1 (VP1), and structural protein VP2 (VP2) genes, complete cds | 4269 | 36.6% |
| 122 | MK895490 | Carnivore protoparvovirus 1 strain IZSSI_PA1464/19_idPSV21 nonstructural protein NS1 (NS1), nonstructural protein NS2 (NS2), structural protein VP1 (VP1), and structural protein VP2 (VP2) genes, complete cds | 4269 | 36.5% |
| 123 | MN451652 | Feline panleukopenia virus isolate BFPV, complete genome | 4269 | 36.0% |
| 124 | MN451653 | Canine parvovirus isolate CPV6, complete genome | 4269 | 36.2% |
| 125 | MN451654 | Canine parvovirus isolate CPV9, complete genome | 4269 | 36.2% |
| 126 | MN451655 | Canine parvovirus isolate CPV12, complete genome | 4269 | 36.3% |
| 127 | MN451656 | Canine parvovirus isolate CPV27, complete genome | 4269 | 36.3% |
| 128 | MN451657 | Canine parvovirus isolate CPV29, complete genome | 4269 | 36.4% |
| 129 | MN451658 | Canine parvovirus isolate CPV30, complete genome | 4269 | 36.3% |
| 130 | MN451659 | Canine parvovirus isolate CPV31, complete genome | 4269 | 36.2% |
| 131 | MN451660 | Canine parvovirus isolate CPV33, complete genome | 4269 | 36.3% |
| 132 | MN451661 | Canine parvovirus isolate CPV35, complete genome | 4269 | 36.3% |
| 133 | MN451662 | Canine parvovirus isolate CPV36, complete genome | 4269 | 36.4% |
| 134 | MN451663 | Canine parvovirus isolate CPV39, complete genome | 4269 | 36.3% |
| 135 | MN451664 | Canine parvovirus isolate CPV48, complete genome | 4269 | 36.3% |
| 136 | MN451665 | Canine parvovirus isolate CPV50, complete genome | 4269 | 36.3% |
| 137 | MN451666 | Canine parvovirus isolate CPV54, complete genome | 4269 | 36.3% |
| 138 | MN451667 | Canine parvovirus isolate CPV58, complete genome | 4269 | 36.3% |
| 139 | MN451668 | Canine parvovirus isolate CPV63, complete genome | 4269 | 36.2% |
| 140 | MN451669 | Canine parvovirus isolate CPV81, complete genome | 4269 | 36.3% |
| 141 | MN451670 | Canine parvovirus isolate CPV87, complete genome | 4269 | 36.3% |
| 142 | MN451671 | Canine parvovirus isolate CPV220, complete genome | 4269 | 36.2% |
| 143 | MN451672 | Canine parvovirus isolate CPV305, complete genome | 4269 | 36.2% |
| 144 | MN451673 | Canine parvovirus isolate CPV307, complete genome | 4269 | 36.3% |
| 145 | MN451674 | Canine parvovirus isolate CPV353, complete genome | 4269 | 36.5% |
| 146 | MN451675 | Canine parvovirus isolate CPV601, complete genome | 4269 | 36.4% |
| 147 | MN451676 | Canine parvovirus isolate CPV603, complete genome | 4269 | 36.5% |
| 148 | MN451677 | Canine parvovirus isolate CPV604, complete genome | 4269 | 36.6% |
| 149 | MN451678 | Canine parvovirus isolate CPV605, complete genome | 4269 | 36.5% |
| 150 | MN451679 | Canine parvovirus isolate CPV606, complete genome | 4269 | 36.6% |
| 151 | MN451680 | Canine parvovirus isolate CPV607, complete genome | 4269 | 36.4% |
| 152 | MN451681 | Canine parvovirus isolate CPV608, complete genome | 4269 | 36.4% |
| 153 | MN451682 | Canine parvovirus isolate CPV609, complete genome | 4269 | 36.5% |
| 154 | MN451683 | Canine parvovirus isolate CPV610, complete genome | 4269 | 36.6% |
| 155 | MN451684 | Canine parvovirus isolate CPV611, complete genome | 4269 | 36.5% |
| 156 | MN451685 | Canine parvovirus isolate CPV612, complete genome | 4269 | 36.5% |
| 157 | MN451686 | Canine parvovirus isolate CPV613, complete genome | 4269 | 36.5% |
| 158 | MN451687 | Canine parvovirus isolate CPV614, complete genome | 4269 | 36.5% |
| 159 | MN451688 | Canine parvovirus isolate CPV615, complete genome | 4269 | 36.6% |
| 160 | MN451689 | Canine parvovirus isolate CPV616, complete genome | 4269 | 36.6% |

| | | | | |
|---|---|---|---|---|
| 161 | MN451690 | Canine parvovirus isolate CPV617, complete genome | 4269 | 36.4% |
| 162 | MN451691 | Canine parvovirus isolate RACCPV1, complete genome | 4269 | 36.3% |
| 163 | MN451692 | Feline panleukopenia virus isolate RACFPV1, complete genome | 4269 | 36.6% |
| 164 | MN451693 | Canine parvovirus isolate RDPV122, complete genome | 4269 | 36.2% |
| 165 | MN451694 | Canine parvovirus isolate RDPV123, complete genome | 4269 | 36.2% |
| 166 | MN451695 | Canine parvovirus isolate RDPV124, complete genome | 4269 | 36.3% |
| 167 | MN661243 | Canine parvovirus strain CPV new 2a variant genomic sequence, sequence | 4495 | 36.2% |
| 168 | MN832850 | Canine parvovirus 2c isolate Taiwan/2018, complete genome | 4960 | 36.8% |
| 169 | MN840830 | Canine parvovirus isolate Canine/SH/1/2019, complete genome | 4269 | 36.5% |
| 170 | MN862741 | Canine parvovirus isolate CPV-2/American mink/MIVI-73/BC_2019, complete genome | 4450 | 35.9% |
| 171 | MN862742 | Canine parvovirus isolate CPV-2/River otter/OTVI-13/BC_2019, complete genome | 4450 | 35.8% |
| 172 | MT010564 | Canine parvovirus isolate CPV-AHhf1, complete genome | 5061 | 37.0% |
| 173 | MT394163 | Canine parvovirus isolate QD-12, complete genome | 5062 | 37.1% |
| 174 | MT394164 | Canine parvovirus isolate QD-15, complete genome | 5057 | 37.1% |
| 175 | MT394165 | Canine parvovirus isolate QD-202, complete genome | 5055 | 37.0% |
| 176 | MT394166 | Canine parvovirus isolate QD-201, complete genome | 5058 | 37.0% |
| 177 | MT394167 | Canine parvovirus isolate QD-20, complete genome | 5058 | 37.0% |
| 178 | MT441832 | Canine parvovirus strain ABT/CPV/MVC/03, partial genome | 4473 | 35.9% |
| 179 | MT629886 | Canine parvovirus 2a isolate CVASU parvovirus1, complete genome | 4840 | 36.5% |
| 180 | MT648202 | Canine parvovirus isolate CPV-AHhf7, complete genome | 5058 | 37.1% |
| 181 | MT648203 | Canine parvovirus isolate CPV-AHhf27, complete genome | 5058 | 37.0% |
| 182 | MT648204 | Canine parvovirus isolate CPV-AHhf28, complete genome | 5058 | 37.1% |
| 183 | MT648205 | Canine parvovirus isolate CPV-mas2, complete genome | 5055 | 36.9% |
| 184 | MT648206 | Canine parvovirus isolate CPV-AHmas3, complete genome | 5060 | 37.2% |
| 185 | MT648207 | Canine parvovirus isolate CPV-AHmas9, complete genome | 5061 | 36.9% |
| 186 | MT648208 | Canine parvovirus isolate CPV-AHmas16, complete genome | 5059 | 37.1% |
| 187 | MT648209 | Canine parvovirus isolate CPV-AHmas17, complete genome | 5061 | 37.1% |
| 188 | MT648210 | Canine parvovirus isolate CPV-AHcf4, complete genome | 5061 | 37.0% |
| 189 | MT892649 | Canine parvovirus isolate China-XY, complete genome | 5059 | 37.1% |
| 190 | MW539053 | Canine parvovirus 2b isolate I1, complete genome | 4873 | 36.6% |
| 191 | MW653248 | Canine parvovirus 2a strain CPV/Iran, complete genome | 5053 | 37.0% |
| 192 | MW653249 | Canine parvovirus 2a strain CPV/2/Iran, complete genome | 4668 | 36.0% |
| 193 | MW653250 | Canine parvovirus 2a strain CPV/22/Iran, complete genome | 4921 | 36.8% |
| 194 | MW653251 | Canine parvovirus 2b strain CPV/23/Iran, complete genome | 5020 | 37.0% |
| 195 | MW653252 | Canine parvovirus 2b strain CPV/30/Iran, complete genome | 4810 | 36.3% |
| 196 | MW653253 | Canine parvovirus 2c strain CPV/19/Iran, complete genome | 4269 | 36.4% |
| 197 | MW653254 | Canine parvovirus strain CPV/18/Iran, complete genome | 4468 | 36.0% |
| 198 | MW653255 | Canine parvovirus strain CPV/8/Iran, complete genome | 5323 | 35.6% |
| 199 | MW653256 | Canine parvovirus 2a strain 2a/6/Iran, complete genome | 4269 | 36.4% |
| 200 | MW811188 | Canine parvovirus 2c strain CPV-SH2002, complete genome | 4479 | 36.0% |
| 201 | MW811189 | Canine parvovirus 2c strain CPV-SH2003, complete genome | 4508 | 36.0% |
| 202 | MW815496 | Canine parvovirus strain CPV-BJ-C28, complete genome | 4698 | 36.0% |
| 203 | MW889095 | Canine parvovirus 2a strain CPV01, complete genome | 4269 | 36.5% |
| 204 | MW889096 | Canine parvovirus 2a strain CPV02, complete genome | 4269 | 36.4% |
| 205 | MW889097 | Canine parvovirus 2c strain CPV03, complete genome | 4269 | 36.5% |

| | | | | |
|---|---|---|---|---|
| 206 | MW889098 | Canine parvovirus 2a strain CPV04, complete genome | 4269 | 36.4% |
| 207 | MW889099 | Canine parvovirus 2b strain CPV05, complete genome | 4269 | 36.5% |
| 208 | MW889100 | Canine parvovirus 2a strain CPV06, complete genome | 4269 | 36.4% |
| 209 | MW889101 | Canine parvovirus 2b strain CPV07, complete genome | 4269 | 36.5% |
| 210 | MW889102 | Canine parvovirus 2b strain CPV08, complete genome | 4269 | 36.5% |
| 211 | MW889103 | Canine parvovirus 2a strain CPV09, complete genome | 4269 | 36.5% |
| 212 | MW889104 | Canine parvovirus 2a strain CPV10, complete genome | 4269 | 36.4% |
| 213 | MW889105 | Canine parvovirus 2a strain CPV11, complete genome | 4269 | 36.4% |
| 214 | MW889106 | Canine parvovirus 2c strain CPV12, complete genome | 4269 | 36.4% |
| 215 | MW889107 | Canine parvovirus 2c strain CPV13, complete genome | 4269 | 36.5% |
| 216 | MW889108 | Canine parvovirus 2c strain CPV14, complete genome | 4269 | 36.4% |
| 217 | MW889109 | Canine parvovirus 2a strain CPV15, complete genome | 4269 | 36.4% |
| 218 | MW889110 | Canine parvovirus 2c strain CPV16, complete genome | 4269 | 36.5% |
| 219 | MW889111 | Canine parvovirus 2a strain CPV17, complete genome | 4269 | 36.5% |
| 220 | MW889112 | Canine parvovirus 2c strain CPV18, complete genome | 4269 | 36.5% |
| 221 | MW889113 | Canine parvovirus 2a strain CPV19, complete genome | 4269 | 36.4% |
| 222 | MW889114 | Canine parvovirus 2c strain CPV20, complete genome | 4269 | 36.5% |
| 223 | MW889115 | Canine parvovirus 2a strain CPV21, complete genome | 4269 | 36.4% |
| 224 | MW889116 | Canine parvovirus 2c strain CPV22, complete genome | 4269 | 36.5% |
| 225 | MW889117 | Canine parvovirus 2c strain CPV23, complete genome | 4269 | 36.4% |
| 226 | MW889118 | Canine parvovirus 2b strain CPV24, complete genome | 4269 | 36.6% |
| 227 | MW889119 | Canine parvovirus 2c strain CPV25, complete genome | 4269 | 36.5% |
| 228 | MW889120 | Canine parvovirus 2c strain CPV26, complete genome | 4269 | 36.4% |
| 229 | MZ362881 | Canine parvovirus 2b isolate Emerald/Queensland/4720/2019, complete genome | 4504 | 36.0% |
| 230 | MZ362882 | Canine parvovirus 2a isolate Roseworthy/SA/5371/2019, complete genome | 4504 | 36.0% |
| 231 | MZ647470 | Canine parvovirus strain MO, complete genome | 4923 | 36.8% |
| 232 | MZ666397 | Carnivore protoparvovirus 1 strain SDS21601, complete genome | 4269 | 36.5% |
| 233 | MZ666398 | Carnivore protoparvovirus 1 strain SDS21608, complete genome | 4269 | 36.5% |
| 234 | MZ712026 | Feline panleukopenia virus isolate AMPV2020, complete genome | 5277 | 36.8% |
| 235 | NC_001539 | Canine parvovirus, complete genome | 5323 | 35.6% |
| 236 | OM640097 | Canine parvovirus 2 strain CMP10, complete genome | 4450 | 35.8% |
| 237 | OM640098 | Canine parvovirus 2 strain FM4, complete genome | 4450 | 36.0% |
| 238 | OM640099 | Canine parvovirus 2 strain KH18, complete genome | 4450 | 35.6% |
| 239 | OM640100 | Canine parvovirus 2 strain MIH9, complete genome | 4450 | 35.6% |
| 240 | OM640101 | Canine parvovirus 2 strain TN10, complete genome | 4450 | 36.0% |
| 241 | OM640102 | Canine parvovirus 2 strain LSD39, complete genome | 4450 | 35.8% |
| 242 | OM640103 | Canine parvovirus 2 strain LSD40, complete genome | 4450 | 35.8% |
| 243 | OM721655 | Canine parvovirus 2b isolate CPV-2b-K5-TR, complete genome | 4450 | 36.1% |
| 244 | OM721656 | Canine parvovirus 2b isolate CPV-2b-O1-TR, complete genome | 4452 | 36.2% |
| 245 | ON733252 | Canine parvovirus 2b strain FR1/CPV2-2021-HUN, complete genome | 5016 | 37.0% |
| 246 | OP093952 | Canine parvovirus strain CPV/PK06/2019/BRA, complete genome | 5049 | 37.1% |
| 247 | OP093953 | Canine parvovirus strain CPV/PK05/2019/BRA, complete genome | 5049 | 37.1% |
| 248 | OP093954 | Canine parvovirus strain CPV/PK04/2019/BRA, complete genome | 5049 | 37.1% |
| 249 | OP093955 | Canine parvovirus strain CPV/PK03/2019/BRA, complete genome | 5049 | 37.1% |
| 250 | OP093956 | Canine parvovirus strain CPV/PK02/2019/BRA, complete genome | 5049 | 37.1% |
