## Supplementary Table S2 for "Recovery of complete genomes of canine parvovirus from clinical samples"

**Supplementary Table S2 -** CPV whole genome sequences available in GenBank used in phylogenetic analyses.

| Access number | Isolate | Species | Country | Continent | Year of collection | Clade | Subtype |
|---|---|---|---|---|---|---|---|
| AY742935 | CPV-U6 | Canis familiaris | Germany | Europe | 1995 | I | 2a |
| D26079 | Y1 | NI | Japan | Asia | 1982 | I | 2a |
| JN033694 | Laika-1993 | Canis familiaris | Russia | Europe | 1993 | I | 2b |
| JN867614 | CPV/Raccoon/VA/278-A.us.09 | Procyon lotor | USA | North America | 2009 | I | 2a |
| JN867615 | CPV/Raccoon/GA/287.us.08 | Procyon lotor | USA | North America | 2009 | I | 2a |
| JN867616 | CPV/Raccoon/GA/289.us.08 | Procyon lotor | USA | North America | 2008 | I | 2a |
| JN867617 | CPV/Raccoon/349.us.08 | Procyon lotor | USA | North America | 2008 | I | 2a |
| JN867618 | CPV/Raccoon/WI/37.us.10 | Canis familiaris | USA | North America | 2010 | I | 2a |
| JQ268284 | CPV-LZ2 | Canis familiaris | China | Asia | 2011 | I | 2b |
| JQ686671 | JQ686671 | Canis familiaris | China | Asia | 2011 | I | 2a |
| KF676668 | CPV-JS2 | Canis familiaris | China | Asia | 2009 | I | 2a |
| KM457139 | UY364rec | Canis familiaris | Uruguay | South America | 2011 | I | 2a |
| KR002792 | CPV/CN/SH1/2013 | Canis familiaris | China | Asia | 2013 | I | 2a |
| KR002793 | CPV/CN/HB1/2013 | Canis familiaris | China | Asia | 2013 | I | 2b |
| KR002796 | CPV/CN/JL3/2013 | Canis familiaris | China | Asia | 2013 | I | 2b |
| KR002799 | CPV/CN/JL6/2013 | Canis familiaris | China | Asia | 2013 | I | 2b |
| KR002800 | CPV/CN/LN1/2014 | Canis familiaris | China | Asia | 2014 | I | 2a |
| KR002803 | CPV/CN/SD10/2014 | Canis familiaris | China | Asia | 2014 | I | 2a |
| KR002805 | CPV/CN/SD19/2014 | Canis familiaris | China | Asia | 2014 | I | 2a |
| KT382542 | CPV-SH14 | Canis familiaris | China | Asia | 2014 | I | 2a |
| KX434457 | CPV_IZSSI_987_10 | Canis familiaris | Italy | Europe | 2010 | I | 2a |
| KX618915 | 2a | Paradoxurus musangus | Singapore | Asia | 2016 | I | 2a |
| KX774249 | Bel2014-01 | Canis familiaris | Brazil | South America | 2014 | I | 2b |
| KX774250 | Bel2016-01 | Canis familiaris | Brazil | South America | 2016 | I | 2b |
| KX774251 | Bel2015-01 | Canis familiaris | Brazil | South America | 2015 | I | 2b |
| KX774252 | Bel2015-02 | Canis familiaris | Brazil | South America | 2015 | I | 2b |
| KY403998 | CPV-YH | Canis familiaris | China | Asia | 2008 | I | 2a |
| LC214969 | CPV/dog/HCM/7/2013 | Canis familiaris | Vietnam | Asia | 2013 | I | 2c |
| MF177224 | 43-97 | Canis familiaris | Italy | Europe | 1997 | I | 2a |
| MF177226 | 1-99 | Canis familiaris | Italy | Europe | 1999 | I | 2b |
| MF177231 | 260-00 | Canis familiaris | Italy | Europe | 2000 | I | 2b |
| MF177232 | 201-98 | Canis familiaris | Italy | Europe | 1998 | I | 2b |
| MF177233 | 19-99 | Canis familiaris | Italy | Europe | 1999 | I | 2a |
| MF177241 | Arg9 | Canis familiaris | Argentina | South America | 2003 | I | 2a |
| MF177246 | Arg50 | Canis familiaris | Argentina | South America | 2009 | I | 2b |
| MF177251 | BRA01/13 | Canis familiaris | Brazil | South America | 2013 | I | 2b |
| MF177256 | BRA05/13 | Canis familiaris | Brazil | South America | 2013 | I | 2b |
| MF177258 | BRA03/13 | Canis familiaris | Brazil | South America | 2013 | I | 2b |
| MF177259 | BRA02/13 | Canis familiaris | Brazil | South America | 2013 | I | 2b |
| MF177265 | E12.11 | Canis familiaris | Ecuador | South America | 2011 | I | 2a |
| MF177266 | E13.11 | Canis familiaris | Ecuador | South America | 2011 | I | 2c |
| MF177268 | E19.11 | Canis familiaris | Ecuador | South America | 2011 | I | 2a |
| MF177269 | E20.11 | Canis familiaris | Ecuador | South America | 2011 | I | 2b |

| | | | | | | | |
|---|---|---|---|---|---|---|---|
| MF177271 | E26.11 | Canis familiaris | Ecuador | South America | 2011 | I | 2c |
| MF177275 | E32.11 | Canis familiaris | Ecuador | South America | 2011 | I | 2c |
| MF177276 | E35.11 | Canis familiaris | Ecuador | South America | 2011 | I | 2a |
| MF177277 | E36.11 | Canis familiaris | Ecuador | South America | 2011 | I | 2a |
| MF177280 | E6.11 | Canis familiaris | Ecuador | South America | 2011 | I | 2b |
| MF177281 | UY6.06 | Canis familiaris | Uruguay | South America | 2006 | I | 2a |
| MF423123 | CPV/Coyote/C16/NL_2014 | Canis Latrans | Canada | North America | 2014 | I | 2b |
| MF423124 | CPV/Coyote/C55/NL_2014 | Canis Latrans | Canada | North America | 2014 | I | 2b |
| MF457594 | OH20219 | Canis familiaris | USA | North America | 2015 | I | 2c |
| MF510157 | CPV_IZSSI_2743_17 | Canis familiaris | Italy | Europe | 2017 | I | 2c |
| MF805789 | Canine/China/01/2016 | Canis familiaris | China | Asia | 2016 | I | 2c |
| MF805790 | Canine/China/02/2016 | Canis familiaris | China | Asia | 2016 | I | 2a |
| MF805791 | Canine/China/03/2016 | Canis familiaris | China | Asia | 2016 | I | 2a |
| MF805792 | Canine/China/04/2016 | Canis familiaris | China | Asia | 2016 | I | 2c |
| MF805793 | Canine/China/05/2016 | Canis familiaris | China | Asia | 2016 | I | 2a |
| MF805794 | Canine/China/06/2016 | Canis familiaris | China | Asia | 2016 | I | 2a |
| MF805795 | Canine/China/07/2016 | Canis familiaris | China | Asia | 2016 | I | 2c |
| MF805796 | Canine/China/08/2016 | Canis familiaris | China | Asia | 2016 | I | 2c |
| MF805797 | Canine/China/09/2016 | Canis familiaris | China | Asia | 2016 | I | 2c |
| MF805798 | Canine/China/10/2016 | Canis familiaris | China | Asia | 2016 | I | 2a |
| MG013488 | CPV-SH1516 | Canis familiaris | China | Asia | 2017 | I | 2c |
| MG434738 | CPV_IZSSI_PA43847/2016 | Canis familiaris | Italy | Europe | 2017 | I | 2a |
| MG434739 | CPV_IZSSI_PA48686/2016 | Canis familiaris | Italy | Europe | 2016 | I | 2a |
| MG434740 | CPV_IZSSI_PA3213/2017 | Canis familiaris | Italy | Europe | 2017 | I | 2a |
| MG434741 | CPV_IZSSI_PA5610/2017 | Canis familiaris | Italy | Europe | 2017 | I | 2a |
| MG434742 | CPV_IZSSI_PA10388/2017 | Canis familiaris | Italy | Europe | 2017 | I | 2a |
| MG434743 | CPV_IZSSI_PA13577/2017 | Canis familiaris | Italy | Europe | 2017 | I | 2a |
| MG434744 | CPV_IZSSI_PA13579id90/2017 | Canis familiaris | Italy | Europe | 2017 | I | 2a |
| MG434745 | CPV_IZSSI_PA13579id93/2017 | Canis familiaris | Italy | Europe | 2017 | I | 2a |
| MG583676 | CPV/CN/ya1/2017 | Yak | China | Asia | 2015 | I | 2a |
| MG763189 | CPV-L | Canis familiaris | China | Asia | 2014 | I | 2a |
| MH476580 | Canine/China/11/2017 | Canis familiaris | China | Asia | 2017 | I | 2a |
| MH476581 | Canine/China/12/2017 | Canis familiaris | China | Asia | 2017 | I | 2c |
| MH476583 | Canine/China/14/2017 | Canis familiaris | China | Asia | 2017 | I | 2c |
| MH476584 | Canine/China/15/2017 | Canis familiaris | China | Asia | 2017 | I | 2c |
| MH476585 | Canine/China/16/2017 | Canis familiaris | China | Asia | 2017 | I | 2c |
| MH476587 | Canine/China/18/2017 | Canis familiaris | China | Asia | 2017 | I | 2c |
| MH476589 | Canine/China/20/2017 | Canis familiaris | China | Asia | 2017 | I | 2a |
| MH476591 | Canine/China/22/2017 | Canis familiaris | China | Asia | 2017 | I | 2a |
| MH476592 | Canine/China/23/2017 | Canis familiaris | China | Asia | 2017 | I | 2c |
| MH476593 | Canine/China/24/2017 | Canis familiaris | China | Asia | 2017 | I | 2a |
| MH545963 | TN/CPV2a/2018 | Canis familiaris | India | Asia | 2018 | I | 2a |
| MH660909 | 5 MGL | Canis familiaris | Mongolia | Asia | 2017 | I | 2c |
| MH711894 | CU24 | Canis familiaris | Thailand | Asia | 2016 | I | 2c |
| MH711902 | CU21 | Felis catus | China | Asia | 2016 | I | 2c |
| MK388674 | HB2017 | Canis familiaris | China | Asia | 2017 | I | 2c |
| MK413740 | CPV-2a_PA30636/17 | Canis familiaris | Italy | Europe | 2017 | I | 2a |

| | | | | | | | |
|---|---|---|---|---|---|---|---|
| MK413741 | CPV-2a_PA31209/17 | Canis familiaris | Italy | Europe | 2017 | I | 2a |
| MK413742 | CPV-2b_PA13600/17 | Canis familiaris | Italy | Europe | 2017 | I | 2b |
| MK806279 | IZSSI_PA24478/18_id3184 | Canis familiaris | Italy | Europe | 2018 | I | 2c |
| MK806280 | IZSSI_PA24478/18_id3230 | Canis familiaris | Italy | Europe | 2018 | I | 2c |
| MK806281 | IZSSI_PA31342/18 | Canis familiaris | Italy | Europe | 2018 | I | 2c |
| MK806282 | IZSSI_PA5455/19 | Canis familiaris | Italy | Europe | 2018 | I | 2c |
| MK806283 | IZSSI_PA5446/19 | Canis familiaris | Italy | Europe | 2018 | I | 2c |
| MK806284 | IZSSI_RG3408/19 | Canis familiaris | Italy | Europe | 2019 | I | 2c |
| MK806285 | IZSSI_PA5632/19 | Canis familiaris | Italy | Europe | 2019 | I | 2c |
| MK895483 | IZSSI_PA1464/19_idUV1 | Canis familiaris | Nigeria | Africa | 2018 | I | 2a |
| MK895484 | IZSSI_PA1464/19_idYV8 | Canis familiaris | Nigeria | Africa | 2018 | I | 2a |
| MK895485 | IZSSI_PA1464/19_idUV6 | Canis familiaris | Nigeria | Africa | 2018 | I | 2a |
| MK895486 | IZSSI_PA1464/19_idYV2 | Canis familiaris | Nigeria | Africa | 2018 | I | 2c |
| MK895487 | IZSSI_PA1464/19_idEV8 | Canis familiaris | Nigeria | Africa | 2018 | I | 2c |
| MK895488 | IZSSI_PA1464/19_idJOE2 | Canis familiaris | Nigeria | Africa | 2018 | I | 2c |
| MK895489 | IZSSI_PA1464/19_idNC | Canis familiaris | Nigeria | Africa | 2018 | I | 2c |
| MK895490 | IZSSI_PA1464/19_idPSV21 | Canis familiaris | Nigeria | Africa | 2018 | I | 2c |
| MN451656 | CPV27 | Canis familiaris | USA | North America | 1983 | I | 2a |
| MN451657 | CPV29 | Canis familiaris | USA | North America | 1983 | I | 2a |
| MN451658 | CPV30 | Canis familiaris | USA | North America | 1983 | I | 2a |
| MN451659 | CPV31 | Canis familiaris | USA | North America | 1983 | I | 2a |
| MN451660 | CPV33 | Canis familiaris | USA | North America | 1983 | I | 2a |
| MN451661 | CPV35 | Canis familiaris | USA | North America | 1984 | I | 2a |
| MN451662 | CPV36 | Canis familiaris | USA | North America | 1984 | I | 2b |
| MN451663 | CPV39 | Canis familiaris | USA | North America | 1984 | I | 2b |
| MN451666 | CPV54 | Canis familiaris | France | Europe | 1984 | I | 2b |
| MN451667 | CPV58 | Canis familiaris | France | Europe | 1983 | I | 2a |
| MN451669 | CPV81 | Canis familiaris | Australia | Oceania | 1982 | I | 2a |
| MN451670 | CPV87 | Canis familiaris | Australia | Oceania | 1985 | I | 2a |
| MN451674 | CPV353 | Canis familiaris | USA | North America | 1996 | I | 2a |
| MN451676 | CPV603 | Canis familiaris | USA | North America | 2017 | I | 2c |
| MN451677 | CPV604 | Canis familiaris | USA | North America | 2007 | I | 2b |
| MN451678 | CPV605 | Canis familiaris | Nigeria | Africa | 2018 | I | 2c |
| MN451680 | CPV607 | Canis familiaris | Nigeria | Africa | 2018 | I | 2c |
| MN451681 | CPV608 | Canis familiaris | Nigeria | Africa | 2018 | I | 2c |
| MN451682 | CPV609 | Canis familiaris | Nigeria | Africa | 2018 | I | 2c |
| MN451684 | CPV611 | Canis familiaris | USA | North America | 2019 | I | 2c |
| MN451686 | CPV613 | Canis familiaris | USA | North America | 2019 | I | 2c |
| MN451689 | CPV616 | Canis familiaris | Nigeria | Africa | 2018 | I | 2a |
| MN661243 | CPV new 2a variant | Canis familiaris | India | Asia | 2016 | I | 2a |
| MN832850 | Taiwan/2018 | Pangolin | Taiwan | Asia | 2018 | I | 2c |
| MT010564 | CPV-AHhf1 | Canis familiaris | China | Asia | 2018 | I | 2c |
| OR230510 | CPV/UFT01/2022/BRA | Canis familiaris | Brazil | South America | 2022 | I | 2c |
| OR230511 | CPV/UFT02/2022/BRA | Canis familiaris | Brazil | South America | 2022 | I | 2a |
| OR230512 | CPV/UFT03/2022/BRA | Canis familiaris | Brazil | South America | 2022 | I | 2a |
| OR230513 | CPV/UFT04/2022/BRA | Canis familiaris | Brazil | South America | 2022 | I | 2c |
| OR230514 | CPV/UFT05/2022/BRA | Canis familiaris | Brazil | South America | 2022 | I | 2a |

| | | | | | | | |
|---|---|---|---|---|---|---|---|
| OR230515 | CPV/UFT06/2022/BRA | Canis familiaris | Brazil | South America | 2022 | I | 2a |
| OR230516 | CPV/UFT07/2023/BRA | Canis familiaris | Brazil | South America | 2023 | I | 2a |
| AJ564427 | CPV2a-K_India | Canis familiaris | India | Asia | 1999 | II | 2a |
| AY742932 | CPV2b-193.us.91 | Canis familiaris | USA | North America | 1991 | II | 2b |
| AY742933 | CPV-339 | Canis familiaris | New Zeland | Oceania | 1994 | II | 2a |
| AY742934 | CPV-447 | Canis familiaris | Germany | Europe | 1995 | II | 2b |
| AY742936 | CPV2b-395.us.98 | Canis familiaris | USA | North America | 1998 | II | 2b |
| EF011664 | B-2004 | Canis familiaris | China | Asia | 2004 | II | 2a |
| EU310373 | CPV2a/nj01/06.ch.06 | Canis familiaris | China | Asia | 2007 | II | 2a |
| EU659118 | CPV2a-13.us.81 | Canis familiaris | USA | North America | 1981 | II | 2a |
| EU659119 | CPV2b-410.us.00 | Canis familiaris | USA | North America | 2000 | II | 2b |
| EU659120 | CPV2b-411a.us.98 | Canis familiaris | USA | North America | 1998 | II | 2b |
| EU659121 | CPV2b-411b.us.98 | Canis familiaris | USA | North America | 1998 | II | 2b |
| JN867610 | CPV/Raccoon/VA/118-A.us.07 | Procyon lotor | USA | North America | 2007 | II | 2a |
| JN867611 | CPV/Raccoon/KY/358-B.us.09 | Procyon lotor | USA | North America | 2009 | II | 2a |
| JN867612 | CPV/Raccoon/TN/351.us.09 | Procyon lotor | USA | North America | 2009 | II | 2a |
| JN867613 | CPV/Raccoon/FL/381.us.09 | Procyon lotor | USA | North America | 2009 | II | 2a |
| JQ268283 | CPV-LZ1 | Canis familiaris | China | Asia | 2011 | II | 2a |
| JX660690 | SC02/2011 | Canis familiaris | China | Asia | 2011 | II | 2a |
| KF366250 | CPV/915-H | Canis familiaris | India | Asia | 2013 | II | 2a |
| KF638400 | s5 | Canis familiaris | China | Asia | 2010 | II | 2a |
| KM457102 | UY243.10 | Canis familiaris | Uruguay | South America | 2010 | II | 2a |
| KM457103 | UY12.06 | Canis familiaris | Uruguay | South America | 2006 | II | 2c |
| KM457104 | UY47.06 | Canis familiaris | Uruguay | South America | 2006 | II | 2c |
| KM457105 | UY52.06 | Canis familiaris | Uruguay | South America | 2006 | II | 2c |
| KM457106 | UY55.06 | Canis familiaris | Uruguay | South America | 2006 | II | 2c |
| KM457107 | UY72.07 | Canis familiaris | Uruguay | South America | 2007 | II | 2c |
| KM457108 | UY82.07 | Canis familiaris | Uruguay | South America | 2007 | II | 2c |
| KM457109 | UY95.07 | Canis familiaris | Uruguay | South America | 2007 | II | 2c |
| KM457110 | UY101.07 | Canis familiaris | Uruguay | South America | 2007 | II | 2c |
| KM457111 | UY120.08 | Canis familiaris | Uruguay | South America | 2008 | II | 2c |
| KM457112 | UY135.08 | Canis familiaris | Uruguay | South America | 2008 | II | 2c |
| KM457113 | UY152.08 | Canis familiaris | Uruguay | South America | 2008 | II | 2c |
| KM457114 | UY169.08 | Canis familiaris | Uruguay | South America | 2008 | II | 2c |
| KM457115 | UY173.09 | Canis familiaris | Uruguay | South America | 2009 | II | 2c |
| KM457116 | UY185.09 | Canis familiaris | Uruguay | South America | 2009 | II | 2c |
| KM457117 | UY187.09 | Canis familiaris | Uruguay | South America | 2009 | II | 2c |
| KM457118 | UY190.09 | Canis familiaris | Uruguay | South America | 2009 | II | 2c |
| KM457119 | UY235.10 | Canis familiaris | Uruguay | South America | 2010 | II | 2c |
| KM457120 | UY242.10 | Canis familiaris | Uruguay | South America | 2010 | II | 2c |
| KM457121 | UY247.10 | Canis familiaris | Uruguay | South America | 2010 | II | 2c |
| KM457122 | UY258.10 | Canis familiaris | Uruguay | South America | 2010 | II | 2c |
| KM457123 | UY261.10 | Canis familiaris | Uruguay | South America | 2010 | II | 2c |
| KM457124 | UY307.11 | Canis familiaris | Uruguay | South America | 2011 | II | 2c |
| KM457125 | UY317.11 | Canis familiaris | Uruguay | South America | 2011 | II | 2c |
| KM457126 | UY318.10 | Canis familiaris | Uruguay | South America | 2010 | II | 2c |

| | | | | | | | |
|---|---|---|---|---|---|---|---|
| KM457127 | UY326.11 | Canis familiaris | Uruguay | South America | 2011 | II | 2c |
| KM457128 | UY346.11 | Canis familiaris | Uruguay | South America | 2011 | II | 2c |
| KM457129 | UY349.11 | Canis familiaris | Uruguay | South America | 2011 | II | 2c |
| KM457130 | UY354.11 | Canis familiaris | Uruguay | South America | 2011 | II | 2c |
| KM457131 | UY368.11 | Canis familiaris | Uruguay | South America | 2011 | II | 2c |
| KM457132 | UY245.10 | Canis familiaris | Uruguay | South America | 2010 | II | 2a |
| KM457133 | UY250.10 | Canis familiaris | Uruguay | South America | 2010 | II | 2a |
| KM457134 | UY280.10 | Canis familiaris | Uruguay | South America | 2010 | II | 2a |
| KM457135 | UY306.11 | Canis familiaris | Uruguay | South America | 2011 | II | 2a |
| KM457136 | UY315.11 | Canis familiaris | Uruguay | South America | 2011 | II | 2a |
| KM457137 | UY344.11 | Canis familiaris | Uruguay | South America | 2011 | II | 2a |
| KM457138 | UY363.11 | Canis familiaris | Uruguay | South America | 2011 | II | 2a |
| KM457140 | UY365.11 | Canis familiaris | Uruguay | South America | 2011 | II | 2a |
| KM457141 | UY370a | Canis familiaris | Uruguay | South America | 2011 | II | 2a |
| KM457142 | UY370c | Canis familiaris | Uruguay | South America | 2011 | II | 2c |
| KM457143 | UY364.11 | Canis familiaris | Uruguay | South America | 2011 | II | 2a |
| KR002794 | CPV/CN/HB3/2013 | Canis familiaris | China | Asia | 2013 | II | 2a |
| KR002795 | CPV/CN/JL1/2013 | Canis familiaris | China | Asia | 2013 | II | 2a |
| KR002797 | CPV/CN/JL4/2013 | Canis familiaris | China | Asia | 2013 | II | 2a |
| KR002798 | CPV/CN/JL5/2013 | Canis familiaris | China | Asia | 2013 | II | 2a |
| KR002801 | CPV/CN/SD6/2014 | Canis familiaris | China | Asia | 2014 | II | 2a |
| KR002802 | CPV/CN/SD9/2014 | Canis familiaris | China | Asia | 2014 | II | 2a |
| KR002804 | CPV/CN/SD18/2014 | Canis familiaris | China | Asia | 2014 | II | 2a |
| KU508407 | CPV_IZSSI_25835_09 | Canis familiaris | Italy | Europe | 2009 | II | 2c |
| KU508691 | HB | Canis familiaris | Australia | Oceania | 2015 | II | 2c |
| KU508692 | FH | Canis familiaris | Australia | Oceania | 2015 | II | 2c |
| KU508693 | LW | Canis familiaris | Australia | Oceania | 2015 | II | 2c |
| KX434454 | CPV_IZSSI_29451_09 | Canis familiaris | Italy | Europe | 2009 | II | 2a |
| KX434455 | CPV_IZSSI_23782_09 | Canis familiaris | Italy | Europe | 2009 | II | 2c |
| KX434456 | CPV_IZSSI_45361_09 | Canis familiaris | Italy | Europe | 2009 | II | 2c |
| KX434458 | CPV_IZSSI_2323_11 | Canis familiaris | Italy | Europe | 2011 | II | 2c |
| KX434459 | CPV_IZSSI_27692_1_11 | Canis familiaris | Italy | Europe | 2011 | II | 2c |
| KX434460 | CPV_IZSSI_52238_12 | Canis familiaris | Italy | Europe | 2012 | II | 2c |
| KY073269 | UFMT | Canis familiaris | Brazil | South America | 2015 | II | 2c |
| LC214970 | CPV/dog/HCM/22/2013 | Canis familiaris | Vietnam | Asia | 2012 | II | 2a |
| LC270891 | 9985 | Canis familiaris | Japan | Asia | 2017 | II | 2b |
| LC270892 | 9985-46 | Canis familiaris | Japan | Asia | 2017 | II | 2b |
| MF069442 | CPV/Raccoon/RC14/BC_2010 | Procyon Lotor | Canada | North America | 2010 | II | 2a |
| MF069443 | CPV/Raccoon/RC19/BC_2016 | Procyon Lotor | Canada | North America | 2016 | II | 2a |
| MF069444 | CPV/Raccoon/RC20/BC_2016 | Procyon Lotor | Canada | North America | 2016 | II | 2a |
| MF134808 | CPV-BJ03/17 | Canis familiaris | China | Asia | 2017 | II | 2a |
| MF177225 | 242-98 | Canis familiaris | Italy | Europe | 1998 | II | 2b |
| MF177227 | 202-09 | Canis familiaris | France | Europe | 2009 | II | 2c |
| MF177228 | 485-09 | Canis familiaris | Italy | Europe | 2009 | II | 2c |
| MF177229 | 368-12-17 | Canis familiaris | Albania | Europe | 2012 | II | 2c |
| MF177230 | 57-10 | Canis familiaris | Italy | Europe | 2010 | II | 2c |
| MF177234 | 347-03 | Canis familiaris | Italy | Europe | 2003 | II | 2c |

| | | | | | | | |
|---|---|---|---|---|---|---|---|
| MF177235 | 283-06 | Canis familiaris | Italy | Europe | 2006 | II | 2c |
| MF177236 | 244-04 | Canis familiaris | Italy | Europe | 2004 | II | 2c |
| MF177237 | 188-07 | Canis familiaris | Italy | Europe | 2007 | II | 2c |
| MF177238 | 114-05 | Canis familiaris | Italy | Europe | 2005 | II | 2c |
| MF177239 | 288-01 | Canis familiaris | Italy | Europe | 2001 | II | 2c |
| MF177240 | 189-02 | Canis familiaris | Italy | Europe | 2002 | II | 2c |
| MF177242 | Arg26 | Canis familiaris | Argentina | South America | 2008 | II | 2c |
| MF177243 | Arg32 | Canis familiaris | Argentina | South America | 2008 | II | 2c |
| MF177244 | Arg33 | Canis familiaris | Argentina | South America | 2008 | II | 2c |
| MF177245 | Arg35 | Canis familiaris | Argentina | South America | 2008 | II | 2c |
| MF177247 | Arg66 | Canis familiaris | Argentina | South America | 2010 | II | 2c |
| MF177248 | Arg68 | Canis familiaris | Argentina | South America | 2010 | II | 2c |
| MF177249 | Arg71 | Canis familiaris | Argentina | South America | 2010 | II | 2c |
| MF177250 | BRA01/10 | Canis familiaris | Brazil | South America | 2010 | II | 2c |
| MF177252 | BRA02/10 | Canis familiaris | Brazil | South America | 2010 | II | 2c |
| MF177253 | BRA03/10 | Canis familiaris | Brazil | South America | 2010 | II | 2c |
| MF177254 | BRA04/10 | Canis familiaris | Brazil | South America | 2010 | II | 2c |
| MF177255 | BRA01/14 | Canis familiaris | Brazil | South America | 2014 | II | 2c |
| MF177257 | BRA02/14 | Canis familiaris | Brazil | South America | 2014 | II | 2c |
| MF177260 | BRA01/12 | Canis familiaris | Brazil | South America | 2012 | II | 2c |
| MF177261 | BRA04/13 | Canis familiaris | Brazil | South America | 2013 | II | 2c |
| MF177262 | Py | Canis familiaris | Paraguay | South America | 2009 | II | 2c |
| MF177263 | E1.11 | Canis familiaris | Ecuador | South America | 2011 | II | 2c |
| MF177264 | E10.11 | Canis familiaris | Ecuador | South America | 2011 | II | 2c |
| MF177267 | E16.11 | Canis familiaris | Ecuador | South America | 2011 | II | 2c |
| MF177270 | E23.11 | Canis familiaris | Ecuador | South America | 2011 | II | 2c |
| MF177272 | E28.11 | Canis familiaris | Ecuador | South America | 2011 | II | 2c |
| MF177273 | E29.11 | Canis familiaris | Ecuador | South America | 2011 | II | 2c |
| MF177274 | E31.11 | Canis familiaris | Ecuador | South America | 2011 | II | 2c |
| MF177278 | E4.11 | Canis familiaris | Ecuador | South America | 2011 | II | 2c |
| MF177279 | E48.11 | Canis familiaris | Ecuador | South America | 2011 | II | 2c |
| MF177282 | UY21.06 | Canis familiaris | Uruguay | South America | 2006 | II | 2c |
| MF177283 | UY181.09 | Canis familiaris | Uruguay | South America | 2009 | II | 2c |
| MF177284 | UY196.09 | Canis familiaris | Uruguay | South America | 2009 | II | 2c |
| MF177285 | UY269.10 | Canis familiaris | Uruguay | South America | 2010 | II | 2c |
| MF177286 | UY375.11 | Canis familiaris | Uruguay | South America | 2011 | II | 2c |
| MF423125 | CPV/Coyote/C67/NL_2014 | Canis Latrans | Canada | North America | 2014 | II | 2a |
| MF510158 | CPV_IZSSI_41113_c1_16 | Canis familiaris | Italy | Europe | 2016 | II | 2c |
| MH106698 | CPV-BJL1 | Canis familiaris | China | Asia | 2015 | II | 2a |
| MH106699 | CPV-BJL2 | Canis familiaris | China | Asia | 2015 | II | 2b |
| MH106700 | CPV-BJL3 | Canis familiaris | China | Asia | 2016 | II | 2c |
| MH476582 | Canine/China/13/2017 | Canis familiaris | China | Asia | 2017 | II | 2a |
| MH476586 | Canine/China/17/2017 | Canis familiaris | China | Asia | 2017 | II | 2a |
| MH476588 | Canine/China/19/2017 | Canis familiaris | China | Asia | 2017 | II | 2b |
| MH476590 | Canine/China/21/2017 | Canis familiaris | China | Asia | 2017 | II | 2a |
| MK413743 | CPV-2c_PA15423/16 | Felis catus | Italy | Europe | 2016 | II | 2c |
| MK413744 | CPV-2c_PA36395/16 | Canis familiaris | Italy | Europe | 2016 | II | 2c |

| | | | | | | | |
|---|---|---|---|---|---|---|---|
| MK413746 | CPV-2c_PA41113c2/16 | Canis familiaris | Italy | Europe | 2016 | II | 2c |
| MK413747 | CPV-2c_PA45984/16 | Canis familiaris | Italy | Europe | 2016 | II | 2c |
| MK413748 | CPV-2c_CT1839id0018/17 | Canis familiaris | Italy | Europe | 2017 | II | 2c |
| MK413749 | CPV-2c_CT1839id2213/17 | Canis familiaris | Italy | Europe | 2017 | II | 2c |
| MK413750 | CPV-2c_PA27184/17 | Canis familiaris | Italy | Europe | 2017 | II | 2c |
| MN451672 | CPV305 | Canis familiaris | USA | North America | 1993 | II | 2b |
| MN451673 | CPV307 | Canis familiaris | USA | North America | 1993 | II | 2a |
| MN451675 | CPV601 | Canis familiaris | USA | North America | 2016 | II | 2c |
| MN451679 | CPV606 | Canis familiaris | USA | North America | 2014 | II | 2c |
| MN451685 | CPV612 | Canis familiaris | USA | North America | 2019 | II | 2c |
| MN451690 | CPV617 | Canis familiaris | USA | North America | 2011 | II | 2c |
| MN451691 | RACCPV1 | Canis familiaris | USA | North America | 2009 | II | 2a |
| MN862741 | CPV-2/American mink/MIVI-73/BC_2019 | Neovison vison | Canada | North America | 2019 | II | 2a |
| MN862742 | CPV-2/River otter/OTVI-13/BC_2019 | Lontra Canadensis | Canada | North America | 2019 | II | 2a |
| EU659116 | CPV2-5.us.79 | Canis familiaris | USA | North America | 1978 | O | 2 |
| EU659117 | CPV2-6.us.80 | Canis familiaris | USA | North America | 1980 | O | 2 |
| M19296 | CPV2-N.us.78 | Canis familiaris | USA | North America | 1978 | O | 2 |
| M38245 | CPV2-b.us.79 | NI | USA | North America | 1979 | O | 2 |
| MN451653 | CPV6 | Canis familiaris | USA | North America | 1979 | O | 2 |
| MN451654 | CPV9 | Canis familiaris | USA | North America | 1979 | O | 2 |
| MN451655 | CPV12 | Canis familiaris | USA | North America | 1978 | O | 2 |
| MN451664 | CPV48 | Canis familiaris | USA | North America | 1985 | O | 2 |
| MN451665 | CPV50 | Canis familiaris | USA | North America | 1986 | O | 2 |
| MN451668 | CPV63 | Canis familiaris | USA | North America | 1980 | O | 2 |
| MN451671 | CPV220 | Canis familiaris | USA | North America | 1993 | O | 2 |
| MN451693 | RDPV122 | Nyctereutes procyonoides | Finland | Europe | 1980 | O | 2 |
| MN451694 | RDPV123 | Nyctereutes procyonoides | Finland | Europe | 1980 | O | 2 |
| MN451695 | RDPV124 | Nyctereutes procyonoides | Finland | Europe | 1986 | O | 2 |
