## Supplementary Table S3 for "Recovery of complete genomes of canine parvovirus from clinical samples"

**Supplementary Table S3 -** CPV VP2 nucleotide sequence available in GenBank used in phylogenetic analyses.

| Acces number | Isolate | Species | Country | Continent | Year of collection | Clade | Subtype |
|---|---|---|---|---|---|---|---|
| EU659116 | CPV2-5.us.79 | Canis familiaris | USA | North America | 1978 | O | 2 |
| EU659117 | CPV2-6.us.80 | Canis familiaris | USA | North America | 1980 | O | 2 |
| FJ222824 | 388/05-3 | Canis familiaris | Italy | Europe | 2005 | O | 2 |
| JN867618 | CPV/Raccoon/WI/37.us.10 | Canis familiaris | USA | North America | 2010 | O | 2a |
| M19296 | CPV2-N.us.78 | Canis familiaris | USA | North America | 1978 | O | 2 |
| M38245 | CPV2-b.us.79 | NI | USA | North America | 1979 | O | 2 |
| MG264079 | EC/C3/2017 | Canis familiaris | Ecuador | South America | 2017 | O | 2 |
| MN451653 | CPV6 | Canis familiaris | USA | North America | 1979 | O | 2 |
| MN451654 | CPV9 | Canis familiaris | USA | North America | 1979 | O | 2 |
| MN451655 | CPV12 | Canis familiaris | USA | North America | 1978 | O | 2 |
| MN451664 | CPV48 | Canis familiaris | USA | North America | 1985 | O | 2 |
| MN451665 | CPV50 | Canis familiaris | USA | North America | 1986 | O | 2 |
| MN451668 | CPV63 | Canis familiaris | USA | North America | 1980 | O | 2 |
| MN451671 | CPV220 | Canis familiaris | USA | North America | 1993 | O | 2 |
| MN451693 | RDPV122 | Nyctereutes procyonoides | Finland | Europe | 1980 | O | 2 |
| MN451694 | RDPV123 | Nyctereutes procyonoides | Finland | Europe | 1980 | O | 2 |
| MN451695 | RDPV124 | Nyctereutes procyonoides | Finland | Europe | 1986 | O | 2 |
| MW934264 | 16-UCS | Canis familiaris | Brazil | South America | 2019 | O | 2B |
| D26079 | Y1 | NI | Japan | Asia | 1982 | W | 2a |
| DQ340404 | BR6-80 | Canis familiaris | Brazil | South America | 1980 | W | 2a |
| DQ340405 | BR135-80 | Canis familiaris | Brazil | South America | 1980 | W | 2a |
| DQ340406 | BR137-80 | Canisfamiliaris | Brazil | South America | 1980 | W | 2a |
| DQ340407 | BR145-80 | Canis familiaris | Brazil | South America | 1980 | W | 2a |
| DQ340408 | BR154-80 | Canis familiaris | Brazil | South America | 1980 | W | 2a |
| DQ340409 | BR183-85 | Canis familiaris | Brazil | South America | 1985 | W | 2b |
| DQ340410 | BR315-86 | Canis familiaris | Brazil | South America | 1986 | W | 2a |
| EU659118 | CPV2a-13.us.81 | Canis familiaris | USA | North America | 1981 | W | 2a |
| FJ005263 | 42/05-49 | Canis familiaris | Italy | Europe | 2005 | W | 2b |
| FJ005265 | 140/05 | Canis familiaris | Italy | Europe | 2005 | W | 2b |
| GU569948 | CC8601 | Canis familiaris | China | Asia | 1986 | W | 2a |
| KU662349 | W33/PT/96 | Canis lupus | Portugal | Europe | 1996 | W | 2b |

| | | | | | | | |
|---|---|---|---|---|---|---|---|
| MF177226 | 1-99 | Canis familiaris | Italy | Europe | 1999 | W | 2b |
| MG763189 | CPV-L | Canis familiaris | China | Asia | 2014 | W | 2a |
| MN451656 | CPV27 | Canis familiaris | USA | North America | 1983 | W | 2a |
| MN451657 | CPV29 | Canisfamiliaris | USA | North America | 1983 | W | 2a |
| MN451658 | CPV30 | Canis familiaris | USA | North America | 1983 | W | 2a |
| MN451659 | CPV31 | Canisfamiliaris | USA | North America | 1983 | W | 2a |
| MN451660 | CPV33 | Canisfamiliaris | USA | North America | 1983 | W | 2a |
| MN451661 | CPV35 | Canis familiaris | USA | North America | 1984 | W | 2a |
| MN451662 | CPV36 | Canis familiaris | USA | North America | 1984 | W | 2b |
| MN451663 | CPV39 | Canis familiaris | USA | North America | 1984 | W | 2b |
| MN451666 | CPV54 | Canis familiaris | France | Europe | 1984 | W | 2b |
| MN451667 | CPV58 | Canis familiaris | France | Europe | 1983 | W | 2a |
| MN451669 | CPV81 | Canis familiaris | Australia | Oceania | 1982 | W | 2a |
| MN451670 | CPV87 | Canis familiaris | Australia | Oceania | 1985 | W | 2a |
| MN451672 | CPV305 | Canis familiaris | USA | North America | 1993 | W | 2b |
| AJ564427 | CPV2a-K_India | Canis familiaris | India | Asia | 1999 | W.1 | 2a |
| AY742933 | CPV-339 | Canis familiaris | New Zeland | Oceania | 1994 | W.1 | 2a |
| D78585 | FPV-314 | Felis catus | Japan | Asia | 1995 | W.1 | 2a |
| DQ025950 | 02B9 | Canis familiaris | France | Europe | 2005 | W.1 | 2a |
| DQ354068 | RPPV | Ailurus fulgens | China | Asia | 2004 | W.1 | 2a |
| EF011664 | B-2004 | Canis familiaris | China | Asia | 2004 | W.1 | 2a |
| EF592511 | TWN1 | Canis familiaris | Taiwan | Asia | 2006 | W.1 | 2b |
| EF599096 | DH426 | Canis familiaris | South Korea | Asia | 2007 | W.1 | 2a |
| EF666059 | CPV/BJ004/07 | Canis familiaris | China | Asia | 2007 | W.1 | 2a |
| EF666060 | CPV/BJ005/07 | Canis familiaris | China | Asia | 2007 | W.1 | 2a |
| EF666061 | CPV/BJ008/07 | Canis familiaris | China | Asia | 2007 | W.1 | 2a |
| EF666062 | CPV/BJ010/07 | Canis familiaris | China | Asia | 2007 | W.1 | 2a |
| EF666066 | CPV/BJ018/07 | Canis familiaris | China | Asia | 2017 | W.1 | 2a |
| EU009201 | K014 | Canis familiaris | South Korea | Asia | 2007 | W.1 | 2a |
| EU009203 | K022 | Canis familiaris | South Korea | Asia | 2007 | W.1 | 2a |
| EU145953 | CPV/BJ034/O7 | Canis familiaris | China | Asia | 2007 | W.1 | 2a |
| EU145955 | CPV/BJ050/O7 | Canis familiaris | China | Asia | 2007 | W.1 | 2a |
| EU145956 | CPV/BJ064/O7 | Canis familiaris | China | Asia | 2007 | W.1 | 2a |

| | | | | | | | |
|---|---|---|---|---|---|---|---|
| EU145958 | CPV/BJ069/O7 | Canis familiaris | China | Asia | 2007 | W.1 | 2a |
| EU170352 | CPV-SHZ | Canis familiaris | China | Asia | 2007 | W.1 | 2a |
| EU213073 | CPV-HZ0761 | Canis familiaris | China | Asia | 2007 | W.1 | 2a |
| EU213074 | CPV-APD1 | Canis familiaris | China | Asia | 2007 | W.1 | 2a |
| EU213077 | CPV-HT | Canis familiaris | China | Asia | 2007 | W.1 | 2a |
| EU213078 | CPV-ZD1 | Canis familiaris | China | Asia | 2007 | W.1 | 2b |
| EU213081 | CPV-BD4 | Canis familiaris | China | Asia | 2007 | W.1 | 2a |
| EU213083 | CPV-KT2 | Canis familiaris | China | Asia | 2007 | W.1 | 2a |
| EU213085 | CPV-ZD3 | Canis familiaris | China | Asia | 2007 | W.1 | 2a |
| EU310373 | CPV2a/nj01/06.ch.06 | Canis familiaris | China | Asia | 2007 | W.1 | 2a |
| EU483509 | CPV-JB2 | Canis familiaris | China | Asia | 2008 | W.1 | 2a |
| EU483511 | CPV-ZD4 | Canis familiaris | China | Asia | 2008 | W.1 | 2a |
| EU483513 | CPV-ZD7 | Canis familiaris | China | Asia | 2008 | W.1 | 2a |
| EU483514 | CPV-ZD11 | Canis familiaris | China | Asia | 2008 | W.1 | 2a |
| EU483515 | CPV-ZD13 | Canis familiaris | China | Asia | 2008 | W.1 | 2b |
| EU483516 | CPV-ZD30 | Canis familiaris | China | Asia | 2008 | W.1 | 2a |
| EU483517 | CPV-ZD35 | Canis familiaris | China | Asia | 2008 | W.1 | 2b |
| FJ005254 | 331/05 | Canis familiaris | Italy | Europe | 2005 | W.1 | 2a |
| FJ005258 | 80/08 | Canis familiaris | Italy | Europe | 2008 | W.1 | 2a |
| FJ197823 | CPVK1 | Canis familiaris | South Korea | Asia | 2007 | W.1 | 2a |
| FJ197828 | CPVK6 | Canis familiaris | South Korea | Asia | 2007 | W.1 | 2a |
| FJ197834 | CPVK12 | Canis familiaris | South Korea | Asia | 2007 | W.1 | 2a |
| FJ197835 | CPVK13 | Canis familiaris | South Korea | Asia | 2007 | W.1 | 2a |
| FJ197841 | CPVK19 | Canis familiaris | South Korea | Asia | 2007 | W.1 | 2a |
| FJ265775 | CPV301/TW04 | Canis familiaris | Taiwan | Asia | 2008 | W.1 | 2b |
| FJ265781 | CPV307/TW05 | Canis familiaris | Taiwan | Asia | 2008 | W.1 | 2b |
| FJ432717 | 08-5-WH | Canis familiaris | China | Asia | 2008 | W.1 | 2a |
| FJ435345 | CPV-04/08/CN-4 | Canis familiaris | China | Asia | 2008 | W.1 | 2a |
| FJ869126 | KU5_08 | Canis familiaris | Thailand | Asia | 2008 | W.1 | 2a |
| FJ869134 | KU23_03 | Canis familiaris | Thailand | Asia | 2003 | W.1 | 2a |
| FJ869137 | KU52_03 | Canis familiaris | Thailand | Asia | 2003 | W.1 | 2a |
| GQ169537 | wh-1 | Canis familiaris | China | Asia | 2009 | W.1 | 2a |
| GQ169539 | wh-3 | Canis familiaris | China | Asia | 2009 | W.1 | 2a |

| | | | | | | | |
|---|---|---|---|---|---|---|---|
| GQ169540 | wh-4 | Canis familiaris | China | Asia | 2007 | W.1 | 2a |
| GQ169541 | wh-5 | Canis familiaris | China | Asia | 2009 | W.1 | 2a |
| GQ169542 | wh-6 | Canis familiaris | China | Asia | 2009 | W.1 | 2a |
| GQ169545 | bj-1 | Canis familiaris | China | Asia | 2010 | W.1 | 2a |
| GQ169546 | bj-2 | Canis familiaris | China | Asia | 2010 | W.1 | 2a |
| GQ169549 | bj-5 | Canis familiaris | China | Asia | 2010 | W.1 | 2a |
| GQ379042 | KU13_08 | Canis familiaris | Thailand | Asia | 2008 | W.1 | 2a |
| GQ379043 | KU14_08 | Canis familiaris | Thailand | Asia | 2008 | W.1 | 2a |
| GQ379044 | KU143_09 | Canis familiaris | Thailand | Asia | 2009 | W.1 | 2a |
| GQ379047 | KU616_09 | Canis familiaris | Thailand | Asia | 2009 | W.1 | 2a |
| GQ379048 | KU739_09 | Canis familiaris | Thailand | Asia | 2009 | W.1 | 2a |
| GQ379049 | KU18_08 | Canis familiaris | Thailand | Asia | 2008 | W.1 | 2a |
| GQ857596 | CPV05-01 | Canis familiaris | China | Asia | 2005 | W.1 | 2a |
| GQ857600 | CPV06-01 | Canis familiaris | China | Asia | 2006 | W.1 | 2a |
| GU380303 | 06/09 | Canis familiaris | China | Asia | 2009 | W.1 | 2c |
| GU380305 | 08/09 | Canis familiaris | China | Asia | 2008 | W.1 | 2c |
| GU452715 | 11/09 | Canis familiaris | China | Asia | 2009 | W.1 | 2a |
| GU569942 | JL0202 | Canis familiaris | China | Asia | 2002 | W.1 | 2a |
| GU569946 | JL0201 | Canis familiaris | China | Asia | 2001 | W.1 | 2a |
| HQ602975 | 101-10SA | Canis familiaris | South Africa | Africa | 2010 | W.1 | 2a |
| JF767494 | S7 | Canis familiaris | China | Asia | 2009 | W.1 | 2a |
| JF789638 | BD-1010 | Canis familiaris | China | Asia | 2010 | W.1 | 2a |
| JN403045 | Shanaxi | Canis familiaris | China | Asia | 2011 | W.1 | 2a |
| JN867613 | CPV/Raccoon/FL/381.us.09 | Procyon lotor | USA | North America | 2009 | W.1 | 2a |
| JQ268283 | CPV-LZ1 | Canis familiaris | China | Asia | 2011 | W.1 | 2a |
| JQ268284 | CPV-LZ2 | Canis familiaris | China | Asia | 2011 | W.1 | 2b |
| JQ686671 | JQ686671 | Canis familiaris | China | Asia | 2011 | W.1 | 2a |
| JQ743891 | CPV-10(10) | Canis familiaris | China | Asia | 2010 | W.1 | 2b |
| JQ743898 | CPV-SS3(11) | Canis familiaris | China | Asia | 2011 | W.1 | 2a |
| JQ743900 | CPV-LBLD2(11) | Canis familiaris | China | Asia | 2011 | W.1 | 2a |
| JQ743902 | CPV-13(10) | Canis familiaris | China | Asia | 2010 | W.1 | 2a |
| JQ743903 | CPV-7(10) | Canis familiaris | China | Asia | 2010 | W.1 | 2a |
| JQ743905 | CPV-5(10) | Canis familiaris | China | Asia | 2010 | W.1 | 2a |

| | | | | | | | |
|---|---|---|---|---|---|---|---|
| JX048605 | CPV-42 | Canis familiaris | Taiwan | Asia | 2011 | W.1 | 2a |
| JX048606 | CPV-87 | Canis familiaris | Taiwan | Asia | 2011 | W.1 | 2a |
| JX120178 | CPV-GZ | Canis familiaris | China | Asia | 2010 | W.1 | 2a |
| JX121627 | CPVSH-3/2011 | Canis familiaris | China | Asia | 2011 | W.1 | 2a |
| JX660690 | SC02/2011 | Canis familiaris | China | Asia | 2011 | W.1 | 2a |
| KF366250 | CPV/915-H | Canis familiaris | India | Asia | 2013 | W.1 | 2a |
| KF482472 | S3 | Canis familiaris | China | Asia | 2009 | W.1 | 2a |
| KF482476 | S9 | Canis familiaris | China | Asia | 2009 | W.1 | 2a |
| KF482477 | S10 | Canis familiaris | China | Asia | 2009 | W.1 | 2a |
| KF539789 | H-Erd | Canis familiaris | Hungary | Europe | 2012 | W.1 | 2a |
| KF539790 | H-Illatos | Canis familiaris | Hungary | Europe | 2012 | W.1 | 2a |
| KF638400 | s5 | Canis familiaris | China | Asia | 2010 | W.1 | 2a |
| KF676668 | CPV-JS2 | Canis familiaris | China | Asia | 2009 | W.1 | 2a |
| KF803599 | 2010-BJ-A64 | Canis familiaris | Thailand | Asia | 2010 | W.1 | 2a |
| KF803615 | 2011-BJ-B25 | Canis familiaris | Thailand | Asia | 2011 | W.1 | 2a |
| KF803618 | 2011-BJ-B31 | Canis familiaris | China | Asia | 2011 | W.1 | 2a |
| KF803636 | 2012-BJ-E34 | Canis familiaris | China | Asia | 2012 | W.1 | 2a |
| KF803640 | 2013-BJ-P21 | Canis familiaris | China | Asia | 2013 | W.1 | 2a |
| KF803643 | 2013-BJ-P34 | Canis familiaris | China | Asia | 2013 | W.1 | 2a |
| KJ186142 | JLDAAN08-2 | Canis familiaris | China | Asia | 2008 | W.1 | 2a |
| KJ438798 | Henan11 | Canis familiaris | China | Asia | 2011 | W.1 | 2a |
| KJ438799 | Henan12 | Canis familiaris | China | Asia | 2011 | W.1 | 2a |
| KJ438804 | Henan38 | Canis familiaris | China | Asia | 2012 | W.1 | 2a |
| KJ438805 | Henan42 | Canis familiaris | China | Asia | 2013 | W.1 | 2a |
| KJ674808 | 1-11 | Canis familiaris | China | Asia | 2013 | W.1 | 2a |
| KJ674813 | 1-5 | Canis familiaris | China | Asia | 2013 | W.1 | 2a |
| KJ674814 | 1-6 | Canis familiaris | China | Asia | 2013 | W.1 | 2a |
| KJ674819 | si | Canis familiaris | China | Asia | 2013 | W.1 | 2a |
| KJ813871 | CPV/Raccoon/VT/460/2013 | Procyon lotor | USA | North America | 2013 | W.1 | 2a |
| KM386821 | HLJ/01 | Canis familiaris | China | Asia | 2014 | W.1 | 2a |
| KM457102 | UY243.10 | Canis familiaris | Uruguay | South America | 2010 | W.1 | 2a |
| KM457132 | UY245.10 | Canis familiaris | Uruguay | South America | 2010 | W.1 | 2a |
| KM457133 | UY250.10 | Canis familiaris | Uruguay | South America | 2010 | W.1 | 2a |

| | | | | | | | |
|---|---|---|---|---|---|---|---|
| KM457134 | UY280.10 | Canis familiaris | Uruguay | South America | 2010 | W.1 | 2a |
| KM457135 | UY306.11 | Canis familiaris | Uruguay | South America | 2011 | W.1 | 2a |
| KM457136 | UY315.11 | Canis familiaris | Uruguay | South America | 2011 | W.1 | 2a |
| KM457137 | UY344.11 | Canis familiaris | Uruguay | South America | 2011 | W.1 | 2a |
| KM457138 | UY363.11 | Canis familiaris | Uruguay | South America | 2011 | W.1 | 2a |
| KM457139 | UY364rec | Canis familiaris | Uruguay | South America | 2011 | W.1 | 2a |
| KM457140 | UY365.11 | Canis familiaris | Uruguay | South America | 2011 | W.1 | 2a |
| KM457141 | UY370a | Canis familiaris | Uruguay | South America | 2011 | W.1 | 2a |
| KM457143 | UY364.11 | Canis familiaris | Uruguay | South America | 2011 | W.1 | 2a |
| KP686093 | CPV-ZXY | Canis familiaris | China | Asia | 2014 | W.1 | 2a |
| KP715659 | CPV-VT7 | Canis familiaris | Thailand | Asia | 2015 | W.1 | 2a |
| KP715663 | CPV-VT37 | Canis familiaris | Thailand | Asia | 2015 | W.1 | 2a |
| KP715671 | CPV-VT62 | Canis familiaris | Thailand | Asia | 2015 | W.1 | 2a |
| KP715673 | CPV-VT81 | Canis familiaris | Thailand | Asia | 2015 | W.1 | 2a |
| KP715676 | CPV-VT87 | Canis familiaris | Thailand | Asia | 2015 | W.1 | 2a |
| KP715682 | CPV-VT103 | Canis familiaris | Thailand | Asia | 2015 | W.1 | 2a |
| KP749837 | ANTU-1 | Canis familiaris | China | Asia | 2014 | W.1 | 2a |
| KR002792 | CPV/CN/SH1/2013 | Canis familiaris | China | Asia | 2013 | W.1 | 2a |
| KR002793 | CPV/CN/HB1/2013 | Canis familiaris | China | Asia | 2013 | W.1 | 2b |
| KR002794 | CPV/CN/HB3/2013 | Canis familiaris | China | Asia | 2013 | W.1 | 2a |
| KR002795 | CPV/CN/JL1/2013 | Canis familiaris | China | Asia | 2013 | W.1 | 2a |
| KR002796 | CPV/CN/JL3/2013 | Canis familiaris | China | Asia | 2013 | W.1 | 2b |
| KR002797 | CPV/CN/JL4/2013 | Canis familiaris | China | Asia | 2013 | W.1 | 2a |
| KR002798 | CPV/CN/JL5/2013 | Canis familiaris | China | Asia | 2013 | W.1 | 2a |
| KR002799 | CPV/CN/JL6/2013 | Canis familiaris | China | Asia | 2013 | W.1 | 2b |
| KR002800 | CPV/CN/LN1/2014 | Canis familiaris | China | Asia | 2014 | W.1 | 2a |
| KR002801 | CPV/CN/SD6/2014 | Canis familiaris | China | Asia | 2014 | W.1 | 2a |
| KR002802 | CPV/CN/SD9/2014 | Canis familiaris | China | Asia | 2014 | W.1 | 2a |
| KR002803 | CPV/CN/SD10/2014 | Canis familiaris | China | Asia | 2014 | W.1 | 2a |
| KR002804 | CPV/CN/SD18/2014 | Canis familiaris | China | Asia | 2014 | W.1 | 2a |
| KR002805 | CPV/CN/SD19/2014 | Canis familiaris | China | Asia | 2014 | W.1 | 2a |
| KR611467 | CPV-HB-14-9 | Canis familiaris | Thailand | Asia | 2014 | W.1 | 2b |
| KR611481 | CPV-HLJ-14-12 | Canis familiaris | China | Asia | 2014 | W.1 | 2a |

| | | | | | | | |
|---|---|---|---|---|---|---|---|
| KR611504 | CPV-LN-14-7 | Canis familiaris | China | Asia | 2014 | W.1 | 2b |
| KR869653 | CPV/BJ09 | Canis familiaris | China | Asia | 2014 | W.1 | 2a |
| KT156828 | MDJ-15 | Canis familiaris | China | Asia | 2014 | W.1 | 2a |
| KT156835 | HRB-ee7 | Canis familiaris | China | Asia | 2014 | W.1 | 2a |
| KT382542 | CPV-SH14 | Canis familiaris | China | Asia | 2014 | W.1 | 2a |
| KU983477 | YAD02 | Canis familiaris | China | Asia | 2016 | W.1 | 2a |
| KU983479 | YAD06 | Canis familiaris | China | Asia | 2016 | W.1 | 2a |
| KU983482 | YAD09 | Canis familiaris | China | Asia | 2016 | W.1 | 2a |
| KU983484 | YAD12 | Canis familiaris | China | Asia | 2016 | W.1 | 2a |
| KU983485 | YAD13 | Canis familiaris | China | Asia | 2016 | W.1 | 2a |
| KU983486 | YAD14 | Canis familiaris | China | Asia | 2016 | W.1 | 2a |
| KU983487 | YAD15 | Canis familiaris | China | Asia | 2016 | W.1 | 2a |
| KU983488 | YAD18 | Canis familiaris | China | Asia | 2016 | W.1 | 2a |
| KU983490 | YAD22 | Canis familiaris | China | Asia | 2016 | W.1 | 2a |
| KU983491 | YAD23 | Canis familiaris | China | Asia | 2016 | W.1 | 2a |
| KX219736 | TKM-Orissa | Canis familiaris | India | Asia | 2012 | W.1 | 2a |
| KX219737 | Ker-2 | Canis familiaris | India | Asia | 2015 | W.1 | 2a |
| KX219738 | Ker-3 | Canis familiaris | India | Asia | 2015 | W.1 | 2a |
| KX434454 | CPV_IZSSI_29451_09 | Canis familiaris | Italy | Europe | 2009 | W.1 | 2a |
| KX469431 | Alok/new CPV-2a/2012/Ind | Canis familiaris | India | Asia | 2012 | W.1 | 2a |
| KX469433 | KN-5/new CPV-2a/2015/Ind | Canis familiaris | India | Asia | 2015 | W.1 | 2a |
| KX469435 | RJ-7/new CPV-2a/2016/Ind | Canis familiaris | India | Asia | 2016 | W.1 | 2a |
| KX618915 | 2a | Paradoxurus musangus | Singapore | Asia | 2016 | W.1 | 2a |
| KY083096 | M2-2 | NI | Singapore | Asia | 2014 | W.1 | 2a |
| KY083097 | M272-6 | NI | Singapore | Asia | 2014 | W.1 | 2a |
| KY386850 | GY-1 | Canis familiaris | China | Asia | 2015 | W.1 | 2a |
| KY386852 | GY-3 | Canis familiaris | China | Asia | 2015 | W.1 | 2a |
| KY386855 | GY-6 | Canis familiaris | China | Asia | 2016 | W.1 | 2a |
| KY403998 | CPV-YH | Canis familiaris | China | Asia | 2008 | W.1 | 2a |
| LC214970 | CPV/dog/HCM/22/2013 | Canis familiaris | Vietnam | Asia | 2012 | W.1 | 2a |
| LC270891 | 9985 | Canis familiaris | Japan | Asia | 2017 | W.1 | 2b |
| LC270892 | 9985-46 | Canis familiaris | Japan | Asia | 2017 | W.1 | 2b |
| MF069442 | CPV/Raccoon/RC14/BC_2010 | Procyon Lotor | Canada | North America | 2010 | W.1 | 2a |

| | | | | | | | |
|---|---|---|---|---|---|---|---|
| MF069443 | CPV/Raccoon/RC19/BC_2016 | Procyon Lotor | Canada | North America | 2016 | W.1 | 2a |
| MF069444 | CPV/Raccoon/RC20/BC_2016 | Procyon Lotor | Canada | North America | 2016 | W.1 | 2a |
| MF134808 | CPV-BJ03/17 | Canis familiaris | China | Asia | 2017 | W.1 | 2a |
| MF423125 | CPV/Coyote/C67/NL_2014 | Canis Latrans | Canada | North America | 2014 | W.1 | 2a |
| MF467224 | CPV-ZJ1579 | Canis familiaris | China | Asia | 2015 | W.1 | 2a |
| MF467226 | CPV-JS1591 | Canis familiaris | China | Asia | 2015 | W.1 | 2a |
| MF467228 | CPV-HN1618 | Canis familiaris | China | Asia | 2016 | W.1 | 2a |
| MF467230 | CPV-HN1616 | Canis familiaris | China | Asia | 2016 | W.1 | 2a |
| MF467233 | CPV-HN1587 | Canis familiaris | China | Asia | 2015 | W.1 | 2b |
| MF467234 | CPV-HN1586 | Canis familiaris | China | Asia | 2015 | W.1 | 2a |
| MF467235 | CPV-HN1582 | Canis familiaris | China | Asia | 2015 | W.1 | 2a |
| MF467237 | CPV-HN1524 | Canis familiaris | China | Asia | 2015 | W.1 | 2a |
| MF467240 | CPV-HN1506 | Canis familiaris | China | Asia | 2015 | W.1 | 2a |
| MF467241 | CPV-HN1503 | Canis familiaris | China | Asia | 2015 | W.1 | 2a |
| MF805790 | Canine/China/02/2016 | Canis familiaris | China | Asia | 2016 | W.1 | 2a |
| MF805791 | Canine/China/03/2016 | Canis familiaris | China | Asia | 2016 | W.1 | 2a |
| MF805793 | Canine/China/05/2016 | Canis familiaris | China | Asia | 2016 | W.1 | 2a |
| MF805794 | Canine/China/06/2016 | Canis familiaris | China | Asia | 2016 | W.1 | 2a |
| MF805798 | Canine/China/10/2016 | Canis familiaris | China | Asia | 2016 | W.1 | 2a |
| MG434738 | CPV_IZSSI_PA43847/2016 | Canis familiaris | Italy | Europe | 2017 | W.1 | 2a |
| MG434739 | CPV_IZSSI_PA48686/2016 | Canis familiaris | Italy | Europe | 2016 | W.1 | 2a |
| MG434740 | CPV_IZSSI_PA3213/2017 | Canis familiaris | Italy | Europe | 2017 | W.1 | 2a |
| MG434741 | CPV_IZSSI_PA5610/2017 | Canis familiaris | Italy | Europe | 2017 | W.1 | 2a |
| MG434742 | CPV_IZSSI_PA10388/2017 | Canis familiaris | Italy | Europe | 2017 | W.1 | 2a |
| MG434743 | CPV_IZSSI_PA13577/2017 | Canis familiaris | Italy | Europe | 2017 | W.1 | 2a |
| MG434744 | CPV_IZSSI_PA13579id90/2017 | Canis familiaris | Italy | Europe | 2017 | W.1 | 2a |
| MG434745 | CPV_IZSSI_PA13579id93/2017 | Canis familiaris | Italy | Europe | 2017 | W.1 | 2a |
| MG583676 | CPV/CN/ya1/2017 | Yak | China | Asia | 2015 | W.1 | 2a |
| MH106698 | CPV-BJL1 | Canis familiaris | China | Asia | 2015 | W.1 | 2a |
| MH106699 | CPV-BJL2 | Canis familiaris | China | Asia | 2015 | W.1 | 2b |
| MH106700 | CPV-BJL3 | Canis familiaris | China | Asia | 2016 | W.1 | 2c |
| MH476580 | Canine/China/11/2017 | Canis familiaris | China | Asia | 2017 | W.1 | 2a |
| MH476582 | Canine/China/13/2017 | Canis familiaris | China | Asia | 2017 | W.1 | 2a |

| | | | | | | | |
|---|---|---|---|---|---|---|---|
| MH476586 | Canine/China/17/2017 | Canis familiaris | China | Asia | 2017 | W.1 | 2a |
| MH476588 | Canine/China/19/2017 | Canis familiaris | China | Asia | 2017 | W.1 | 2b |
| MH476589 | Canine/China/20/2017 | Canis familiaris | China | Asia | 2017 | W.1 | 2a |
| MH476590 | Canine/China/21/2017 | Canis familiaris | China | Asia | 2017 | W.1 | 2a |
| MH476591 | Canine/China/22/2017 | Canis familiaris | China | Asia | 2017 | W.1 | 2a |
| MH476593 | Canine/China/24/2017 | Canis familiaris | China | Asia | 2017 | W.1 | 2a |
| MH545963 | TN/CPV2a/2018 | Canis familiaris | India | Asia | 2018 | W.1 | 2a |
| MK344435 | SV27/16 | Canis familiaris | Brazil | South America | 2016 | W.1 | 2a |
| MK344437 | SV56/18 | Canis familiaris | Brazil | South America | 2018 | W.1 | 2a |
| MK344442 | SV164/17 | Canis familiaris | Brazil | South America | 2017 | W.1 | 2a |
| MK344443 | SV166/17 | Canis familiaris | Brazil | South America | 2017 | W.1 | 2a |
| MK344444 | SV168/17 | Canis familiaris | Brazil | South America | 2017 | W.1 | 2a |
| MK344445 | SV176/17 | Canis familiaris | Brazil | South America | 2017 | W.1 | 2a |
| MK344447 | SV184/17 | Canis familiaris | Brazil | South America | 2017 | W.1 | 2a |
| MK344449 | SV187/17 | Canis familiaris | Brazil | South America | 2017 | W.1 | 2a |
| MK344451 | SV210/16 | Canis familiaris | Brazil | South America | 2016 | W.1 | 2a |
| MK344468 | SV697/15 | Canis familiaris | Brazil | South America | 2015 | W.1 | 2a |
| MK413740 | CPV-2a_PA30636/17 | Canis familiaris | Italy | Europe | 2017 | W.1 | 2a |
| MK413741 | CPV-2a_PA31209/17 | Canis familiaris | Italy | Europe | 2017 | W.1 | 2a |
| MK895483 | IZSSI_PA1464/19_idUV1 | Canis familiaris | Nigeria | Africa | 2018 | W.1 | 2a |
| MK895484 | IZSSI_PA1464/19_idYV8 | Canis familiaris | Nigeria | Africa | 2018 | W.1 | 2a |
| MK895485 | IZSSI_PA1464/19_idUV6 | Canis familiaris | Nigeria | Africa | 2018 | W.1 | 2a |
| MN451673 | CPV307 | Canis familiaris | USA | North America | 1993 | W.1 | 2a |
| MN451689 | CPV616 | Canis familiaris | Nigeria | Africa | 2018 | W.1 | 2a |
| MN661243 | CPV new 2a variant | Canis familiaris | India | Asia | 2016 | W.1 | 2a |
| MN862741 | CPV-2/American mink/MIVI-73/BC_2019 | Neovison vison | Canada | North America | 2019 | W.1 | 2a |
| MN862742 | CPV-2/River otter/OTVI-13/BC_2019 | Lontra Canadensis | Canada | North America | 2019 | W.1 | 2a |
| MW648348 | 1-UCS | Canis familiaris | Brazil | South America | 2019 | W.1 | 2a |
| MW648349 | 2-UCS | Canis familiaris | Brazil | South America | 2019 | W.1 | 2a |
| MW648350 | 3-UCS | Canis familiaris | Brazil | South America | 2020 | W.1 | 2a |
| MW648351 | 4-UCS | Canis familiaris | Brazil | South America | 2020 | W.1 | 2a |
| MW648352 | 5-UCS | Canis familiaris | Brazil | South America | 2020 | W.1 | 2a |
| MW648353 | 6-UCS | Canis familiaris | Brazil | South America | 2020 | W.1 | 2a |

| | | | | | | | |
|---|---|---|---|---|---|---|---|
| MW648354 | 7-UCS | Canis familiaris | Brazil | South America | 2020 | W.1 | 2a |
| MW648355 | 8-UCS | Canis familiaris | Brazil | South America | 2020 | W.1 | 2a |
| MW648356 | 11-UCS | Canis familiaris | Brazil | South America | 2020 | W.1 | 2a |
| MW648357 | 14-UCS | Canis familiaris | Brazil | South America | 2020 | W.1 | 2a |
| MW648358 | 15-UCS | Canis familiaris | Brazil | South America | 2020 | W.1 | 2a |
| MW648359 | 18-UCS | Canis familiaris | Brazil | South America | 2020 | W.1 | 2a |
| MW648360 | 19-UCS | Canis familiaris | Brazil | South America | 2020 | W.1 | 2a |
| MW648361 | 21-UCS | Canis familiaris | Brazil | South America | 2020 | W.1 | 2a |
| MW648362 | 23-UCS | Canis familiaris | Brazil | South America | 2020 | W.1 | 2a |
| MW648363 | 25-UCS | Canis familiaris | Brazil | South America | 2020 | W.1 | 2a |
| MW648364 | 26-UCS | Canis familiaris | Brazil | South America | 2020 | W.1 | 2a |
| MW648365 | 27-UCS | Canis familiaris | Brazil | South America | 2020 | W.1 | 2c |
| MW648366 | 28-UCS | Canis familiaris | Brazil | South America | 2020 | W.1 | 2c |
| MW648367 | 29-UCS | Canis familiaris | Brazil | South America | 2020 | W.1 | 2a |
| U72695 | T4 | Canis familiaris | Taiwan | Asia | 1996 | W.1 | 2a |
| U72698 | T37 | Canis familiaris | Taiwan | Asia | 1996 | W.1 | 2a |
| AY742932 | CPV2b-193.us.91 | Canis familiaris | USA | North America | 1991 | W.2 | 2b |
| AY742934 | CPV-447 | Canis familiaris | Germany | Europe | 1995 | W.2 | 2b |
| AY742936 | CPV2b-395.us.98 | Canis familiaris | USA | North America | 1998 | W.2 | 2b |
| AY742955 | CPV-436 | Canis familiaris | USA | North America | 2003 | W.2 | 2b |
| DQ025961 | 03C5 | Canis familiaris | France | Europe | 2005 | W.2 | 2b |
| DQ025991 | 04S22 | Canis familiaris | France | Europe | 2005 | W.2 | 2b |
| EU009205 | K029 | Canis familiaris | South Korea | Asia | 2007 | W.2 | 2b |
| EU659119 | CPV2b-410.us.00 | Canis familiaris | USA | North America | 2000 | W.2 | 2b |
| EU659120 | CPV2b-411a.us.98 | Canis familiaris | USA | North America | 1998 | W.2 | 2b |
| EU659121 | CPV2b-411b.us.98 | Canis familiaris | USA | North America | 1998 | W.2 | 2b |
| FJ005195 | 136/00 | Canis familiaris | Italy | Europe | 2000 | W.2 | 2c |
| FJ005196 | G7/97 | Canis familiaris | Germany | Europe | 1997 | W.2 | 2c |
| FJ005203 | G52-9/2-98 | Canis familiaris | Germany | Europe | 1998 | W.2 | 2c |
| FJ005204 | G333/99 | Canis familiaris | Germany | Europe | 1999 | W.2 | 2c |
| FJ005205 | 279/04 | Canis familiaris | Italy | Europe | 2004 | W.2 | 2c |
| FJ005206 | 287/04 | Canis familiaris | Italy | Europe | 2004 | W.2 | 2c |
| FJ005207 | 290/04 | Canis familiaris | Italy | Europe | 2004 | W.2 | 2c |

| | | | | | | | |
|---|---|---|---|---|---|---|---|
| FJ005208 | 291/04 | Canis familiaris | Italy | Europe | 2004 | W.2 | 2c |
| FJ005209 | 303/04 | Canis familiaris | Italy | Europe | 2004 | W.2 | 2c |
| FJ005210 | 307/04 | Canis familiaris | Italy | Europe | 2004 | W.2 | 2c |
| FJ005211 | 342/04 | Canis familiaris | Italy | Europe | 2004 | W.2 | 2c |
| FJ005212 | 349/04 | Canis familiaris | Italy | Europe | 2004 | W.2 | 2c |
| FJ005213 | 9/05 | Canis familiaris | Italy | Europe | 2005 | W.2 | 2c |
| FJ005214 | 67/06 | Canis familiaris | Spain | Europe | 2006 | W.2 | 2c |
| FJ005215 | 252/06 | Canis familiaris | Italy | Europe | 2006 | W.2 | 2c |
| FJ005216 | 284/06 | Canis familiaris | Italy | Europe | 2006 | W.2 | 2c |
| FJ005217 | 327/06 | Canis familiaris | Italy | Europe | 2006 | W.2 | 2c |
| FJ005218 | 330/06 | Canis familiaris | Italy | Europe | 2006 | W.2 | 2c |
| FJ005219 | 336/06 | Canis familiaris | Italy | Europe | 2006 | W.2 | 2c |
| FJ005220 | 337/06 | Canis familiaris | Italy | Europe | 2006 | W.2 | 2c |
| FJ005221 | 340/06 | Canis familiaris | Italy | Europe | 2006 | W.2 | 2c |
| FJ005222 | 359/06 | Canis familiaris | Italy | Europe | 2006 | W.2 | 2c |
| FJ005223 | 365/06 | Canis familiaris | Italy | Europe | 2006 | W.2 | 2c |
| FJ005224 | 367/06 | Canis familiaris | Italy | Europe | 2006 | W.2 | 2c |
| FJ005225 | 382/06 | Canis familiaris | Italy | Europe | 2006 | W.2 | 2c |
| FJ005226 | 383/06 | Canis familiaris | Italy | Europe | 2006 | W.2 | 2c |
| FJ005227 | 389/06 | Canis familiaris | Italy | Europe | 2006 | W.2 | 2c |
| FJ005228 | 393/06 | Canis familiaris | Italy | Europe | 2006 | W.2 | 2c |
| FJ005229 | 397/06 | Canis familiaris | Italy | Europe | 2006 | W.2 | 2c |
| FJ005230 | 398/06 | Canis familiaris | Italy | Europe | 2006 | W.2 | 2c |
| FJ005231 | 406/06 | Canis familiaris | Italy | Europe | 2006 | W.2 | 2c |
| FJ005232 | 411/06 | Canis familiaris | Italy | Europe | 2006 | W.2 | 2c |
| FJ005233 | 40/07 | Canis familiaris | Italy | Europe | 2007 | W.2 | 2c |
| FJ005234 | 43/03 | Canis familiaris | Italy | Europe | 2007 | W.2 | 2c |
| FJ005235 | 67/07-11 | Canis familiaris | USA | North America | 2007 | W.2 | 2c |
| FJ005236 | 110/07-27 | Canis familiaris | USA | North America | 2007 | W.2 | 2c |
| FJ005237 | 159/07 | Canis familiaris | Italy | Europe | 2007 | W.2 | 2c |
| FJ005238 | 158/07 | Canis familiaris | Italy | Europe | 2007 | W.2 | 2c |
| FJ005239 | 165/07-A | Canis familiaris | Italy | Europe | 2007 | W.2 | 2c |
| FJ005240 | 208/07 | Canis familiaris | Italy | Europe | 2007 | W.2 | 2c |

| | | | | | | | |
|---|---|---|---|---|---|---|---|
| FJ005241 | 215/07-2 | Canis familiaris | Italy | Europe | 2007 | W.2 | 2c |
| FJ005242 | 217/07 | Canis familiaris | Italy | Europe | 2007 | W.2 | 2c |
| FJ005243 | 243/07 | Canis familiaris | Italy | Europe | 2007 | W.2 | 2c |
| FJ005244 | 127/08-A | Canis familiaris | Italy | Europe | 2008 | W.2 | 2c |
| FJ005245 | 127/08-B | Canis familiaris | Italy | Europe | 2008 | W.2 | 2c |
| FJ005246 | 128/08 | Canis familiaris | Spain | Europe | 2008 | W.2 | 2c |
| FJ005247 | 195/08 | Canis familiaris | Belgium | Europe | 2008 | W.2 | 2c |
| FJ005248 | 219/08-2 | Canis familiaris | Italy | Europe | 2008 | W.2 | 2c |
| FJ005249 | 219/08-5 | Canis familiaris | Italy | Europe | 2008 | W.2 | 2c |
| FJ005250 | 219/08-13 | Canis familiaris | Italy | Europe | 2008 | W.2 | 2c |
| FJ005251 | 239/08 | Canis familiaris | Italy | Europe | 2008 | W.2 | 2c |
| FJ005257 | 54/08 | Canis familiaris | Italy | Europe | 2008 | W.2 | 2a |
| FJ005260 | G82/97 | Canis familiaris | Germany | Europe | 1997 | W.2 | 2b |
| FJ005262 | 311/04 | Canis familiaris | Italy | Europe | 2004 | W.2 | 2b |
| FJ005264 | 134/05 | Canis familiaris | Italy | Europe | 2005 | W.2 | 2b |
| FJ222821 | 56/00 | Canis familiaris | Italy | Europe | 2000 | W.2 | 2c |
| GQ865518 | GR51/08 | Canis familiaris | Greece | Europe | 2008 | W.2 | 2c |
| GQ865519 | GR09/09 | Canis familiaris | Greece | Europe | 2009 | W.2 | 2c |
| GU362935 | cat234/08 | Felis catus | Italy | Europe | 2008 | W.2 | 2c |
| GU569944 | GZ0201 | Canis familiaris | China | Asia | 2002 | W.2 | 2b |
| JF414818 | Arg32 | Canis familiaris | Argentina | South America | 2008 | W.2 | 2c |
| JF414819 | Arg35 | Canis familiaris | Argentina | South America | 2008 | W.2 | 2c |
| JF414820 | Arg44 | Canis familiaris | Argentina | South America | 2009 | W.2 | 2c |
| JF414821 | Arg48 | Canis familiaris | Argentina | South America | 2009 | W.2 | 2c |
| JF414822 | Arg64 | Canis familiaris | Argentina | South America | 2010 | W.2 | 2c |
| JF414823 | Arg60 | Canis familiaris | Argentina | South America | 2009 | W.2 | 2c |
| JF414824 | Arg66 | Canis familiaris | Argentina | South America | 2010 | W.2 | 2c |
| JF414825 | Arg67 | Canis familiaris | Argentina | South America | 2010 | W.2 | 2c |
| JF414826 | Arg 68 | Canis familiaris | Argentina | South America | 2010 | W.2 | 2c |
| JN033694 | Laika-1993 | Canis familiaris | Russia | Europe | 1993 | W.2 | 2b |
| JN867602 | CPV-2b/Dog/CA/148743/08 | Canis familiaris | USA | North America | 2008 | W.2 | 2b |
| JN867603 | CPV-2b/Dog/KS/81213/09 | Canis familiaris | USA | North America | 2009 | W.2 | 2b |
| JN867604 | CPV-2b/Dog/IL/137654/08 | Canis familiaris | USA | North America | 2008 | W.2 | 2b |

| | | | | | | | |
|---|---|---|---|---|---|---|---|
| JX305946 | 46/09-4060 | Canis familiaris | Italy | Europe | 2009 | W.2 | 2c |
| JX305947 | 46/09-4061 | Canis familiaris | Italy | Europe | 2009 | W.2 | 2c |
| JX305948 | 140/09-15 | Canis familiaris | Italy | Europe | 2009 | W.2 | 2c |
| JX305949 | 140/09-17 | Canis familiaris | Italy | Europe | 2009 | W.2 | 2c |
| JX305950 | 140/09-33 | Canis familiaris | Italy | Europe | 2009 | W.2 | 2c |
| JX305951 | 341/09 | Canis familiaris | Italy | Europe | 2009 | W.2 | 2c |
| JX305952 | 10/10 | Canis familiaris | Italy | Europe | 2009 | W.2 | 2c |
| JX305953 | 289/10 | Canis familiaris | Italy | Europe | 2010 | W.2 | 2c |
| JX305954 | 261/10 | Canis familiaris | Italy | Europe | 2010 | W.2 | 2c |
| JX305955 | 275/10-A | Canis familiaris | Italy | Europe | 2010 | W.2 | 2c |
| JX305956 | 275/10-B | Canis familiaris | Italy | Europe | 2010 | W.2 | 2c |
| JX305957 | 480/10 | Canis familiaris | Italy | Europe | 2010 | W.2 | 2c |
| JX305958 | 132/11-4 | Canis familiaris | Italy | Europe | 2011 | W.2 | 2c |
| JX305959 | 132/11-33 | Canis familiaris | Italy | Europe | 2011 | W.2 | 2c |
| JX305960 | 503/11-1 | Canis familiaris | Italy | Europe | 2011 | W.2 | 2c |
| JX305961 | 503/11-2 | Canis familiaris | Italy | Europe | 2011 | W.2 | 2c |
| JX305962 | 503/11-3 | Canis familiaris | Italy | Europe | 2011 | W.2 | 2c |
| JX305963 | 503/11-4 | Canis familiaris | Italy | Europe | 2011 | W.2 | 2c |
| JX305964 | 503/11-5 | Canis familiaris | Italy | Europe | 2011 | W.2 | 2c |
| JX305965 | 271/12-Miky | Canis familiaris | Italy | Europe | 2012 | W.2 | 2c |
| JX475237 | CT/372/11 | Procyon lotor | USA | Asia | 2011 | W.2 | 2b |
| JX475242 | WI/18268/02 | Canis lupus nubilus | USA | Asia | 2002 | W.2 | 2b |
| JX475247 | CO/1246/10 | Puma concolor | USA | North America | 2010 | W.2 | 2b |
| JX475251 | CO/2235/09 | Puma concolor | USA | North America | 2009 | W.2 | 2b |
| JX475278 | AR/1069/12 | Canis latrans | USA | North America | 2012 | W.2 | 2b |
| KJ813827 | CPV/Fisher/ND/F1M111211/2013 | Martes pennanti | USA | North America | 2013 | W.2 | 2b |
| KJ813828 | CPV/Fisher/ND/F1F010712/2013 | Martes pennanti | USA | North America | 2013 | W.2 | 2b |
| KJ813844 | CPV/Bobcat/ND/885/2013 | Lynx rufus | USA | North America | 2013 | W.2 | 2b |
| KJ813851 | CPV/Bobcat/ND/1168/2013 | Lynx rufus | USA | North America | 2013 | W.2 | 2b |
| KJ813852 | CPV/Bobcat/ND/1170/2013 | Lynx rufus | USA | North America | 2013 | W.2 | 2b |
| KJ813873 | CPV/Gray wolf/MI/850/2012 | Canis lupus | USA | North America | 2012 | W.2 | 2b |
| KJ813881 | CPV/Gray wolf/MI/832/2012 | Canis lupus | USA | North America | 2012 | W.2 | 2b |
| KJ813882 | CPV/Raccoon/NJ/1423/2012 | Procyon lotor | USA | North America | 2012 | W.2 | 2b |

| | | | | | | | |
|---|---|---|---|---|---|---|---|
| KJ813892 | CPV/Coyote/AK/218/2013 | Canis latrans | USA | North America | 2013 | W.2 | 2b |
| KM457103 | UY12.06 | Canis familiaris | Uruguay | South America | 2006 | W.2 | 2c |
| KM457104 | UY47.06 | Canis familiaris | Uruguay | South America | 2006 | W.2 | 2c |
| KM457105 | UY52.06 | Canis familiaris | Uruguay | South America | 2006 | W.2 | 2c |
| KM457106 | UY55.06 | Canis familiaris | Uruguay | South America | 2006 | W.2 | 2c |
| KM457107 | UY72.07 | Canis familiaris | Uruguay | South America | 2007 | W.2 | 2c |
| KM457108 | UY82.07 | Canis familiaris | Uruguay | South America | 2007 | W.2 | 2c |
| KM457109 | UY95.07 | Canis familiaris | Uruguay | South America | 2007 | W.2 | 2c |
| KM457110 | UY101.07 | Canis familiaris | Uruguay | South America | 2007 | W.2 | 2c |
| KM457111 | UY120.08 | Canis familiaris | Uruguay | South America | 2008 | W.2 | 2c |
| KM457112 | UY135.08 | Canis familiaris | Uruguay | South America | 2008 | W.2 | 2c |
| KM457113 | UY152.08 | Canis familiaris | Uruguay | South America | 2008 | W.2 | 2c |
| KM457114 | UY169.08 | Canis familiaris | Uruguay | South America | 2008 | W.2 | 2c |
| KM457115 | UY173.09 | Canis familiaris | Uruguay | South America | 2009 | W.2 | 2c |
| KM457116 | UY185.09 | Canis familiaris | Uruguay | South America | 2009 | W.2 | 2c |
| KM457117 | UY187.09 | Canis familiaris | Uruguay | South America | 2009 | W.2 | 2c |
| KM457118 | UY190.09 | Canis familiaris | Uruguay | South America | 2009 | W.2 | 2c |
| KM457119 | UY235.10 | Canis familiaris | Uruguay | South America | 2010 | W.2 | 2c |
| KM457120 | UY242.10 | Canis familiaris | Uruguay | South America | 2010 | W.2 | 2c |
| KM457121 | UY247.10 | Canis familiaris | Uruguay | South America | 2010 | W.2 | 2c |
| KM457122 | UY258.10 | Canis familiaris | Uruguay | South America | 2010 | W.2 | 2c |
| KM457123 | UY261.10 | Canis familiaris | Uruguay | South America | 2010 | W.2 | 2c |
| KM457124 | UY307.11 | Canis familiaris | Uruguay | South America | 2011 | W.2 | 2c |
| KM457125 | UY317.11 | Canis familiaris | Uruguay | South America | 2011 | W.2 | 2c |
| KM457126 | UY318.10 | Canis familiaris | Uruguay | South America | 2010 | W.2 | 2c |
| KM457127 | UY326.11 | Canis familiaris | Uruguay | South America | 2011 | W.2 | 2c |
| KM457128 | UY346.11 | Canis familiaris | Uruguay | South America | 2011 | W.2 | 2c |
| KM457129 | UY349.11 | Canis familiaris | Uruguay | South America | 2011 | W.2 | 2c |
| KM457130 | UY354.11 | Canis familiaris | Uruguay | South America | 2011 | W.2 | 2c |
| KM457131 | UY368.11 | Canis familiaris | Uruguay | South America | 2011 | W.2 | 2c |
| KM457142 | UY370c | Canis familiaris | Uruguay | South America | 2011 | W.2 | 2c |
| KU508407 | CPV_IZSSI_25835_09 | Canis familiaris | Italy | Europe | 2009 | W.2 | 2c |
| KU508691 | HB | Canis familiaris | Australia | Oceania | 2015 | W.2 | 2c |

| | | | | | | | |
|---|---|---|---|---|---|---|---|
| KU508692 | FH | Canis familiaris | Australia | Oceania | 2015 | W.2 | 2c |
| KU508693 | LW | Canis familiaris | Australia | Oceania | 2015 | W.2 | 2c |
| KX434455 | CPV_IZSSI_23782_09 | Canis familiaris | Italy | Europe | 2009 | W.2 | 2c |
| KX434456 | CPV_IZSSI_45361_09 | Canis familiaris | Italy | Europe | 2009 | W.2 | 2c |
| KX434458 | CPV_IZSSI_2323_11 | Canis familiaris | Italy | Europe | 2011 | W.2 | 2c |
| KX434459 | CPV_IZSSI_27692_1_11 | Canis familiaris | Italy | Europe | 2011 | W.2 | 2c |
| KX434460 | CPV_IZSSI_52238_12 | Canis familiaris | Italy | Europe | 2012 | W.2 | 2c |
| KY073269 | UFMT | Canis familiaris | Brazil | South America | 2015 | W.2 | 2c |
| LC214969 | CPV/dog/HCM/7/2013 | Canis familiaris | Vietnam | Asia | 2013 | W.2 | 2c |
| MF177225 | 242-98 | Canis familiaris | Italy | Europe | 1998 | W.2 | 2b |
| MF177227 | 202-09 | Canis familiaris | France | Europe | 2009 | W.2 | 2c |
| MF177228 | 485-09 | Canis familiaris | Italy | Europe | 2009 | W.2 | 2c |
| MF177229 | 368-12-17 | Canis familiaris | Albana | Europe | 2012 | W.2 | 2c |
| MF177230 | 57-10 | Canis familiaris | Italy | Europe | 2010 | W.2 | 2c |
| MF177232 | 201-98 | Canis familiaris | Italy | Europe | 1998 | W.2 | 2b |
| MF177234 | 347-03 | Canis familiaris | Italy | Europe | 2003 | W.2 | 2c |
| MF177235 | 283-06 | Canis familiaris | Italy | Europe | 2006 | W.2 | 2c |
| MF177236 | 244-04 | Canis familiaris | Italy | Europe | 2004 | W.2 | 2c |
| MF177237 | 188-07 | Canis familiaris | Italy | Europe | 2007 | W.2 | 2c |
| MF177238 | 114-05 | Canis familiaris | Italy | Europe | 2005 | W.2 | 2c |
| MF177239 | 288-01 | Canis familiaris | Italy | Europe | 2001 | W.2 | 2c |
| MF177240 | 189-02 | Canis familiaris | Italy | Europe | 2002 | W.2 | 2c |
| MF177242 | Arg26 | Canis familiaris | Argentina | South America | 2008 | W.2 | 2c |
| MF177243 | Arg32 | Canis familiaris | Argentina | South America | 2008 | W.2 | 2c |
| MF177244 | Arg33 | Canis familiaris | Argentina | South America | 2008 | W.2 | 2c |
| MF177245 | Arg35 | Canis familiaris | Argentina | South America | 2008 | W.2 | 2c |
| MF177249 | Arg71 | Canis familiaris | Argentina | South America | 2010 | W.2 | 2c |
| MF177250 | BRA01/10 | Canis familiaris | Brazil | South America | 2010 | W.2 | 2c |
| MF177252 | BRA02/10 | Canis familiaris | Brazil | South America | 2010 | W.2 | 2c |
| MF177253 | BRA03/10 | Canis familiaris | Brazil | South America | 2010 | W.2 | 2c |
| MF177254 | BRA04/10 | Canis familiaris | Brazil | South America | 2010 | W.2 | 2c |
| MF177255 | BRA01/14 | Canis familiaris | Brazil | South America | 2014 | W.2 | 2c |
| MF177257 | BRA02/14 | Canis familiaris | Brazil | South America | 2014 | W.2 | 2c |

| | | | | | | | |
|---|---|---|---|---|---|---|---|
| MF177260 | BRA01/12 | Canis familiaris | Brazil | South America | 2012 | W.2 | 2c |
| MF177261 | BRA04/13 | Canis familiaris | Brazil | South America | 2013 | W.2 | 2c |
| MF177262 | Py | Canis familiaris | Paraguay | South America | 2009 | W.2 | 2c |
| MF177263 | E1.11 | Canis familiaris | Ecuador | South America | 2011 | W.2 | 2c |
| MF177264 | E10.11 | Canis familiaris | Ecuador | South America | 2011 | W.2 | 2c |
| MF177265 | E12.11 | Canis familiaris | Ecuador | South America | 2011 | W.2 | 2a |
| MF177266 | E13.11 | Canis familiaris | Ecuador | South America | 2011 | W.2 | 2c |
| MF177267 | E16.11 | Canis familiaris | Ecuador | South America | 2011 | W.2 | 2c |
| MF177268 | E19.11 | Canis familiaris | Ecuador | South America | 2011 | W.2 | 2a |
| MF177269 | E20.11 | Canis familiaris | Ecuador | South America | 2011 | W.2 | 2b |
| MF177270 | E23.11 | Canis familiaris | Ecuador | South America | 2011 | W.2 | 2c |
| MF177271 | E26.11 | Canis familiaris | Ecuador | South America | 2011 | W.2 | 2c |
| MF177272 | E28.11 | Canis familiaris | Ecuador | South America | 2011 | W.2 | 2c |
| MF177273 | E29.11 | Canis familiaris | Ecuador | South America | 2011 | W.2 | 2c |
| MF177274 | E31.11 | Canis familiaris | Ecuador | South America | 2011 | W.2 | 2c |
| MF177275 | E32.11 | Canis familiaris | Ecuador | South America | 2011 | W.2 | 2c |
| MF177276 | E35.11 | Canis familiaris | Ecuador | South America | 2011 | W.2 | 2a |
| MF177277 | E36.11 | Canis familiaris | Ecuador | South America | 2011 | W.2 | 2a |
| MF177278 | E4.11 | Canis familiaris | Ecuador | South America | 2011 | W.2 | 2c |
| MF177279 | E48.11 | Canis familiaris | Ecuador | South America | 2011 | W.2 | 2c |
| MF177280 | E6.11 | Canis familiaris | Ecuador | South America | 2011 | W.2 | 2b |
| MF177282 | UY21.06 | Canis familiaris | Uruguay | South America | 2006 | W.2 | 2c |
| MF177283 | UY181.09 | Canis familiaris | Uruguay | South America | 2009 | W.2 | 2c |
| MF177284 | UY196.09 | Canis familiaris | Uruguay | South America | 2009 | W.2 | 2c |
| MF177285 | UY269.10 | Canis familiaris | Uruguay | South America | 2010 | W.2 | 2c |
| MF177286 | UY375.11 | Canis familiaris | Uruguay | South America | 2011 | W.2 | 2c |
| MF423123 | CPV/Coyote/C16/NL_2014 | Canis Latrans | Canada | North America | 2014 | W.2 | 2b |
| MF423124 | CPV/Coyote/C55/NL_2014 | Canis Latrans | Canada | North America | 2014 | W.2 | 2b |
| MF457594 | OH20219 | Canis familiaris | USA | North America | 2015 | W.2 | 2c |
| MF510157 | CPV_IZSSI_2743_17 | Canis familiaris | Italy | Europe | 2017 | W.2 | 2c |
| MF510158 | CPV_IZSSI_41113_c1_16 | Canis familiaris | Italy | Europe | 2016 | W.2 | 2c |
| MF805789 | Canine/China/01/2016 | Canis familiaris | China | Asia | 2016 | W.2 | 2c |
| MF805792 | Canine/China/04/2016 | Canis familiaris | China | Asia | 2016 | W.2 | 2c |

| | | | | | | | |
|---|---|---|---|---|---|---|---|
| MF805795 | Canine/China/07/2016 | Canis familiaris | China | Asia | 2016 | W.2 | 2c |
| MF805796 | Canine/China/08/2016 | Canis familiaris | China | Asia | 2016 | W.2 | 2c |
| MF805797 | Canine/China/09/2016 | Canis familiaris | China | Asia | 2016 | W.2 | 2c |
| MG013488 | CPV-SH1516 | Canis familiaris | China | Asia | 2017 | W.2 | 2c |
| MG264076 | EC/04/2017 | Canis familiaris | Ecuador | South America | 2017 | W.2 | 2a |
| MG264077 | EC/08/2017 | Canis familiaris | Ecuador | South America | 2017 | W.2 | 2c |
| MG264078 | EC/24/2017 | Canis familiaris | Ecuador | South America | 2017 | W.2 | 2b |
| MH476581 | Canine/China/12/2017 | Canis familiaris | China | Asia | 2017 | W.2 | 2c |
| MH476583 | Canine/China/14/2017 | Canis familiaris | China | Asia | 2017 | W.2 | 2c |
| MH476584 | Canine/China/15/2017 | Canis familiaris | China | Asia | 2017 | W.2 | 2c |
| MH476585 | Canine/China/16/2017 | Canis familiaris | China | Asia | 2017 | W.2 | 2c |
| MH476587 | Canine/China/18/2017 | Canis familiaris | China | Asia | 2017 | W.2 | 2c |
| MH476592 | Canine/China/23/2017 | Canis familiaris | China | Asia | 2017 | W.2 | 2c |
| MH660909 | 5 MGL | Canis familiaris | Mongolia | Asia | 2017 | W.2 | 2c |
| MH711894 | CU24 | Canis familiaris | Thailand | Asia | 2016 | W.2 | 2c |
| MH711902 | CU21 | Felis catus | China | Asia | 2016 | W.2 | 2c |
| MK344433 | SV17/17 | Canis familiaris | Brazil | South America | 2017 | W.2 | 2a |
| MK344434 | SV18/16 | Canis familiaris | Brazil | South America | 2016 | W.2 | 2c |
| MK344436 | SV32/16 | Canis familiaris | Brazil | South America | 2016 | W.2 | 2c |
| MK344438 | SV129/16 | Canis familiaris | Brazil | South America | 2016 | W.2 | 2c |
| MK344446 | SV181/17 | Canis familiaris | Brazil | South America | 2017 | W.2 | 2c |
| MK344448 | SV186/17 | Canis familiaris | Brazil | South America | 2017 | W.2 | 2c |
| MK344450 | SV190/17 | Canis familiaris | Brazil | South America | 2017 | W.2 | 2b |
| MK344452 | SV229/16 | Canis familiaris | Brazil | South America | 2016 | W.2 | 2c |
| MK344453 | SV239/15 | Canis familiaris | Brazil | South America | 2015 | W.2 | 2c |
| MK344454 | SV240/15 | Canis familiaris | Brazil | South America | 2015 | W.2 | 2c |
| MK344455 | SV242/15 | Canis familiaris | Brazil | South America | 2015 | W.2 | 2c |
| MK344456 | SV243/15 | Canis familiaris | Brazil | South America | 2015 | W.2 | 2c |
| MK344457 | SV251/15 | Canis familiaris | Brazil | South America | 2015 | W.2 | 2c |
| MK344458 | SV253/15 | Canis familiaris | Brazil | South America | 2015 | W.2 | 2c |
| MK344459 | SV301/17 | Canis familiaris | Brazil | South America | 2017 | W.2 | 2c |
| MK344460 | SV406/15 | Canis familiaris | Brazil | South America | 2015 | W.2 | 2c |
| MK344461 | SV407/15 | Canis familiaris | Brazil | South America | 2015 | W.2 | 2c |

| | | | | | | | |
|---|---|---|---|---|---|---|---|
| MK344462 | SV600/15 | Canis familiaris | Brazil | South America | 2015 | W.2 | 2c |
| MK344463 | SV601/15 | Canis familiaris | Brazil | South America | 2015 | W.2 | 2c |
| MK344464 | SV615/15 | Canis familiaris | Brazil | South America | 2015 | W.2 | 2c |
| MK344466 | SV638/15 | Canis familiaris | Brazil | South America | 2015 | W.2 | 2c |
| MK344467 | SV678/15 | Canis familiaris | Brazil | South America | 2015 | W.2 | 2c |
| MK344470 | SV731/15 | Canis familiaris | Brazil | South America | 2015 | W.2 | 2b |
| MK388674 | HB2017 | Canis familiaris | China | Asia | 2017 | W.2 | 2c |
| MK413743 | CPV-2c_PA15423/16 | Felis catus | Italy | Europe | 2016 | W.2 | 2c |
| MK413744 | CPV-2c_PA36395/16 | Canis familiaris | Italy | Europe | 2016 | W.2 | 2c |
| MK413746 | CPV-2c_PA41113c2/16 | Canis familiaris | Italy | Europe | 2016 | W.2 | 2c |
| MK413747 | CPV-2c_PA45984/16 | Canis familiaris | Italy | Europe | 2016 | W.2 | 2c |
| MK413748 | CPV-2c_CT1839id0018/17 | Canis familiaris | Italy | Europe | 2017 | W.2 | 2c |
| MK413749 | CPV-2c_CT1839id2213/17 | Canis familiaris | Italy | Europe | 2017 | W.2 | 2c |
| MK413750 | CPV-2c_PA27184/17 | Canis familiaris | Italy | Europe | 2017 | W.2 | 2c |
| MK806279 | IZSSI_PA24478/18_id3184 | Canis familiaris | Italy | Europe | 2018 | W.2 | 2c |
| MK806280 | IZSSI_PA24478/18_id3230 | Canis familiaris | Italy | Europe | 2018 | W.2 | 2c |
| MK806281 | IZSSI_PA31342/18 | Canis familiaris | Italy | Europe | 2018 | W.2 | 2c |
| MK806282 | IZSSI_PA5455/19 | Canis familiaris | Italy | Europe | 2018 | W.2 | 2c |
| MK806283 | IZSSI_PA5446/19 | Canis familiaris | Italy | Europe | 2018 | W.2 | 2c |
| MK806284 | IZSSI_RG3408/19 | Canis familiaris | Italy | Europe | 2019 | W.2 | 2c |
| MK806285 | IZSSI_PA5632/19 | Canis familiaris | Italy | Europe | 2019 | W.2 | 2c |
| MK895486 | IZSSI_PA1464/19_idYV2 | Canis familiaris | Nigeria | Africa | 2018 | W.2 | 2c |
| MK895487 | IZSSI_PA1464/19_idEV8 | Canis familiaris | Nigeria | Africa | 2018 | W.2 | 2c |
| MK895488 | IZSSI_PA1464/19_idJOE2 | Canis familiaris | Nigeria | Africa | 2018 | W.2 | 2c |
| MK895489 | IZSSI_PA1464/19_idNC | Canis familiaris | Nigeria | Africa | 2018 | W.2 | 2c |
| MK895490 | IZSSI_PA1464/19_idPSV21 | Canis familiaris | Nigeria | Africa | 2018 | W.2 | 2c |
| MN451675 | CPV601 | Canis familiaris | USA | North America | 2016 | W.2 | 2c |
| MN451676 | CPV603 | Canis familiaris | USA | North America | 2017 | W.2 | 2c |
| MN451677 | CPV604 | Canis familiaris | USA | North America | 2007 | W.2 | 2b |
| MN451678 | CPV605 | Canis familiaris | Nigeria | Africa | 2018 | W.2 | 2c |
| MN451679 | CPV606 | Canis familiaris | USA | North America | 2014 | W.2 | 2c |
| MN451680 | CPV607 | Canis familiaris | Nigeria | Africa | 2018 | W.2 | 2c |
| MN451681 | CPV608 | Canis familiaris | Nigeria | Africa | 2018 | W.2 | 2c |

| | | | | | | | |
|---|---|---|---|---|---|---|---|
| MN451682 | CPV609 | Canis familiaris | Nigeria | Africa | 2018 | W.2 | 2c |
| MN451684 | CPV611 | Canis familiaris | USA | North America | 2019 | W.2 | 2c |
| MN451685 | CPV612 | Canis familiaris | USA | North America | 2019 | W.2 | 2c |
| MN451686 | CPV613 | Canis familiaris | USA | North America | 2019 | W.2 | 2c |
| MN451690 | CPV617 | Canis familiaris | USA | North America | 2011 | W.2 | 2c |
| MN832850 | Taiwan/2018 | Pangolin | Taiwan | Asia | 2018 | W.2 | 2c |
| MT010564 | CPV-AHhf1 | Canis familiaris | China | Asia | 2018 | W.2 | 2c |
| MW648368 | 30-UCS | Canis familiaris | Brazil | South America | 2020 | W.2 | 2a |
| MW648369 | 31-UCS | Canis familiaris | Brazil | South America | 2020 | W.2 | 2a |
| AY742953 | CPV-435 | Canis familiaris | USA | North America | 2003 | W.3 | 2a |
| DQ340411 | BR8-90 | Canis familiaris | Brazil | South America | 1990 | W.3 | 2a |
| DQ340412 | BR17-90 | Canis familiaris | Brazil | South America | 1990 | W.3 | 2a |
| DQ340413 | BR18-90 | Canis familiaris | Brazil | South America | 1990 | W.3 | 2a |
| DQ340414 | BR31-90 | Canis familiaris | Brazil | South America | 1990 | W.3 | 2a |
| DQ340415 | BR43-91 | Canis familiaris | Brazil | South America | 1991 | W.3 | 2a |
| DQ340416 | BR47-91 | Canis familiaris | Brazil | South America | 1991 | W.3 | 2a |
| DQ340417 | BR52-91 | Canis familiaris | Brazil | South America | 1991 | W.3 | 2a |
| DQ340418 | BR491-92 | Canis familiaris | Brazil | South America | 1992 | W.3 | 2a |
| DQ340419 | BR570-92 | Canis familiaris | Brazil | South America | 1992 | W.3 | 2a |
| DQ340420 | BR593-92 | Canis familiaris | Brazil | South America | 1992 | W.3 | 2a |
| DQ340421 | BR597-92 | Canis familiaris | Brazil | South America | 1992 | W.3 | 2a |
| DQ340422 | BR22-93 | Canis familiaris | Brazil | South America | 1993 | W.3 | 2a |
| DQ340423 | BR136-93 | Canis familiaris | Brazil | South America | 1993 | W.3 | 2a |
| DQ340424 | BR137-93 | Canis familiaris | Brazil | South America | 1993 | W.3 | 2a |
| DQ340425 | BR227-93 | Canis familiaris | Brazil | South America | 1993 | W.3 | 2a |
| DQ340426 | BR84-94 | Canis familiaris | Brazil | South America | 1994 | W.3 | 2a |
| DQ340427 | BR133-94 | Canis familiaris | Brazil | South America | 1994 | W.3 | 2a |
| DQ340428 | BR209-94 | Canis familiaris | Brazil | South America | 1994 | W.3 | 2a |
| DQ340429 | BR237-94 | Canis familiaris | Brazil | South America | 1994 | W.3 | 2a |
| DQ340431 | BR56-95 | Canis familiaris | Brazil | South America | 1995 | W.3 | 2a |
| DQ340432 | BR62-95 | Canis familiaris | Brazil | South America | 1995 | W.3 | 2a |
| DQ340433 | BR7168-00 | Canis familiaris | Brazil | South America | 2000 | W.3 | 2a |
| DQ340434 | BR8155-00 | Canis familiaris | Brazil | South America | 2000 | W.3 | 2a |

| | | | | | | | |
|---|---|---|---|---|---|---|---|
| KX774249 | Bel2014-01 | Canis familiaris | Brazil | South America | 2014 | W.3 | 2b |
| KX774250 | Bel2016-01 | Canis familiaris | Brazil | South America | 2016 | W.3 | 2b |
| KX774251 | Bel2015-01 | Canis familiaris | Brazil | South America | 2015 | W.3 | 2b |
| KX774252 | Bel2015-02 | Canis familiaris | Brazil | South America | 2015 | W.3 | 2b |
| MF177251 | BRA01/13 | Canis familiaris | Brazil | South America | 2013 | W.3 | 2b |
| MF177259 | BRA02/13 | Canis familiaris | Brazil | South America | 2013 | W.3 | 2b |
| OR230511 | CPV/UFT02/2022/BRA | Canis familiaris | Brazil | South America | 2022 | W.3 | 2a |
| OR230514 | CPV/UFT05/2022/BRA | Canis familiaris | Brazil | South America | 2022 | W.3 | 2a |
| OR230516 | CPV/UFT07/2023/BRA | Canis familiaris | Brazil | South America | 2023 | W.3 | 2a |
| AB054215 | V120 | Felis Catus | Japan | Asia | 2001 | W.4 | 2a |
| AY742935 | CPV-U6 | Canis familiaris | Germany | Europe | 1995 | W.4 | 2a |
| DQ025943 | 01S1 | Canis familiaris | France | Europe | 2005 | W.4 | 2a |
| DQ025944 | 02B2 | Canis familiaris | France | Europe | 2005 | W.4 | 2a |
| DQ025945 | 02B3 | Canis familiaris | France | Europe | 2005 | W.4 | 2a |
| DQ025947 | 02B5 | Canis familiaris | France | Europe | 2005 | W.4 | 2a |
| DQ025958 | 03C2 | Canis familiaris | France | Europe | 2005 | W.4 | 2a |
| DQ025962 | 03C6 | Canis familiaris | France | Europe | 2005 | W.4 | 2a |
| DQ025982 | 04S13 | Canis familiaris | France | Europe | 2005 | W.4 | 2a |
| DQ025983 | 04S14 | Canis familiaris | France | Europe | 2005 | W.4 | 2a |
| DQ025984 | 04S15 | Canis familiaris | France | Europe | 2005 | W.4 | 2a |
| DQ025986 | 04S17 | Canis familiaris | France | Europe | 2005 | W.4 | 2a |
| DQ026001 | 04S32 | Canis familiaris | France | Europe | 2005 | W.4 | 2a |
| DQ026002 | 04S33 | Canis familiaris | France | Europe | 2005 | W.4 | 2a |
| DQ340430 | BR46-95 | Canis familiaris | Brazil | South America | 1995 | W.4 | 2a |
| FJ005252 | 96/02 | Canis familiaris | Italy | Europe | 2002 | W.4 | 2a |
| FJ005253 | 67/05 | Canis familiaris | Italy | Europe | 2005 | W.4 | 2a |
| FJ005255 | 333/05 | Canis familiaris | Italy | Europe | 2005 | W.4 | 2a |
| FJ005256 | 17/08 | Canis familiaris | Italy | Europe | 2008 | W.4 | 2a |
| FJ005259 | 100/08 | Canis familiaris | Italy | Europe | 2008 | W.4 | 2a |
| GU362932 | cat11/08 | Felis catus | Italy | Europe | 2008 | W.4 | 2a |
| GU362933 | 12/08-A | Canis familiaris | Italy | Europe | 2008 | W.4 | 2a |
| GU362934 | 12/08-B | Canis familiaris | Italy | Europe | 2008 | W.4 | 2a |
| JF346754 | Arg9 | Canis familiaris | Argentina | South America | 2003 | W.4 | 2a |

| | | | | | | | |
|---|---|---|---|---|---|---|---|
| JF414817 | Arg5 | Canis familiaris | Argentina | South America | 2003 | W.4 | 2b |
| JN867610 | CPV/Raccoon/VA/118-A.us.07 | Procyon lotor | USA | North America | 2007 | W.4 | 2a |
| JN867611 | CPV/Raccoon/KY/358-B.us.09 | Procyon lotor | USA | North America | 2009 | W.4 | 2a |
| JN867612 | CPV/Raccoon/TN/351.us.09 | Procyon lotor | USA | North America | 2009 | W.4 | 2a |
| JN867614 | CPV/Raccoon/VA/278-A.us.09 | Procyon lotor | USA | North America | 2009 | W.4 | 2a |
| JN867615 | CPV/Raccoon/GA/287.us.08 | Procyon lotor | USA | North America | 2009 | W.4 | 2a |
| JN867616 | CPV/Raccoon/GA/289.us.08 | Procyon lotor | USA | North America | 2008 | W.4 | 2a |
| JN867617 | CPV/Raccoon/349.us.08 | Procyon lotor | USA | North America | 2008 | W.4 | 2a |
| KF539793 | H-5 | Canis familiaris | Hungary | Europe | 2012 | W.4 | 2a |
| KF539794 | H-7 | Canis familiaris | Hungary | Europe | 2012 | W.4 | 2a |
| KF539796 | H-9 | Canis familiaris | Hungary | Europe | 2012 | W.4 | 2a |
| KF539800 | H-27 | Canis familiaris | Hungary | Europe | 2012 | W.4 | 2a |
| KF539805 | H-36 | Canis familiaris | Hungary | Europe | 2012 | W.4 | 2a |
| KR559891 | PT032/12 | Canis familiaris | Portugal | Europe | 2012 | W.4 | 2a |
| KX434457 | CPV_IZSSI_987_10 | Canis familiaris | Italy | Europe | 2010 | W.4 | 2a |
| MF177224 | 43-97 | Canis familiaris | Italy | Europe | 1997 | W.4 | 2a |
| MF177231 | 260-00 | Canis familiaris | Italy | Europe | 2000 | W.4 | 2b |
| MF177233 | 19-99 | Canis familiaris | Italy | Europe | 1999 | W.4 | 2a |
| MF177246 | Arg50 | Canis familiaris | Argentina | South America | 2009 | W.4 | 2b |
| MF177256 | BRA05/13 | Canis familiaris | Brazil | South America | 2013 | W.4 | 2b |
| MF177258 | BRA03/13 | Canis familiaris | Brazil | South America | 2013 | W.4 | 2b |
| MF177281 | UY6.06 | Canis familiaris | Uruguay | South America | 2006 | W.4 | 2a |
| MG264075 | EC/01/2017 | Canis familiaris | Ecuador | South America | 2017 | W.4 | 2a |
| MK344439 | SV155/16 | Canis familiaris | Brazil | South America | 2016 | W.4 | 2b |
| MK344440 | SV156/16 | Canis familiaris | Brazil | South America | 2016 | W.4 | 2b |
| MK344441 | SV157/16 | Canis familiaris | Brazil | South America | 2016 | W.4 | 2b |
| MK344465 | SV637/15 | Canis familiaris | Brazil | South America | 2015 | W.4 | 2b |
| MK413742 | CPV-2b_PA13600/17 | Canis familiaris | Italy | Europe | 2017 | W.4 | 2b |
| MN451674 | CPV353 | Canis familiaris | USA | North America | 1996 | W.4 | 2a |
| MN451691 | RACCPV1 | Canis familiaris | USA | North America | 2009 | W.4 | 2a |
| OR230510 | CPV/UFT01/2022/BRA | Canis familiaris | Brazil | South America | 2022 | W.4 | 2c |
| OR230512 | CPV/UFT03/2022/BRA | Canis familiaris | Brazil | South America | 2022 | W.4 | 2a |
| OR230513 | CPV/UFT04/2022/BRA | Canis familiaris | Brazil | South America | 2022 | W.4 | 2c |

| | | | | | | | |
|---|---|---|---|---|---|---|---|
| OR230515 | CPV/UFT06/2022/BRA | Canis familiaris | Brazil | South America | 2022 | W.4 | 2a |
