## Supplementary Table S4 for "Recovery of complete genomes of canine parvovirus from clinical samples"

**Table S4**. Details of the 185 complete and near-complete CPV genome sequences sampled from NCBI and used in the Bayesian phylogenetic reconstruction.

| Accession | Year | Country | Accession | Year | Country |
|---|---|---|---|---|---|
| EU659118 | 1981 | USA | KX434459 | 2011 | Italy |
| MN451673 | 1993 | USA | MF177278 | 2011 | Ecuador |
| MN451672 | 1993 | USA | MF177272 | 2011 | Ecuador |
| MF177233 | 1999 | Italy | MF177273 | 2011 | Ecuador |
| MF177226 | 1999 | Italy | MF177270 | 2011 | Ecuador |
| MF177231 | 2000 | Italy | MF177263 | 2011 | Ecuador |
| MF177239 | 2001 | Italy | MF177274 | 2011 | Ecuador |
| MF177240 | 2002 | Italy | MF177264 | 2011 | Ecuador |
| MF177234 | 2003 | Italy | MF177267 | 2011 | Ecuador |
| EF011664 | 2004 | China | MF177279 | 2011 | Ecuador |
| MF177236 | 2004 | Italy | KM457131 | 2011 | Uruguay |
| MF177238 | 2005 | Italy | KM457125 | 2011 | Uruguay |
| MF177281 | 2006 | Uruguay | KM457124 | 2011 | Uruguay |
| MF177235 | 2006 | Italy | KM457127 | 2011 | Uruguay |
| KM457105 | 2006 | Uruguay | KM457130 | 2011 | Uruguay |
| MF177282 | 2006 | Uruguay | KM457129 | 2011 | Uruguay |
| KM457104 | 2006 | Uruguay | KM457142 | 2011 | Uruguay |
| KM457106 | 2006 | Uruguay | KM457128 | 2011 | Uruguay |
| KM457103 | 2006 | Uruguay | MF177286 | 2011 | Uruguay |
| JN867610 | 2007 | USA | MF177260 | 2012 | Brazil |
| MF177237 | 2007 | Italy | KX434460 | 2012 | Italy |
| KM457110 | 2007 | Uruguay | KR002795 | 2013 | China |
| KM457108 | 2007 | Uruguay | KR002797 | 2013 | China |
| KM457107 | 2007 | Uruguay | KR002794 | 2013 | China |
| KM457109 | 2007 | Uruguay | KR002798 | 2013 | China |
| JN867616 | 2008 | USA | MF177256 | 2013 | Brazil |
| JN867617 | 2008 | USA | MF177258 | 2013 | Brazil |
| MF177244 | 2008 | Argentina | MF177259 | 2013 | Brazil |
| MF177242 | 2008 | Argentina | MF177261 | 2013 | Brazil |
| MF177245 | 2008 | Argentina | KR002801 | 2014 | China |
| MF177243 | 2008 | Argentina | MF423125 | 2014 | Canada |
| KM457111 | 2008 | Uruguay | KR002802 | 2014 | China |
| KM457112 | 2008 | Uruguay | KR002804 | 2014 | China |
| KM457113 | 2008 | Uruguay | MF423123 | 2014 | Canada |
| KM457114 | 2008 | Uruguay | MF423124 | 2014 | Canada |
| JN867613 | 2009 | USA | MF177255 | 2014 | Brazil |
| JN867611 | 2009 | USA | MF177257 | 2014 | Brazil |
| KX434454 | 2009 | Italy | MN451679 | 2014 | USA |
| MF177246 | 2009 | Argentina | MH106699 | 2015 | China |
| JN867614 | 2009 | USA | MH106698 | 2015 | China |
| JN867615 | 2009 | USA | KX774252 | 2015 | Brazil |

| Accession | Year | Country | Accession | Year | Country |
|---|---|---|---|---|---|
| MF177262 | 2009 | Paraguay | MF457594 | 2015 | USA |
| MF177283 | 2009 | Uruguay | KY073269 | 2015 | Brazil |
| MF177284 | 2009 | Uruguay | KU508693 | 2015 | Australia |
| KX434455 | 2009 | Italy | KU508692 | 2015 | Australia |
| KU508407 | 2009 | Italy | KU508691 | 2015 | Australia |
| MF177227 | 2009 | France | KX618915 | 2016 | Singapore |
| MF177228 | 2009 | Italy | MH106700 | 2016 | China |
| KM457115 | 2009 | Uruguay | MH711902 | 2016 | Thailand |
| KM457118 | 2009 | Uruguay | MH476588 | 2017 | China |
| KM457117 | 2009 | Uruguay | MH476586 | 2017 | China |
| KM457116 | 2009 | Uruguay | MF134808 | 2017 | China |
| MF069442 | 2010 | Canada | MH476582 | 2017 | China |
| KF638400 | 2010 | China | MG013488 | 2017 | China |
| KM457102 | 2010 | Uruguay | MH476584 | 2017 | China |
| KM457132 | 2010 | Uruguay | MH476587 | 2017 | China |
| KM457134 | 2010 | Uruguay | MH476581 | 2017 | China |
| KM457133 | 2010 | Uruguay | MH476592 | 2017 | China |
| KX434457 | 2010 | Italy | MF510157 | 2017 | Italy |
| MF177252 | 2010 | Brazil | MH476583 | 2017 | China |
| MF177253 | 2010 | Brazil | MN451678 | 2018 | Nigeria |
| MF177248 | 2010 | Argentina | MN451682 | 2018 | Nigeria |
| MF177247 | 2010 | Argentina | MT010564 | 2018 | China |
| MF177249 | 2010 | Argentina | OP093955 | 2019 | Brazil |
| MF177285 | 2010 | Uruguay | OP093952 | 2019 | Brazil |
| MF177230 | 2010 | Italy | OP093953 | 2019 | Brazil |
| MF177250 | 2010 | Brazil | OP093956 | 2019 | Brazil |
| MF177254 | 2010 | Brazil | OP093954 | 2019 | Brazil |
| KM457122 | 2010 | Uruguay | MN451684 | 2019 | USA |
| KM457121 | 2010 | Uruguay | MN451686 | 2019 | USA |
| KM457126 | 2010 | Uruguay | MZ362881 | 2019 | Australia |
| KM457119 | 2010 | Uruguay | MN451685 | 2019 | USA |
| KM457120 | 2010 | Uruguay | MW653251 | 2020 | Iran |
| KM457123 | 2010 | Uruguay | MW539053 | 2020 | Turkey |
| JX660690 | 2011 | China | MW811189 | 2020 | China |
| JQ268283 | 2011 | China | MW811188 | 2020 | China |
| KM457136 | 2011 | Uruguay | OM721656 | 2021 | Turkey |
| KM457143 | 2011 | Uruguay | OM523069 | 2021 | China |
| KM457135 | 2011 | Uruguay | ON733252 | 2021 | Hungary |
| KM457140 | 2011 | Uruguay | OM523076 | 2021 | China |
| KM457137 | 2011 | Uruguay | OM523075 | 2021 | China |
| KM457141 | 2011 | Uruguay | OM523072 | 2021 | China |
| KM457138 | 2011 | Uruguay | OM523071 | 2021 | China |
| MF177280 | 2011 | Ecuador | OM523073 | 2021 | China |
| MF177275 | 2011 | Ecuador | OM523074 | 2021 | China |

| Accession | Year | Country | Accession | Year | Country |
|---|---|---|---|---|---|
| MF177271 | 2011 | Ecuador | OR230510 | 2022 | Brazil |
| MF177266 | 2011 | Ecuador | OR230511 | 2022 | Brazil |
| MF177276 | 2011 | Ecuador | OR230512 | 2022 | Brazil |
| MF177277 | 2011 | Ecuador | OR230513 | 2022 | Brazil |
| MF177269 | 2011 | Ecuador | OR230514 | 2022 | Brazil |
| MF177268 | 2011 | Ecuador | OR230515 | 2022 | Brazil |
| MF177265 | 2011 | Ecuador | OR230516 | 2023 | Brazil |
| MN451690 | 2011 | USA | | | |
