## Supplementary Table S5 for "Recovery of complete genomes of canine parvovirus from clinical samples"

**Table S5**: Model comparison of strict molecular clock, uncorrelated relaxed clock and demographic growth models through path sampling (PS) and stepping stone (SS) methods. Bold numbers indicate the best fitting model.

| | Uncorrelated relaxed clock | | Strict clock | |
|---|---|---|---|---|
| Demographic growth model | PS | SS | PS | SS |
| Bayesian SkyGrid | -13376.61 | -13382.40 | -13429.85 | -13432.36 |
| Bayesian Skyline | **-13340.16** | **-13345.81** | -13392.48 | -13397.19 |
